# Perforin-mediated pore formation at the lytic synapse triggers the canonical pyroptotic cell death pathway

**DOI:** 10.1101/2024.05.24.595698

**Authors:** Brienne McKenzie, Marie-Pierre Puissegur, Lucie Demeersseman, Frecia Rodriguez, Emma Cini, Ashwin Jainarayanan, Marcus A. Widdess, Claire C. Staton, Lina Chen, Vincent Sibaud, Laurence Lamant, Michael Dustin, Agathe Bernand, Sabina Müller, Salvatore Valitutti

**Author notes:** These authors contributed equally to this manuscript (Brienne McKenzie, Marie-Pierre Puissegur).

## Abstract

Prokaryotic pore-forming toxins drive inflammasome activation and pyroptosis through K^+^-dependent activation of the canonical NLRP3/caspase-1/gasdermin D signaling axis. In this study, we hypothesized that perforin, a eukaryotic pore-forming protein released into the lytic synapse by antigen-specific cytotoxic T lymphocytes (**CTLs**) upon cognate antigen recognition, mimics the pro-pyroptotic activity of ancestral pore-forming toxins, complementing its role as a conduit for granzymes. Utilizing imaging and molecular approaches, we demonstrate that perforation of target cells upon CTL attack elicits swift K^+^ efflux followed by NLRP3-dependent activation of proinflammatory caspase-1 and its major substrate, the pyroptotic executioner gasdermin D (**GSDMD**). Acute target cell death upon CTL attack is gasdermin-dependent and demonstrates morphological and molecular features of pyroptosis, including pyroptotic body formation, cell bloating, plasma membrane rupture, and release of intracellular contents. Perforation of target cells by soluble perforin is sufficient to trigger rapid K^+^ efflux, caspase-1 activation, and pyroptosis. By contrast, sustained interaction with CTLs unmasks a delayed apoptotic phenotype in the remaining target cells. Interestingly, exposure of target cells to exogenous supramolecular attack particles (**SMAPs**) recapitulates this apoptotic phenotype, suggesting that soluble perforin and SMAPs play dichotomous roles in target cell death. Our results reveal a novel mechanism for engagement of pyroptotic machinery upon CTL attack, in which perforin itself can autonomously engage programmed cell death (**PCD**), highlighting the complexity and diversity of the CTL lytic arsenal.

## INTRODUCTION

CD8^+^ cytotoxic T lymphocytes (**CTLs**) are potent, sensitive effector cells of the adaptive immune system, which eliminate target cells upon recognition of cognate antigen using a variety of cytotoxic strategies, including exocytosis of soluble cytotoxic molecules, death receptor ligation, and release of particulate killing entities known as supra-molecular attack particles (**SMAPs**)^1–3^.

SMAPs are characterized by a glycoprotein shell within which are encapsulated a variety of different molecules, including cytotoxic granzymes and a variety of chemokines^4^. Isolated SMAPs have been shown to kill target cells over a timeframe of hours to days^4,5^, but their precise mechanism of killing remains elusive.

By contrast, the perforin/granzyme pathway is well-characterized, ultra-rapid pathway that involves directional secretion of soluble cytotoxic molecules within specialized regions at the CTL-target cell interface termed lytic synapses (**LS**). Within seconds of lytic granule release, perforation of the target cell membrane by the pore-forming protein perforin occurs; this facilitates delivery of soluble monomeric granzymes, cytotoxic proteases that cleave intracellular substrates and engage the target cell’s caspase network to promote programmed cell death (**PCD**)^6^. When the target cell’s plasma membrane is perforated, Ca^2+^ influx into the target cell can activate ultra-rapid Ca^2+^-dependent reparative mechanisms at the LS (including lysosomal exposure and ESCRT-mediated membrane repair)^7–10^ that dampen CTL-mediated cytotoxicity by promoting membrane repair and perforin degradation. Whether an individual target cell ultimately succumbs to CTL attack depends upon the accumulation of intracellular damage events inflicted by attacking CTLs, weighed against the ultimately finite reparative resources of the target^11,12^. Thus, the LS provides the spatiotemporal context within which different pro-survival and pro-death processes interact and compete to determine target cell fate.

Within this context, perforin plays a particularly crucial role in ultra-rapid lethal hit delivery, undergoing a transformation (at neutral pH and in the presence of Ca^2+^) from soluble secreted monomers to complex oligomeric transmembrane pores that breach the target cell plasma membrane and provide the physical conduit for trafficking of soluble cytotoxic granzymes into the cytosol^6,13–15^. However, this well-studied function may overshadow additional biologically significant functions that emerge independently of granzyme entry as a result of perforin’s pore-forming properties. Perforin is a member of the diverse and ancient class of pore-forming proteins (**PFPs**), and more specifically the membrane attack complex perforin/cholesterol-dependent cytolysin (**MACPF/CDC**) super-family, which includes eukaryotic proteins such as the membrane attack complex (**MAC**) of the complement cascade, as well as prokaryotic pore-forming virulence factors such as streptolycin O (**SLO**) and listeriolysin O (**LLO**)^14,16,17^. Perforation of the eukaryotic plasma membrane by PFPs has an array of intracellular consequences, including the perturbation of local osmotic and electrochemical gradients (particularly K^+^ and Ca^2+^ homeostasis) as a result of compromised plasma membrane integrity and directional ion flux through transmembrane pores^18^. This ion flux activates a multitude of downstream signaling cascades, including cell stress responses, membrane repair, inflammation, and in some cases, PCD^18,19^.

One particularly important consequence of plasma membrane perforation by PFPs is the activation of one or more K^+^-sensitive inflammasomes (e.g. NLRP3^20–22^), multi-protein complexes that mediate the proximity-induced auto-proteolysis of inflammatory caspases such as caspase-1^22–27^. In addition to mediating cytokine secretion in certain cells, canonical inflammasome activation can also trigger the activation of gasdermin D (**GSDMD**), a member of the gasdermin family of pore-forming executioner proteins responsible for driving **pyroptosis** (“fiery death”), a lytic and proinflammatory PCD modality^28–31^. Upon activation, cleaved GSDMD translocates to the plasma membrane, where it oligomerizes and forms stable pores^32,33^, which are impermeable to large molecules such as lactate dehydrogenase (**LDH)**^34^ but selectively permeable to positively charged ions and small non-acidic proteins (e.g. mature interleukin (**IL**)-1β)^35–38^. In cells with highly functional membrane repair mechanisms, GSDMD pores can be removed, rendering pyroptosis reversible at very early stages^39–42^. Otherwise, loss of membrane integrity leads to a collapse in electrochemical gradients, driving a rapid influx of water through GSDMD pores^34,43^. This causes a characteristic “necrotic” cell death morphology^44^, characterized by cell bloating/swelling and formation of pyroptotic bodies, bubbling membrane protrusions that swell and contract stochastically in response to changing osmotic conditions^37,45,46^. Late-stage pyroptosis often involves catastrophic plasma membrane rupture (**PMR**), which releases proinflammatory alarmins (e.g. high mobility group box 1, **HMGB1**) and large cytoplasmic molecules (e.g. LDH) into the microenvironment. Although multiple pyroptotic pathways have since been identified (reviewed in ^28,37,47^), this “canonical” NLRP3/caspase-1/GSDMD pathway remains a widely recognized and crucial pillar of innate immunity.

Within the context of CTL attack, apoptosis has been classically considered the main mediator of CTL-mediated target cell death, yet evidence for inflammatory, immunogenic and non-apoptotic cell death modalities upon CTL attack has also been gradually accumulating in the literature^48–52^. Most recently, several studies have provided direct evidence to support granzyme-mediated pyroptosis upon T cell attack^53–55^. These findings have prompted the re-examination of CTL-mediated killing through the lens of a more nuanced contemporary understanding of non-apoptotic PCD modalities^56^, and highlighted how different cytotoxic molecules may each have their own unique manner of engaging the target cell’s PCD pathways.

In this study, we hypothesize that pore formation by perforin can activate the same inflammatory pro-pyroptotic pathways associated with ancestral pore-forming toxins, independently of cytotoxic granzymes. We utilized imaging and molecular approaches to test the hypothesis that perforin-mediated pore formation upon attack by human antigen-specific CTLs activates the canonical pyroptotic signaling pathway in transformed target cells. We provide evidence that the pyroptosis-initiating signal originates at the LS, whereupon perforin-mediated pore formation leads to rapid K^+^ efflux and activation of the canonical NLRP3/caspase-1/GSDMD pathway. Conversely, we show that sustained interaction with CTLs unmasks a delayed apoptotic phenotype in the remaining target cells. Exposure of target cells to exogenous SMAPs preferentially triggered apoptosis indicating that perforin and SMAPs activate parallel death pathways, with perforin triggering pyroptosis and SMAPs steering the target towards apoptosis. Collectively these results provide evidence for a proinflammatory perforin-dependent caspase-1/GSDMD pyroptotic signaling cascade upon target perforation that contrasts with SMAP-induced apoptotic cell death revealing novel mechanisms for engagement of pyroptotic/apoptotic machinery upon CTL attack, and highlighting the complexity, diversity, and versatility of the CTL lytic arsenal.

## RESULTS

### Potassium efflux is observed in target cells upon perforation by CTLs

To investigate whether a canonical pyroptotic signaling cascade is induced downstream of target cell perforation, we first utilized a multi-cell system wherein clonal antigen-specific human CTLs were conjugated with JY cells (an Epstein Barr virus (**EBV**)-transformed human B cell line used as a conventional target for human CTLs^57^), which had been pulsed or not with 10μM cognate antigenic peptide (peptide from human cytomegalovirus protein pp65); this well-characterized system enabled us to focus upon the target cell response to CTL attack, in the context of strong antigenic stimulation and optimal CTL activation as previously described^7,8,58,59^. JY cells are known to be highly sensitive to CTL attack^7,8,58^, providing a suitable model system for investigating CTL-induced target cell death.

Since the insertion of pore-forming toxins (such as aerolysin^23^, *C. perfringens* β-toxin^60^, *S. aureus-*derived PVL^26^, *T. pyogenes*-derived pyolysin^27^) into the plasma membrane are known to drive rapid K^+^ efflux (reviewed in ^18^), we hypothesized that perforin pore formation may likewise trigger disruption of intracellular K^+^ homeostasis in target cells upon CTL attack, thus providing an activating signal to NLRP3, the prototypic K^+^-sensitive canonical inflammasome^20–22^. Although perforin pores are known to be selectively permeable to cationic cargo^15^, direct assessment of K^+^ loss upon target perforation by human CTLs is largely absent from the literature. To address this question, we made use of the well-described cell membrane-permeable fluorescent K^+^ indicator, Ion Potassium Green-2 AM (IPG-2 AM; formerly Asante Potassium Green-AM, APG-AM^61^), which has been widely used to assess intracellular K^+^ concentration (**[K^+^]*i***) using both imaging and flow cytometry approaches^62–64^. Prior to assessing K^+^ in target cells upon CTL attack, we first validated our flow cytometry approach using nigericin, a well-characterized bacterially-derived K^+^ ionophore whose primary mechanism of action is the reduction of [K^+^]***i*** through K^+^/H^+^ exchange^65–67^, and which has been previously validated using APG-AM probes^63^. As shown in **Supplementary Figure 1a-c**, IPG-2 AM intracellular fluorescent signal decreased in a time-dependent manner upon exposure of JY cells to nigericin, confirming that a reduction in [K^+^]***i*** is indeed detectable in JY cells using this flow cytometry approach.

To assess changes in intracellular K^+^ upon CTL attack, non-pulsed and cognate antigen-pulsed target cells were loaded with IPG-2 AM, conjugated with CTLs for 5 or 15min at various effector:target (**E:T**) ratios, and subjected to analysis by flow cytometry in the presence of a viability dye. As shown in **Figure 1a-d** , a rapid and E:T ratio-dependent loss of intracellular K^+^ was observed in antigen-pulsed target cells upon 5 and 15min conjugation with CTLs; the majority of K^+^ efflux occurred rapidly upon CTL attack (ie within the first 5min of conjugation, **Figure 1a,c**). Dead cells were excluded from this analysis, supporting the interpretation that the IPG-2 AM^dim^ population identified in **Figure 1a-d** contained viable cells with diminished [K^+^]i rather than simply dead cells.

**Figure 1.**
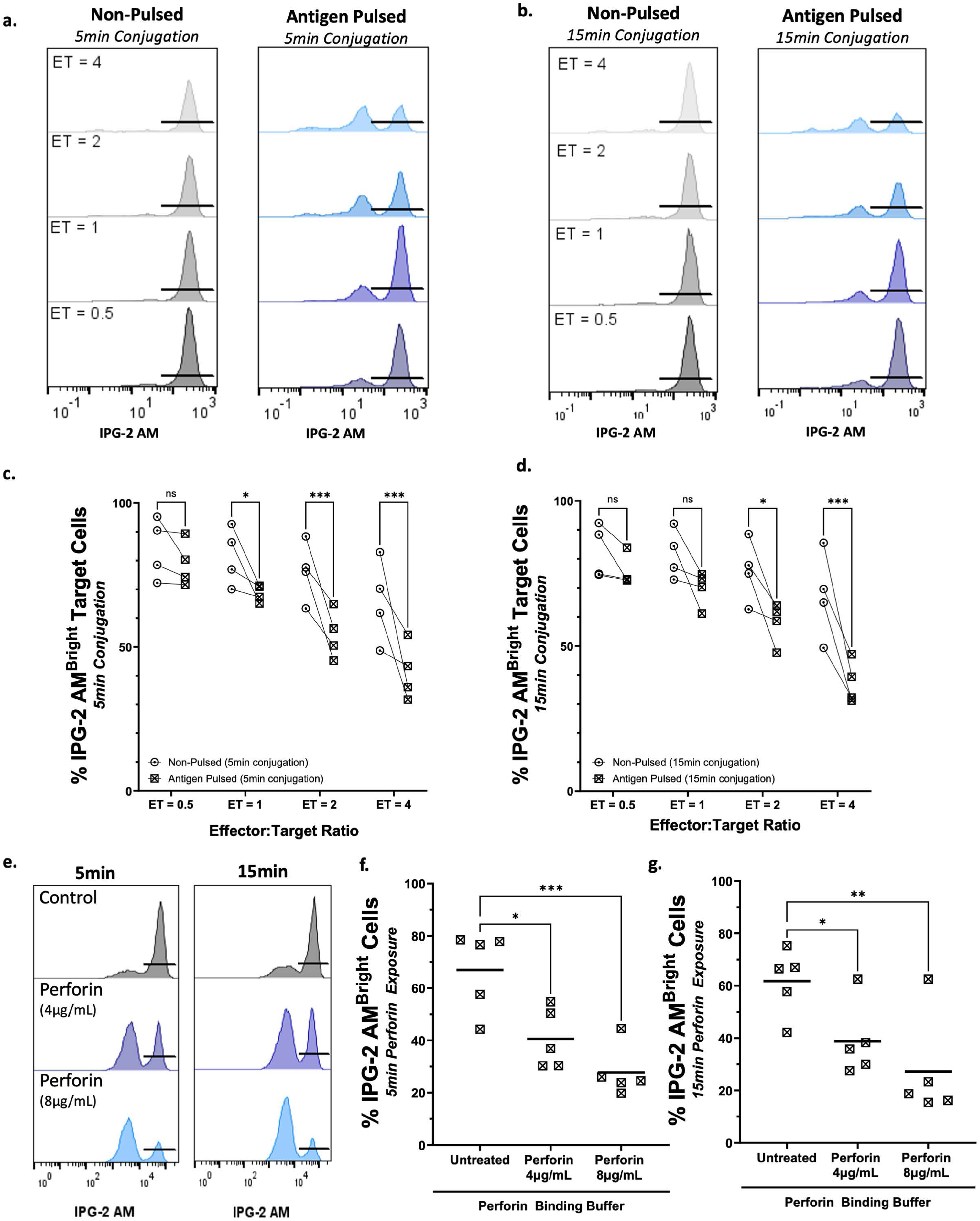
Potassium efflux from target cells occurs upon CTL attack and exposure to soluble perforin **a,b.** JY target cells were pulsed or not with 10μM antigenic peptide, stained with the fluorescent potassium indicator Ion Potassium Green (**IPG-2 AM**), conjugated with antigen-specific CTLs at the indicated E:T ratios for 5 or 15 min, and assessed using flow cytometry. Representative histograms are shown, with the IPG-2 AM^Bright^ population indicated by black bars. **c,d.** Quantification of the percentage of target cells in the IPG-2 AM^Bright^ gate after 5min (**c.**) and 15min (**d.**) conjugation. Each datapoint represents the average of two technical replicates from an individual experiment, with n=4 independent experiments represented on the graph. Two-way ANOVA with Sidak’s multiple comparison test was utilized to assess statistical significance. **e.** JY target cells were preloaded with IPG-2 AM, treated with either solvent control or the indicated concentrations of recombinant human perforin in calcium-enriched perforin binding buffer for 5 or 15min, and assessed by flow cytometry. Representative histograms are shown, with the IPG-2 AM^Bright^ population indicated by black bars. **f,g.** Quantification of the percentage of IPG-2 AM^Bright^ target cells after 5min (**f.**) and 15min (**g.**). Each datapoint represents the average of two technical replicates from an individual experiment, with n=5 independent experiments represented on the graph. Horizontal black lines represent the mean of independent experiments. One-way ANOVA with Sidak’s test for multiple comparisons was utilized to assess statistical differences. \**p*<0.05, ** *p*<0.01, *** *p*<0.001, ns = not significant.

### Soluble human perforin is sufficient to initiate potassium efflux

We next sought to assess whether soluble perforin alone was sufficient to initiate K^+^ efflux from perforated target cells in the absence of cytotoxic granzymes. A rapid and dose-dependent loss of intracellular K^+^ was indeed observed in JY cells upon exposure to soluble perforin, at 5 and 15min, based on the reduction of the percentage of IPG-2 AM^bright^ cells (**Figure 1e-g**). To illustrate that this reduced signal was specific to loss of K^+^ and not merely leakage of IPG-2 AM out of perforated cells, a parallel set of experiments was conducted in which soluble perforin was applied to JY cells in the presence of excess extracellular K^+^, conditions under which the flow of K^+^ ions out of the perforated cell is inhibited^20,22,68^. As shown in **Supplementary Figure 2a**, performing the experiments in high [K^+^] buffer rescued perforin-induced IPG-2 AM signal reduction, supporting the interpretation that loss of IPG-2 AM signal in perforin-exposed cells was indeed attributable specifically to K^+^ efflux.

### Caspase-1 activity is detectable in target cells upon CTL attack

We next sought to test the hypothesis that caspase-1, the canonical inflammatory caspase, was activated in target cells upon CTL attack downstream of K^+^ efflux. To accomplish this, we first verified full-length caspase-1 expression at baseline in JY cells, using the triple-negative breast cancer line MDA-MB231 as a positive control^69,70^ (**Supplementary Figure 3a**). To test whether caspase-1 became activated specifically in targets upon CTL attack, we utilized a well-validated cell-permeable activity-dependent fluorescent probe that irreversibly binds to active caspase-1 through covalent coupling^71,72^. This highly sensitive approach enabled the quantification of caspase-1 activity at the single-cell level by flow cytometry. To first validate this system, JY cells were exposed to nigericin (described above), a well-characterized and widely utilized activator caspase-1^73,74^, and caspase-1 assessed by flow cytometry. Compared to solvent-treated controls, nigericin exposure resulted in a significant increase in the percentage of cells displaying caspase-1 activity (**Supplementary Figure 3b**). To assess caspase-1 activity in target cells upon CTL attack, CTLs were conjugated with antigen-pulsed or non-pulsed JY cells that had been pre-loaded with the caspase-1 probe; after 5 and 15min conjugations, the conjugates were disrupted and subjected to flow cytometry (gating strategy shown in **Supplementary Figure 3c**). Under these conditions, both the percentage of target cells with high caspase-1 activity (**Figure 2a,b**) and the median caspase-1 fluorescence intensity **(Figure 2a,c)** significantly increased following CTL conjugation, compared to non-pulsed controls. To validate the specificity of this read-out, we transfected JY cells with either non-targeting or *CASP1*-targeting siRNA (siRNA validation shown in **Supplementary Figure 3d**) and demonstrated that caspase-1 activity in antigen-pulsed cells (normalized to non-pulsed controls) was significantly decreased in the *CASP1* siRNA-transfected condition compared to non-targeting siRNA-transfected controls (**Supplementary Figure 3e**).

**Figure 2.**
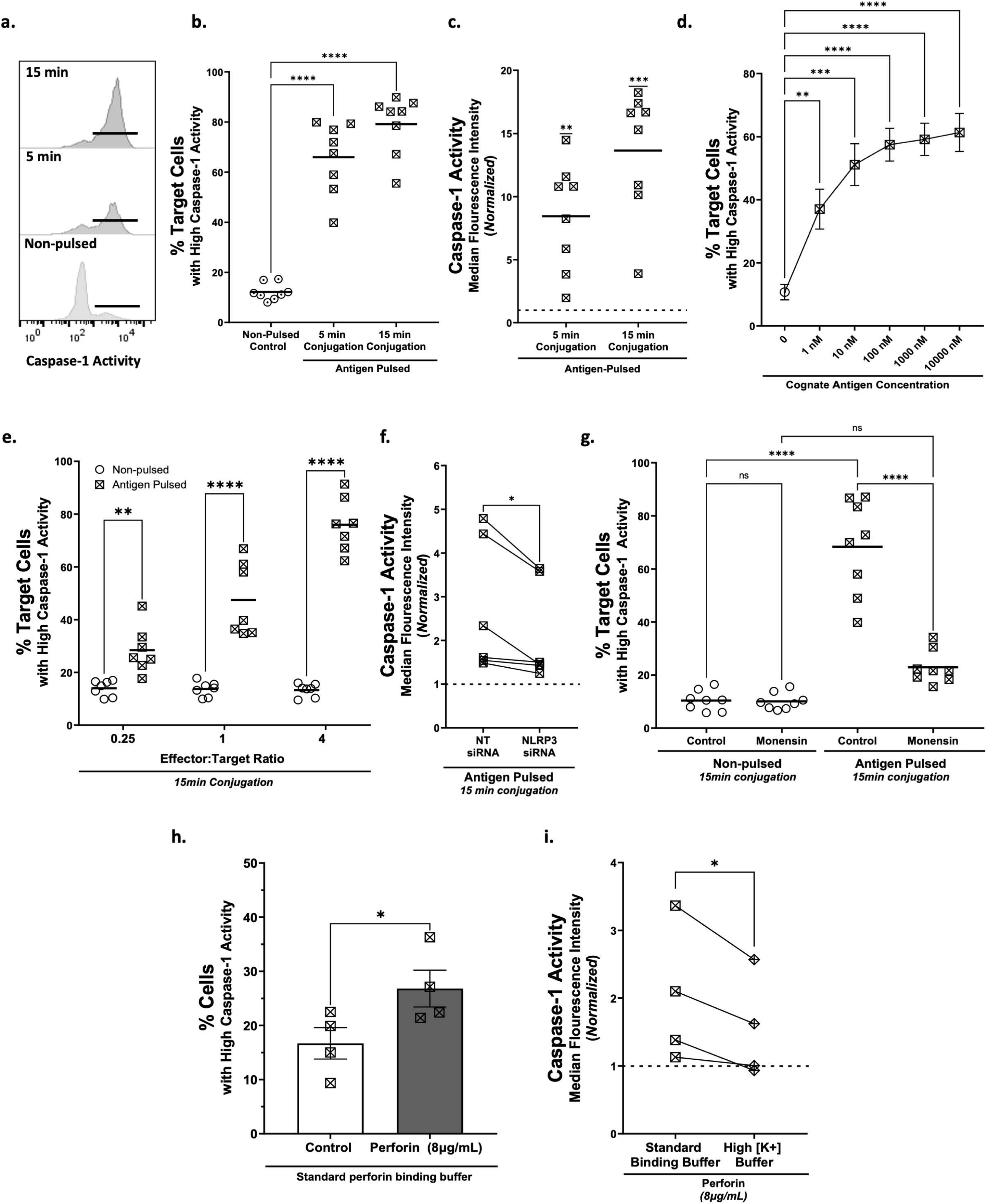
Caspase-1 is activated in JY target cells upon CTL attack and exposure to soluble perforin a-c. JY target cells were pulsed with 10μM antigenic peptide and conjugated with CTLs at an E:T ratio of 4:1 for 5 or 15 min, before being stained with an activity-dependent fluorescent caspase-1 probe and assessed by flow cytometry. Non-pulsed target cells were conjugated with CTLs under identical conditions for 15min as a control. **a.** Representative histograms shown. Black bars represent targets with high caspase-1 activity. **b.** Quantification of the percentage of target cells in the high caspase-1 activity gate. Each datapoint represents the average of two technical replicates from an individual experiment, with a total of n=8 independent experiments represented on the graph. Horizontal black lines represent the mean of independent experiments for each condition. One-way ANOVA with Dunnet’s test for multiple comparisons was utilized to assess statistical significance. **c.** Median caspase-1 fluorescence intensity from the experiments shown in (**b.**), represented as fold increase over non-pulsed control (dotted horizontal line). One-sample *t*-tests were utilized to compare each time point to the hypothetical value of 1.0, representing the non-pulsed control. **d.** JY cells were pulsed with the indicated concentration of antigenic peptide at an E:T ratio of 4:1 for 15min and assessed as in **a-c.** Data shown represent mean +/- SEM, n=5 independent experiments (each with two technical replicates per condition). One-way ANOVA with Dunnet’s test for multiple comparisons was utilized to assess statistical significance. **e.** JY cells were pulsed with 10μM antigenic peptide and conjugated with CTLs at the indicated E:T ratios for 15min and assessed as above. Each datapoint represents the average of two technical replicates from an individual experiment, with n=7 independent experiments represented on the graph. Bars represent the mean of independent experiments. Two-way ANOVA with Sidak’s multiple comparison test was utilized to assess statistical significance. **f.** JY target cells were transfected with non-targeting (**NTsiRNA**) or NLRP3-targeting siRNA (***NLRP3* siRNA**) for 24hr before being pulsed or not with 10μM antigenic peptide and conjugated with CTLs at an E:T ratio of 4:1 for 15 min. JY cells were stained with the fluorescent caspase-1 probe as above. Data shown represent the fold increase in caspase-1 median fluorescence intensity relative to non-pulsed controls (indicated by dotted line) for NT siRNA and NLRP3-targeting siRNA respectively (n=6 biological experiments). A ratio paired *t*-test was utilized to assess statistical differences. **g.** JY cells were pulsed or not with 10μM antigenic peptide and conjugated for 15min at an E:T ratio of 4:1 with CTLs that had been pre-treated with either solvent or monensin (to de-acidify lytic granules) before being stained with the fluorescent caspase-1 probe. Each datapoint represents the average of two technical replicates from an individual experiment, with n=8 independent experiments represented on the graph. Horizontal black lines represent the mean of independent experiments for each condition. Ordinary one-way ANOVA with Tukey’s test for multiple comparisons was utilized to assess statistical significance. **h.** JY cells were pre-loaded with caspase-1 probe in perforin binding buffer, treated with solvent control or 8μg/mL recombinant human perforin for 30min, and assessed by flow cytometry. Each datapoint represents the average of two technical replicates from an individual experiment, with n=4 independent experiments represented on the graph. Bars represent mean +/- SEM. Paired *t*-test was utilized to assess statistical difference. **i.** In parallel, JY cells were pre-loaded with caspase-1 probe in high-potassium perforin binding buffer, treated with 8μg/mL recombinant human perforin for 30min and assessed by flow cytometry. Each datapoint represents the average of two technical replicates from an individual experiment, with n=4 independent experiments represented on the graph. A ratio paired *t*-test was utilized to assess statistical difference.

We next tested whether caspase-1 activity upon CTL attack was antigen-dose dependent. Target cells were pulsed with different concentrations of peptide (from 1nM to 10μm) and conjugated with CTLs for 15 min; a significant increase in the percentage of cells with high caspase-1 activity was observed with as little as 1nM antigenic peptide (**Figure 2d**). We performed a similar analysis to assess the effect of E:T ratio, wherein cells were pulsed with 10μm of antigen and subsequently conjugated with CTLs at an E:T ratio of 0.25, 1, or 4. As shown in **Figure 2e**, the percentage of cells with high caspase-1 activity increased proportionally to E:T ratio, with an E:T ratio as low as 0.25 resulting in a detectable increase in caspase-1 activity after 15min conjugation.

### Caspase-1 activity is diminished upon NLRP3 knockdown

Although various potassium-sensitive inflammasomes can serve as a platform for caspase-1 activity, the NLRP3 inflammasome was of specific interest in this study since it is well-known to be activated by pore-forming toxins in a potassium-dependent manner^22–27^. To test whether NLRP3 was involved in caspase-1 activation upon CTL attack, we first verified NLRP3 protein expression in our target cells of interest (**Supplementary Figure 3a**) before assessing whether NLRP3 deficiency affected caspase-1 activity following CTL attack. We compared the fold increase in caspase-1 fluorescence intensity in targets that had been transfected with either non-coding or *NLRP3*-targeting siRNA (validation shown in **Supplementary Figure 3d**) and subsequently conjugated with CTLs. This analysis revealed a consistent decrease in caspase-1 fluorescence intensity when NLRP3 was knocked down compared to non-targeting controls (**Figure 2f**). The modest nature of this decrease is not unexpected considering that there are alternative inflammasomes that can be engaged under stress conditions to process caspase-1 (e.g. NLRP1, which detects abnormal protease activity and may also be sensitive to K^+^ concentration^68,75^ and NLRC4, which has been shown to be activated alongside NLRP3 in response to bacterial pore forming toxins^24^). Nonetheless, these data suggest a role for the NLRP3 inflammasome in this signaling axis upstream of caspase-1 activation.

### Caspase-1 and caspase-3 activity are often detectable in the same cells upon CTL attack

Caspase-3 is a well-characterized dual-function caspase that can either initiate apoptosis or GSDME-mediated pyroptosis; it is known to be robustly activated upon CTL attack^54,55,76,77^. As such, we next assessed activation of caspase-3 under parallel conditions as caspase-1, utilizing a cell-permeable activity-dependent fluorescent probe. As expected, these experiments confirmed a significant time-dependent and antigen dose-dependent activation of caspase-3 following 5 and 15min conjugation of target cells with CTLs (**Supplementary Figure 4a,b**), which paralleled our observations with caspase-1. As little as 1nM of antigenic stimulation was sufficient to induce significant caspase-3 activity when pulsed cells were conjugated with CTLs (**Supplementary Figure 4b**). We then conducted double-staining experiments to determine whether caspase-1 and caspase-3 activity were detectable in the same cells (**Supplementary Figure 4c**). These experiments revealed a significant caspase-1 single-positive population, along with a double-positive population at both time points tested; caspase-3 single-positive cells remained relatively rare. Collectively, these data suggest that the two caspases are indeed frequently (but not exclusively) activated in the same cells upon CTL attack, and that caspase-1 can be activated prior to (or in the absence of) caspase-3.

### Caspase-1 activity is perforin-dependent and soluble perforin activates caspase-1

We next sought to determine whether caspase-1 activation upon CTL attack required the perforin secretory pathway by pre-treating CTLs with the well-established carboxylic ionophore monensin to increase lytic granule pH, as described previously^7,8,78^. Interfering with the pH of lytic granules has been shown to markedly attenuate perforin-mediated cytotoxicity^7,79^. Herein, we pre-treated CTLs with monensin prior to conjugation with pulsed or non-pulsed targets; as shown in **Figure 2g**, monensin pre-treatment of CTLs largely abrogated target cell caspase-1 activity, supporting the interpretation that the perforin secretory pathway is needed for caspase-1 activation.

We then tested whether target perforation by soluble perforin was sufficient to activate the caspase-1 pathway. As shown in **Figure 2h**, treatment with soluble perforin for 15min significantly increased the percentage of cells with high caspase-1 activity. To assess whether this caspase-1 activity was dependent upon upstream K^+^ efflux as a result of target perforation, we performed the same experiments in the presence of excess extracellular K^+^ to limit K^+^ efflux from perforated targets (as above). When the assays were performed in high [K^+^] perforin binding buffer, caspase-1 fluorescence intensity in perforin-exposed cells was reduced compared to cells in the standard perforin-binding buffer (**Figure 2i**). This supported the hypothesis that caspase-1 activity in perforated target cells was dependent upon K^+^ efflux.

Collectively, these data support a model wherein perforin-mediated pore formation leads to swift K^+^ efflux from perforated cells, leading to activation of the potassium-sensitive NLRP3 inflammasome and the caspase-1 pathway.

### Target cells express multiple gasdermins at the transcript and protein level

Knowing that caspase-1 was activated in target cells upon CTL attack, we next sought to determine whether GSDMD, a canonical executioner of pyroptosis and the quintessential substrate of caspase-1, was likewise activated following CTL attack. GSDMD has been proposed to have a role in anti-tumor immunity^80^ but its activation in the context of CTL attack has never been demonstrated. It is known that gasdermins are expressed to varying degrees in transformed and untransformed cells^28,38^, and to test this we first assessed transcript expression of *GSDMA-E* at baseline in JY cells and a diverse panel of other target cells. As shown in **Supplementary Figure 5a**, *GSDMD* and *GSDME* were the most highly and consistently expressed of the five gasdermins at baseline across a range of target cells. We validated GSDMD expression at the protein level in two well-characterized target cell lines: JY (described above) and D10, a well-characterized melanoma cell line, moderately resistant to CTL attack due to reparative membrane turnover^7,8,81^. MDA-MB231 (previously shown to highly express GSDMD^82^ and GSDME^83^) was used as a positive control. Consistent with transcript data, full-length GSDMD protein was detectable in all three cell lines at baseline (**Supplementary Figure 5b**). Specificity of the GSDMD antibody was validated using siRNA approaches, wherein GSDMD signal was strongly reduced in *GSDMD-*targeting siRNA-transfected but not *GSDME-*targeting siRNA-transfected JY cells (**Supplementary Figure 5c**).

### Cleaved GSDMD accumulates in target cells upon attack by CTLs and is enriched near the immunological synapse

To assess the accumulation of cleaved GSDMD in target cells upon conjugation with CTLs, we profited from two commercially available and widely utilized antibodies^84–87^: the first specifically detects the active N-terminal of GSDMD (“cleaved GSDMD”) but not full-length GSDMD; and the second (also utilized for immunoblots above) detects both full-length and cleaved GSDMD (“total GSDMD”). JY cells were conjugated with CTLs for 15min, fixed, permeabilized, and stained with antibodies against cleaved GSDMD, total GSDMD, and perforin before being subjected to confocal microscopy. As shown in the illustrative 3D reconstruction in **Figure 3a**, total but not cleaved GSDMD immunoreactivity is detectable in non-pulsed target cells interacting with CTLs; however, upon interaction of pulsed JY cells with CTLs and initiation of target cell death, target cells display a high degree of cleaved GSDMD immunoreactivity, frequently associated with the bubbling morphology typical of pyroptosis. To facilitate quantification of cleaved GSDMD immunoreactivity in target cells upon CTL attack, we next focused upon CTL-target cell conjugates where lytic granules were polarized towards the target but the target was not yet excessively bubbling. As shown in **Figure 3b-e**, while total GSDMD immunoreactivity was equivalent in both non-pulsed and pulsed target cells interacting with CTLs (**Figure 3d**), cleaved GSDMD MFI was significantly increased in targets under pulsed conditions after 15min conjugation (**Figure 3e**). JY cells transfected with siRNA targeting *GSDMD* exhibited significantly reduced fluorescent signal from the cleaved GSDMD antibody upon interaction with CTLs compared to non-targeting siRNA-transfected controls (**Supplementary Figure 6a-c**).

**Figure 3.**
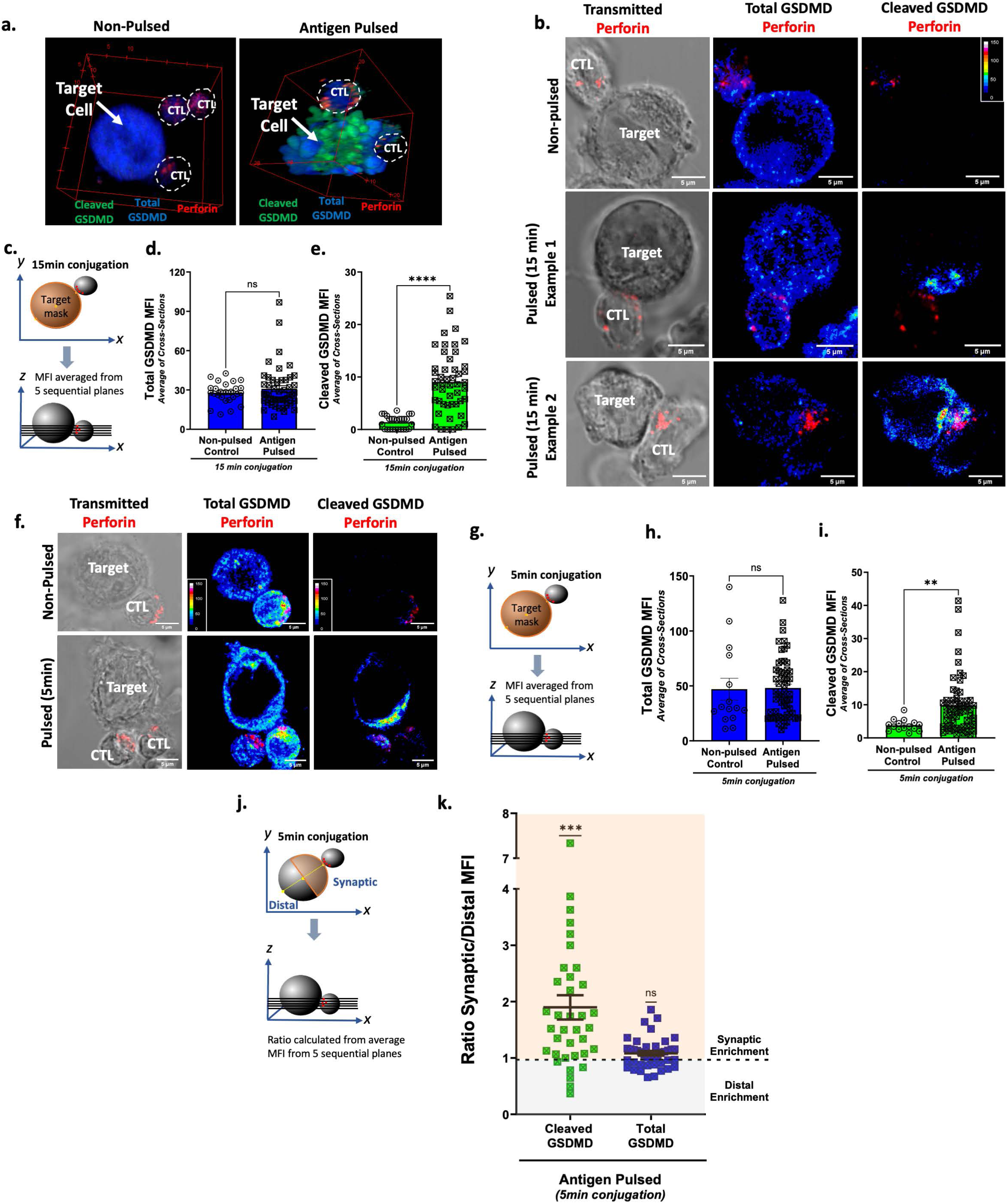
Cleaved GSDMD is detectable upon CTL attack and is enriched adjacent to the synapse a-d. JY target cells were pulsed with 10μM antigenic peptide and conjugated at an E:T ratio of 1:1 with antigen-specific human CTLs for 15min. Conjugates were fixed and subject to immunostaining for total GSDMD, cleaved GSDMD and perforin. **a.** 3D reconstruction with total GSDMD (*blue*), cleaved GSDMD (*green*) and perforin (*red*). **b.** Representative conjugates from non-pulsed and pulsed conditions, with total and cleaved GSDMD shown in pseudocolor and perforin in red. Images shown are from a single plane of the *z*-stack and are illustrative examples from n=4 biological experiments. **c.** Schematic for the quantification of cleaved and total GSDMD mean fluorescence intensity (**MFI**) within the target cell mask across five sequential planes of the *z*-stack. The five planes were chosen to correspond to the height of the lytic synapse in the *z*-dimension and MFI averaged across the five planes. **d-e.** MFI of total **(d.)** and cleaved **(e.)** GSDMD within the target cell mask was measured on 5 sequential planes of the *z*-stack (separated by 1μm) at the height of the lytic synapse and subsequently averaged. Each datapoint corresponds to one target cell, from n=4 biological experiments. Statistical differences between groups were assessed by unpaired *t*-tests. **f-i.** Target cells and CTLs were prepared as above and conjugated for 5min before being subjected to immunostaining and quantification as described above. **g.** Schematic for the quantification of cleaved and total GSDMD MFI across five sequential planes of the *z*-stack. **h-i.** MFI of total **(h.)** and cleaved **(i.)** GSDMD within the target cell was measured on 5 different planes of the *z*-stack (separated by 1μm) at the height of the synpase and subsequently averaged. Each datapoint corresponds to one target cell, from n=4 biological experiments. **j,k.** To assess whether cleaved or total GSDMD was enriched proximal to the synapse, each target cell mask was divided in half in the *x-y* dimension to create “synaptic” and “distal” masks, for which MFI was separately quantified. The ratio of synaptic:distal signal was calculated and averaged across the 5 sequential planes. Ratios higher than 1 indicate synaptic enrichment, lower than 1 indicates distal enrichment, and equal to 1 indicates even distribution throughout the cell. Each datapoint represents the average ratio of synaptic:distal signal across 5 planes for a single cell. Mean ratio of synaptic:distal MFI for both cleaved and total GSDMD were compared to a hypothetical value of 1 (indicating even distribution of signal throughout the cell) using one-sample *t*-tests. Scale bars are 5μm. Error bars represent mean +/- SEM. ** *p*<0.01, *** *p*<0.01, **** *p*<0.0001, ns = non-significant.

Upon visual examination of cleaved GSDMD immunoreactivity in pulsed target cells upon CTL attack (**Figure 3b**), it became apparent that while total GSDMD was evenly distributed throughout the target cell, cleaved GSDMD immunoreactivity appeared concentrated near the LS in a subset of pulsed target cells, suggesting that the initial signal to activate GSDMD may first be triggered locally near the LS. To quantify this observation, we assessed both cleaved and total GSDMD MFI in the synaptic half versus the distal half of pulsed target cells in conjugates with CTLs and generated a ratio of synaptic MFI /distal MFI; in this analysis, a ratio >1 indicated synaptic enrichment of the protein of interest, a ratio = 1 indicated even distribution throughout the cell, and a ratio <1 indicated distal enrichment of the protein of interest. As shown in **Supplementary Figure 7**, after 15min conjugation, total GSDMD was evenly distributed throughout target cells in conjugation with CTLs under non-pulsed and pulsed conditions. However, cleaved GSDMD (which was only consistently detectable under antigen-pulsed conditions) was frequently enriched on the synaptic side of the target cell. To determine whether this phenomenon might occur even more dramatically at earlier timepoints, we performed the same analysis after only 5min conjugation. As shown in **Figure 3f-i**, cleaved GSDMD (but not total GSDMD) was significantly enriched in target cells upon CTL attack even at this early time point. Likewise, cleaved GSDMD was strongly enriched near the LS, while total GSDMD was evenly distributed throughout the cell (**Figure 3j,k**). This observation was conceptually consistent with a localized synaptic trigger for GSDMD activation.

### Target cell death upon attack by human Ag-specific CTL is characterized by membrane bubbling and loss of membrane integrity typical of pyroptosis

Having established that the K^+^/NLRP3/caspase-1/GSDMD canonical pyroptotic pathway was activated in target cells upon CTL attack, we next sought to determine whether target cell death displayed the quintessential molecular and morphological indicators of pyroptosis. To address this, we first utilized time-lapse microscopy to visualize target cell death. Although microscopy alone does not distinguish amongst the various types of programmed necrosis (pyroptosis, necroptosis, etc.), time-lapse microscopy remains a powerful and pragmatic approach for identifying the quintessential morphological features of non-apoptotic PCD modalities (such as swelling, membrane bubbling, plasma membrane rupture, and vital dye uptake)^44^. To visualize cell death morphology, target cells were pulsed (or not) with cognate antigen for 2hr and conjugated with antigen-specific CTLs in the presence of low-concentration (6.6 µg/mL) PI, which served as a vital dye. As expected, non-pulsed target cells displayed transient non-lethal interactions with CTLs (**Supplementary Figure 8a,d)**, while antigen-pulsed JY cells readily formed conjugates with CTLs, and of these, 78% became PI^+^ within 120min (**Supplementary Figure 8b,c**).

Using this system, we sought to observe and quantify the morphological features of target cell death upon CTL attack. In the example shown in **Figure 4a** and **Supplementary Movie 1**, CTL target recognition was indicated by Ca^2+^ flux and polarization of the CTL’s microtubule organizing centre (**MTOC**) towards the LS. Rapidly thereafter, the target displayed irregular bubbling membrane protrusions, which expanded and contracted in the stochastic, erratic manner typical of pyroptotic bodies^37,46^. This bubbling was followed by cessation of cellular motility, bloating of the target, and ultimately explosive PMR, concurrent with instantaneous PI influx, all of which are prototypical morphological signs of pyroptosis and other necrotic cell death modalities^44^. These observations were recapitulated in the second example shown in **Figure 4b** and **Supplementary Movie 2** (wherein PI influx was slightly more gradual), as well as additional examples from three independent experiments (**Supplementary Figure 8c**). Such morphological features closely recapitulated the bubbling, lytic phenotype of pyroptosis exemplified by cells exposed to nigericin^65,66,88^, shown in **Supplementary Figure 9a,d** and **Supplementary Movies 3-5**, but contrasted starkly with the morphology of apoptosis (exemplified by cells treated with the broad-spectrum kinase inhibitor staurosporine^44^). As shown in **Supplementary Figure 9b,d** and **Supplementary Movies 6-9**, staurosporine-treated apoptotic cell death was slow and controlled, with the gradual appearance of small fragmented apoptotic bodies rather than violent swelling, bubbling, or catastrophic PMR.

**Figure 4.**
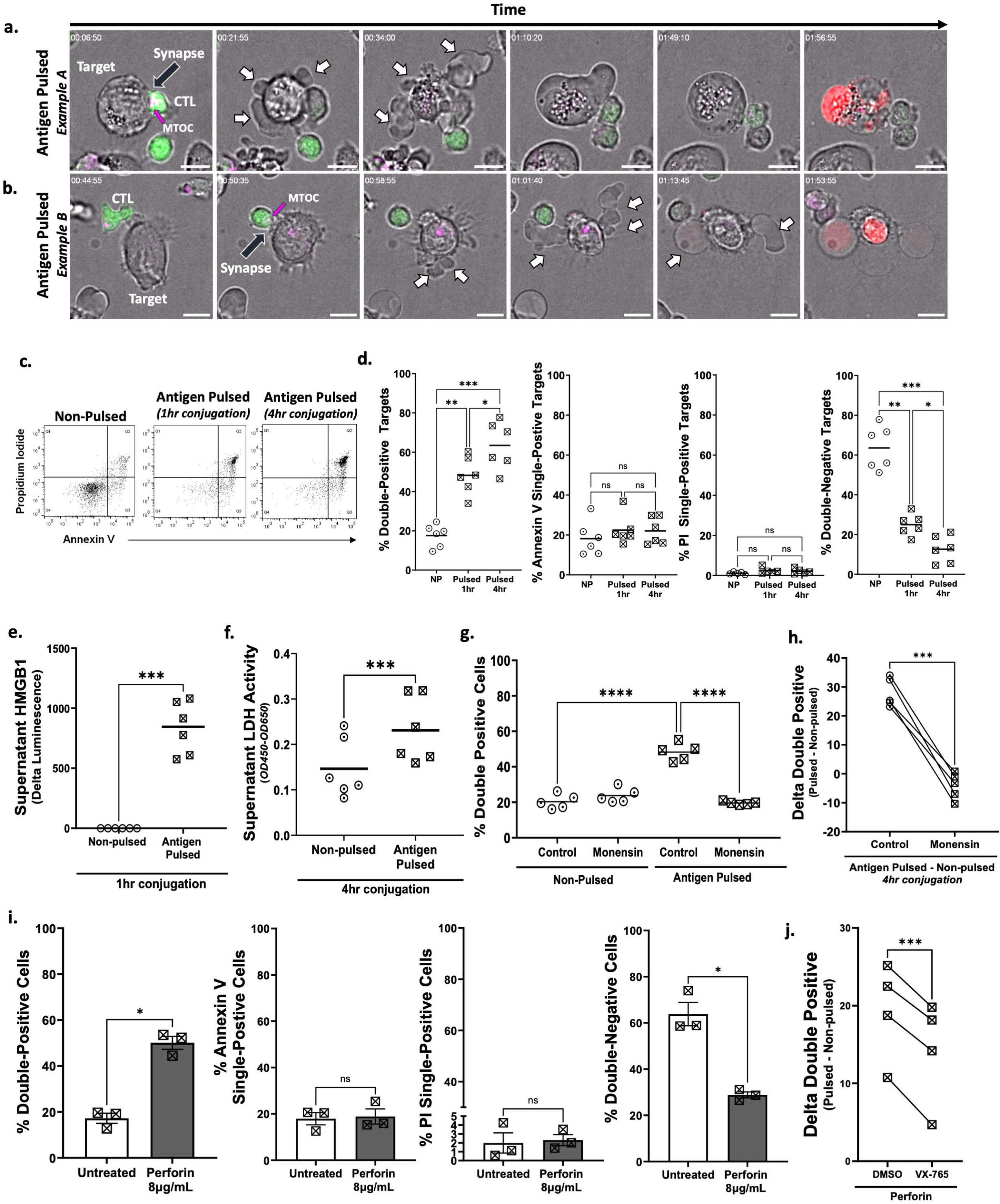
Target cells die by pyroptosis upon CTL attack a,b. JY target cells were pulsed with 10μM antigenic peptide, and conjugated at an E:T ratio of 1:1 with antigen-specific human CTLs pre-loaded with Fluo-8-AM (*green*) and SPY650 Tubulin (*magenta*) in the presence of 6.6µg/mL propidium iodide (*red*). Conjugate formation and cell death morphology was assessed using spinning disk time lapse microscopy for 2hr. Snapshots (from **Supplementary Movie 1** and **2**) illustrate various stages of CTL attack including initial CTL contact, MTOC polarization (*magenta arrows*), synapse formation (*black arrows*), pyroptotic body formation (*white arrows*), and loss of membrane integrity, indicated by PI entry. Two representative examples from n=3 independent experiments are shown; additional examples are shown in **Supplementary** Figure 8. Scale bar is 10μm. **c,d.** JY cells were pulsed or non-pulsed (**NP**) with antigenic peptide and conjugated at an E:T ratio of 4:1 with antigen-specific human CTLs for 1 or 4 hours. Following conjugation, cells were subjected to Annexin V/PI staining and analyzed by flow cytometry. **c.** Representative dotplots of target cells from n=6 biological experiments. **d**. Data shown represent the percentage of target cells in each quadrant (Annexin V/PI double positive, Annexin V single-positive, PI single-positive, or double negative) at the indicated time points. Each datapoint represents the average of two technical replicates from an individual experiment, with n=6 independent experiments represented on the graph. Black bars represent the mean of n=6 biological experiments. Repeated measurement one-way ANOVA with Tukey’s test for multiple comparisons was utilized to assess statistical significance. **e.** Antigen-pulsed or NP JY cells were conjugated at a 4:1 E:T ratio with CTLs for 1hr and supernatants subjected to luminescent HMGB1 immunoassay. Results shown represent the difference in luminescence between pulsed and non-pulsed conditions. Datapoints represent the average of two technical replicates from an individual biological experiment, with a total of n=6 independent experiments represented on the graph. A paired *t*-test was utilized to assess statistical significance. **f.** Antigen-pulsed or non-pulsed JY cells were conjugated at a 4:1 E:T ratio with CTLs for 4hr and supernatants subjected to LDH activity assay. Results shown indicate OD450 values (with background subtracted). Each datapoint represents the average of two technical replicates from an individual experiment, with n=6 independent experiments represented on the graph. A paired *t*-test was utilized to assess statistical significance. **g.** JY cells were pulsed or non-pulsed with antigenic peptide and conjugated at an E:T ratio of 4:1 with CTLs that had been pre-treated or not with monensin to de-acidify lytic granules. After 4hr conjugation, cells were subjected to Annexin V/PI staining and analyzed by flow cytometry. Data shown represent the percentage of target cells in the Annexin V/PI double-positive quadrant. Each datapoint represents the average of two technical replicates from an individual experiment, with a total of n=5 independent experiments represented on the graph. One-way ANOVA with Tukey’s test for multiple comparisons was utilized to assess statistical significance. **h.** The difference in percentage of double positive cells (Pulsed – Non-pulsed) from the experiment shown in **(g.)**. **i.** JY target cells were treated with 8μg/mL recombinant human perforin for 60min in perforin binding buffer, subjected to Annexin V/PI staining, and analyzed by flow cytometry. Each datapoint represents the average of two technical replicates from an individual experiment, with a total of n=3 biological experiments represented on the graph. Bars represent mean +/- SEM. Paired *t*-tests were utilized to assess statistical differences. **j.** JY cells were pre-treated with solvent or 100μM caspase-1 inhibitor VX-765 for 120mins before exposure to 8μg/mL recombinant human perforin for 60min. Cells were stained with Annexin V/PI and analyzed by flow cytometry. Data shown represents the difference between control and perforin-treated conditions for solvent-treated and VX-765-treated cells respectively. Each datapoint represents the average of two technical replicates from an individual experiment, with n=4 independent experiments shown on the graph. A paired *t*-test was utilized to assess statistical difference. \**p*<0.05, ** *p*<0.01, *** *p*<0.001, , **** *p*<0.0001, ns = not significant.

To assess cell death morphology in individual cells longitudinally, we utilized a semi-automated image-processing pipeline to analyze changes in target cell size and shape over time following CTL attack. Non-pulsed cells were analyzed as a control (**Supplementary Figure 10** and **Supplementary Movie 10**) and revealed the mild rhythmic oscillations in size (cross-sectional area) and shape (aspect ratio) typical of a live target cell. By contrast, upon CTL attack, dying target cells exhibited sharp, erratic fluctuations in cross-sectional area and aspect ratio concurrent with initiation of membrane bubbling, both of which stabilized upon cessation of cell motility (**Supplementary Figure 11a-d** and **Supplementary Movies 11-12**). These bubbling membrane protrusions combined with cell bloating caused transient expansion of target cells, with representative cells exceeding 160% of their original cross-sectional area (**Supplementary Figure 11b**). This phenotype closely resembled nigericin-induced pyroptosis, wherein cross-sectional area and aspect ratio varied dramatically within the bubbling phase before stabilization upon cessation of cell movement (**Supplementary Figure 12a-d** and **Supplementary Movies 4-5**), with representative cells swelling up to 200% of their initial cross-sectional area immediately before lysis (**Supplementary Figure 12d**). By contrast, apoptotic cells’ cross-sectional area and aspect ratio remained relatively consistent throughout the imaging period (**Supplementary Figure 13a-d** and **Supplementary Movies 8-9**). Collectively, these results indicated that dying target cells exhibited a morphology that was consistent with pyroptosis following CTL attack.

### Target cell death upon attack by human Ag-specific CTL is lytic and inflammatory

To determine whether CTL-mediated cell death also exhibited a molecular profile consistent with non-apoptotic cell death modalities, antigen-specific CTLs were conjugated with antigen-pulsed or non-pulsed target cells, subjected to Annexin V/PI staining and assessed by flow cytometry (gating strategy in **Supplementary Figure 14**). The Annexin V/PI assay is a highly sensitive and well-validated assay designed to distinguish both between early and late apoptosis, and between apoptosis and programmed necrosis^44^. It is based upon the premise that FITC-conjugated Annexin V binds to externalized phosphatidylserine (**PS**) on the outer membrane of apoptotic cells, prior to the loss of membrane integrity and uptake of vital dyes; thus, when combined with PI as a vital dye, the Annexin V/PI assay permits discrimination between live cells (double negative), early-stage apoptotic cells (Annexin V single-positive), and end-stage apoptotic or necrotic cells (Annexin V/PI double-positive)^44^. Necrotic cell death modalities (including pyroptosis) do not accumulate a sizeable Annexin V single-positive population, and instead Annexin V staining occurs concurrently with PI uptake^44^. Of note, under prolonged *in vitro* conditions wherein phagocytic clearance of apoptotic cells cannot occur, end-stage apoptotic cells tend to undergo a process of secondary necrosis (example shown in **Supplemental Figure 9d**), leading to the uptake of vital dyes and the eventual accumulation of an AnnexinV/PI double-positive population^44^. As such, the double-positive population is not a distinguishing feature between programmed necrosis and late apoptosis; the important distinguishing feature in apoptosis is the presence of an Annexin V single-positive population, which is absent during pyroptosis and other forms of necrotic PCD.

As shown in **Figure 4c-d**, Annexin V/PI double-positive target cells accumulated in a time-dependent fashion after 1hr and 4hr conjugation with CTLs. Conversely, the Annexin V single-positive population remained unchanged across all time points tested, consistent with a non-apoptotic cell death mechanism. This closely recapitulated the phenotype of nigericin-exposed pyroptotic cells, which likewise acquired a substantial double-positive population and failed to accumulate an Annexin V single-positive population (**Supplementary Figure 15a,b**). By contrast, a significant increase in the Annexin V single-positive population was observed following staurosporine exposure (**Supplementary Figure 15a,b**), a classical feature of apoptosis that is notably absent during acute CTL-mediated killing. Of note, the population of staurosporine-treated cells also contained a sizeable number of Annexin V/PI double-positive (secondary necrotic) cells, which is quite typical of end-stage apoptosis *in vitro*^44^.

To assess whether CTL-mediated cell death was associated with release of lytic or inflammatory molecules, supernatants were collected following conjugation of non-pulsed or antigen-pulsed target cells with CTLs for 1 or 4hr. As illustrated in **Figure 4e**, the classical alarmin and immunogenic cell death (**ICD**) marker HMGB1 was readily detectable in supernatant after 1hr. Likewise, supernatant LDH activity was detectable after 4hr conjugation (**Figure 4f**), consistent with a lytic cell death modality such as pyroptosis.

Collectively, these experiments indicated that target cell death upon CTL attack possessed the defining morphological features and molecular profile typical of pyroptosis, including rapid expansion and contraction of pyroptotic bodies, stochastic changes in cell size and shape, loss of membrane integrity concurrent with PS exposure, and overt membrane lysis indicated by supernatant HMGB1 and LDH activity.

### GSDMD and GSDME knockdown and knockout reduces CTL-mediated cytotoxicity

As a final demonstration that CTL-mediated cell death was indeed pyroptotic, we asked whether inhibition of pyroptotic pathways would have a measurable effect on target cell death upon CTL attack. As described above, JY cells express high levels of two pyroptotic executioners (GSDMD and GSDME) at the transcript and protein levels, which enables them to undergo pyroptosis either through the caspase-1/GSDMD pathway described above or the caspase-3/GSDME pathway identified by others^54,55^. To inhibit both pathways simultaneously, we transfected JY cells with either non-targeting siRNA or a combination of both *GSDMD*- and *GSDME*-targeting siRNAs; knockdowns were validated by immunoblot (**Supplementary Figure 16a**). Transfected target cells were pulsed or not with antigenic peptide and conjugated with CTLs for 4hr before being subjected to Annexin V/PI analysis, while supernatants were collected for LDH assay. As shown in **Supplementary Figure 16b**, concurrent knockdown of both pyroptotic pathways significantly inhibited the accumulation of Annexin V/PI double-positive cells (**Supplementary Figure 16b**), compared to the non-targeting siRNA condition; no increase in Annexin V single-positive cells was observed within this time frame, indicating that the inhibition of *GSDMD* and *GSDME* did not lead to a compensatory increase in apoptosis at these time points. However, knockdown of the pyroptotic pathways did prevent the loss of the double-negative population, which indicated that cells remained in a healthy state and failed to undergo either pyroptotic or apoptotic cell death in the time frame examined. No difference in the PI single-positive population was observed. To validate these observations in a knock-out system, these experiments were repeated using *GSDMD* and *GSDME* double knock-out target cells generated using the Crispr Cas9 system (validation shown in **Supplementary Figure 16c**). These results recapitulated our observations from the siRNA experiments, whereupon loss of both pyroptotic executioners significantly inhibited the accumulation of Annexin V/PI double-positive cells upon CTL attack (**Supplementary Figure 16d**). Collectively, these results illustrate that CTL-mediated end-stage cell death is dependent upon the pyroptotic executioners GSDMD and GSDME.

To further examine the effect of *GSDMD* and *GSDME* loss on the cell death phenotype, LDH activity was assessed in both non-targeting siRNA-transfected and *GSDMD*/*GSDME* double knockdown JY cells upon CTL attack. As anticipated, knockdown of both pyroptotic executioners led to a significant decrease in supernatant LDH activity (**Supplementary Figure 16e**) in pulsed conditions. Interestingly, caspase-1 and caspase-3 were equivalently activated in target cells bearing the *GSDMD/GSDME* double knockdown compared to non-targeting siRNA-transfected controls (**Supplementary Figure 16f,g**). This offered the crucial piece of insight that even when executioner caspases are fully activated in target cells upon CTL attack, those target cells do not die as efficiently without the pyroptotic executioners GSDMD and GSDME as caspase substrates.

### Target cell pyroptosis is dependent upon the perforin pathway and soluble perforin activates pyroptosis

To next determine if target cell pyroptosis was dependent upon the perforin pathway, we utilized monensin (described above) to attenuate the toxicity of the perforin/granzyme pathway through acidification of CTL lytic granules. As shown in **Figure 4g-h**, target cells conjugated with monensin-treated CTLs failed to accumulate an Annexin V/PI double-positive population, indicating that pyroptosis requires intact perforin signaling.

In a second approach, we assessed whether exposure of JY cells to soluble perforin could trigger pyroptotic cell death. To assess this, we exposed JY cells to recombinant perforin for 60min and assessed cell death utilizing the Annexin V/PI assay. Consistent with a pyroptotic cell death modality, perforin exposure resulted in a significant increase in Annexin V/PI double-positive cells (**Figure 4i**) but no increase in Annexin V single-positive cells (which would indicate apoptosis). Moreover, no increase in PI single-positive cells was apparent, which argued against overt (non-programmed) necrotic cell death occurring from perforin-induced osmotic shock. Unlike pyroptosis, non-programmed necrotic phenomena such as osmotic shock cannot be prevented or delayed through pharmacological or genetic intervention^56^; to further illustrate that perforin-mediated cell death was programmed rather than overtly necrotic, we pre-treated JY cells with VX-765, a well-validated, highly specific and cell-permeable peptidomimetic caspase-1 inhibitor^85,89,90^. Compared to solvent-treated controls, cells that were pre-treated with VX-765 showed a significant reduction in perforin-induced cell death (**Figure 4j**), illustrating that soluble perforin induced caspase-1-mediated PCD under these conditions.

With this insight, we next sought to determine whether the canonical pyroptotic pathway was activated upon CTL attack in a broader panel of transformed human cells.

### Melanoma cells activate pyroptosis upon CTL attack

To systematically test whether human melanoma cells could also undergo pyroptosis upon CTL attack, we performed Annexin V/PI assays using three different cellular systems, including D10 and LB2259-MEL.A melanoma cells that have been previously characterized^7,8^, as well as cells isolated on-site from a patient melanoma sample (Toulouse Melanoma 3, or TouM3). These different target cells were conjugated with CTLs for 4hr, subjected to Annexin V/PI staining, and assessed by flow cytometry.

As shown in **Figure 5a-b**, Annexin V/PI double-positive target cells accumulated in all cell lines tested upon 4hr conjugation with CTLs, whereas the Annexin V single-positive population remained unchanged compared to non-pulsed controls (**Figure 5c**), consistent with a pyroptotic cell death mechanism. The percentage of PI single-positive cells did not change in any cell line tested (**Figure 5d**).

**Figure 5.**
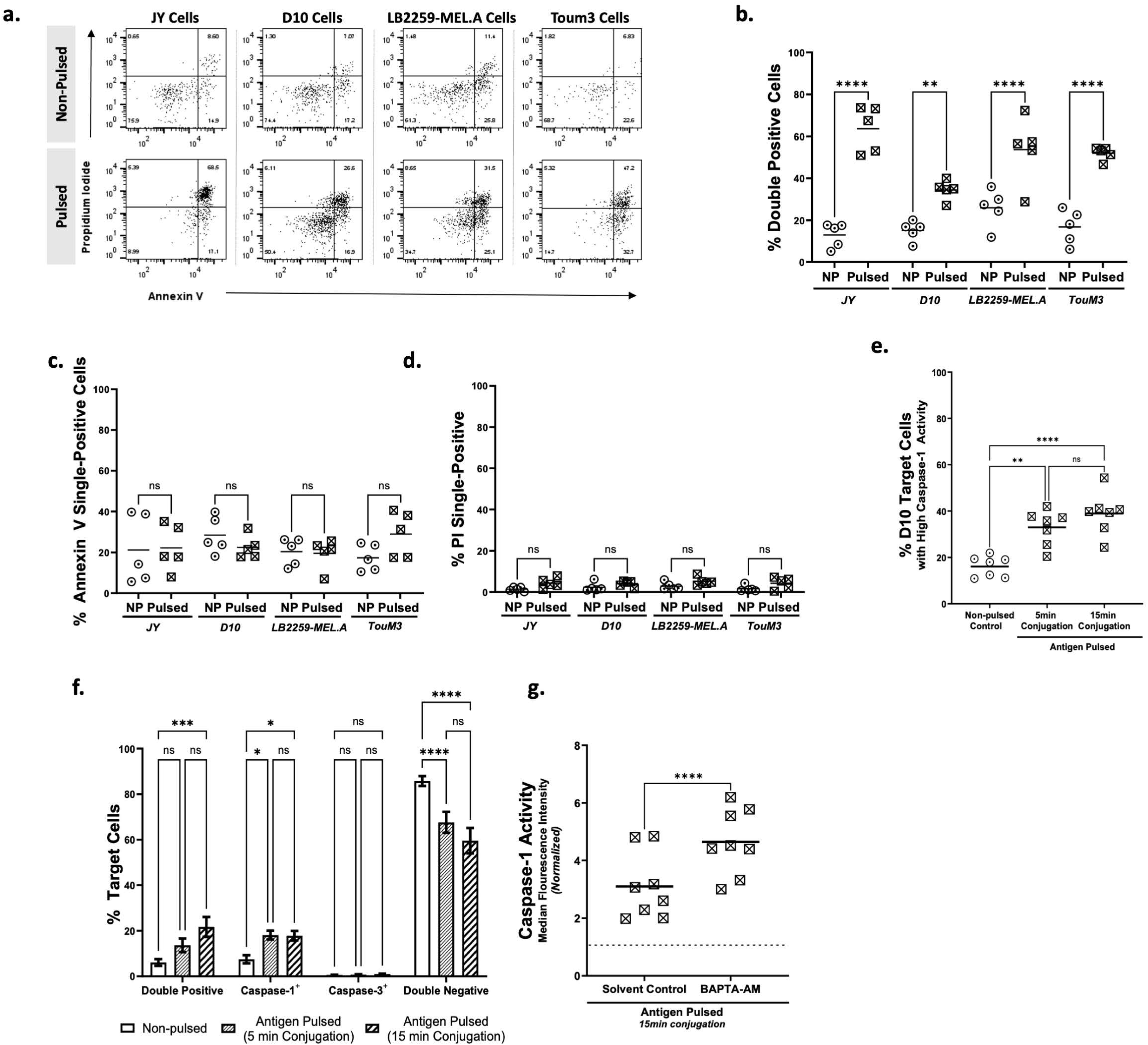
Melanoma cells display a pyroptotic phenotype upon CTL attack a-d. Melanoma cells (D10, RIPA, and TouM3) were pulsed or non-pulsed with 10μM antigenic peptide and conjugated at an E:T ratio of 4:1 with antigen-specific human CTLs for 4hr, with JY cells as a control. Following conjugation, cells were subjected to Annexin V/PI staining and analyzed by flow cytometry. **a.** Representative dotplots of target cells from n=5 independent experiments. **b-d.** Data shown represent the percentage of cells in each quadrant of interest (Annexin V/PI double positive, Annexin V single-positive, PI single-positive) after 4hr conjugation. Each datapoint represents the average of two technical replicates from an individual experiment, with a total of n=5 independent experiments represented on the graph. Black bars represent the mean of independent experiments. One-way ANOVA with Sidak’s test for multiple comparisons was utilized to assess statistical significance. **e.** D10 target cells were pulsed with 10μM antigenic peptide and conjugated with CTLs at an E:T ratio of 4:1 for 5 or 15min, before being stained with an activity-dependent caspase-1 probe and assessed using flow cytometry. Non-pulsed target cells were conjugated with CTLs under identical conditions for 15min as a control. Each datapoint represents the average of two technical replicates from an individual experiment, with n=7 independent experiments represented on the graph. One-way ANOVA with Tukey’s test for multiple comparisons was utilized to assess statistical significance. **f.** D10 cells were conjugated with CTLs as above and double-stained with activity-dependent caspase-1 and caspase-3 probes before being assessed by cytometry. Cells were classified as single-positive, double-positive, or double-negative for each active caspase at each timepoint shown. Bars represent mean +/- SEM, with n=7 independent experiments. RM two-way ANOVA with Tukey’s test for multiple comparisons was utilized to assess statistical significance. **g.** D10 cells were pre-treated with solvent control or the calcium chelator BAPTA-AM to inhibit calcium-dependent membrane repair mechanisms, conjugated with antigen-specific CTLs for 15min, and subjected to caspase-1 activity assay. Data represent fold increase in caspase-1 MFI, normalized to respective non-pulsed controls. Each datapoint represents the average of two technical replicates from an individual experiment, with n=8 independent experiments represented on the graphs respectively. Paired *t*-tests were utilized to assess statistical significance. \**p*<0.05, ** *p*<0.01, *** *p*<0.001, **** *p*<0.0001, ns = not significant.

Based on these observations, we utilized the D10 melanoma cell line (which is known to be moderately resistant to CTL attack due to enhanced reparative membrane turnover^7,8,58,81^) to determine whether partially resistant cells activate the caspase-1 and whether inhibition of defensive membrane turnover increases signaling through this pyroptotic pathway. After first verifying that D10 cells expressed NLRP3, caspase-1, GSDMD, and GSDME at the protein level at baseline (**Supplementary Figure 3a** and **Supplementary Figure 5b**), we next sought to determine whether caspase-1 was activated in D10 melanoma cells upon CTL attack. As shown in **Figure 5e**, D10 cells display a significant and time-dependent increase in caspase-1 activity upon CTL attack, but in a more limited fashion than JY cells (**Figure 2b**), after 5 and 15min conjugation. As with JY cells, caspase-1 activity was often observed in the same cells wherein caspase-3 activity could be detected, although a discreet population of caspase-1-only cells could also be identified (**Figure 5f**).

All of these results were consistent with the hypothesis that membrane repair mechanisms acting in melanoma cells to remove perforin pores from the target cell membrane^7,8^ might be responsible for dampening perforin-induced caspase activation. To test this concept, we sought to determine the impact of membrane repair on caspase-1 activation. We pre-treated pulsed or non-pulsed D10 melanoma cells with either BAPTA-AM, a calcium chelator which inhibits Ca^2+^-dependent reparative membrane turnover upon perforin pore formation without affecting perforin activity^7,10^ or solvent control before conjugating D10 melanoma cells with CTLs for 15min. Consistent with the hypothesis that inhibition of membrane reparative turnover would increase signaling through perforin-dependent pyroptotic pathways, we demonstrated that caspase-1 activity (**Figure 5g**) in pulsed D10 cells was significantly increased upon inhibition of Ca^2+^-dependent reparative membrane turnover compared to solvent-treated controls.

### Prolonged interaction with CTLs unveils a delayed apoptotic phenotype in multiple cell types

We next sought to investigate whether the natural depletion of intracellular perforin stores following prolonged interactions between targets and cytotoxic lymphocytes (which has been well-documented^59,91–93^) might inhibit the ability of CTLs to induce pyroptosis during the sustained phase of killing. It is known for both CTLs and NK cells that ultra-rapid perforin/granzyme-mediated killing ultimately gives way to slower complementary killing mechanisms such as death-receptor ligation, which is known to drive apoptosis^91–93^. Thus, we hypothesized that apoptotic target cells may be detectable following longer conjugations. To address this, cell death modality was assessed by Annexin V/PI assay following overnight conjugations of CTLs with four different cell types (**Figure 6a-d**). Under these conditions, a sizeable apoptotic subpopulation of Annexin V single-positive target cells was indeed detectable in all cell lines tested (**Figure 6b**), despite being largely absent from the target cell population at earlier timepoints (**Figure 5b**). This suggests that while the initial wave of perforin-dependent CTL-mediated cell death was lytic and inflammatory, traditional apoptotic cell death mechanisms came into play over time as lytic content became depleted.

**Figure 6.**
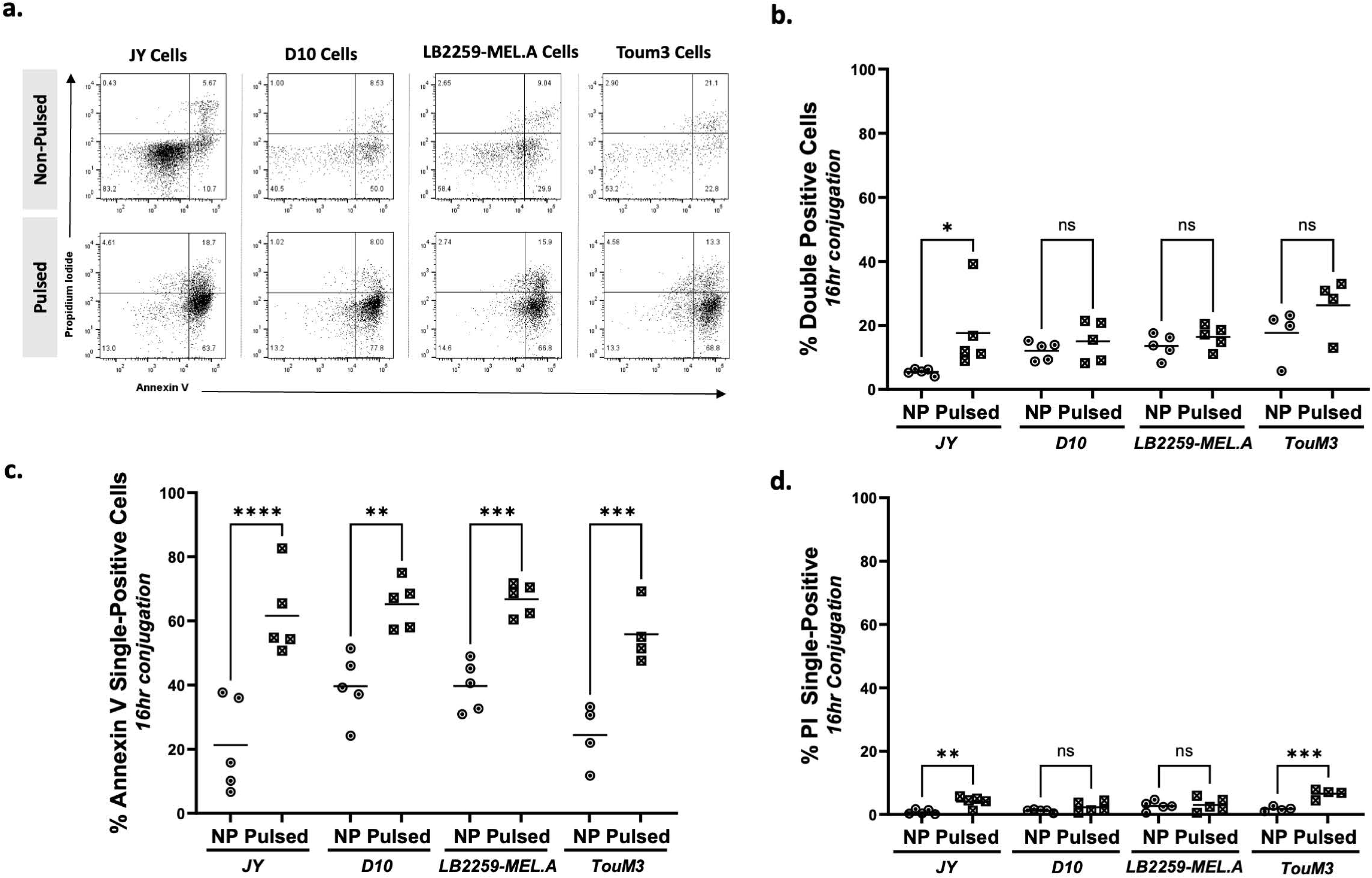
Pulsed cells conjugated overnight with antigen-specific CTLs reveal a delayed apoptotic phenotype a-d. JY and melanoma cells (D10, RIPA, and TouM3) were pulsed or non-pulsed with 10μM antigenic peptide and conjugated at an E:T ratio of 4:1 with antigen-specific human CTLs for 16hr. Following conjugation, cells were subjected to Annexin V/PI staining and analyzed by flow cytometry. Data shown represent the percentage of cells in each quadrant of interest (Annexin V/PI double positive, PI single-positive, Annexin V single-positive, ) at the indicated time points. Each datapoint represents the average of two technical replicates from an individual biological experiment, with a total of n=5 independent experiments represented on the graph. One-way ANOVA with Sidak’s test for multiple comparisons was utilized to assess statistical significance. \**p*<0.05, ** *p*<0.01, *** *p*<0.001, **** *p*<0.0001, ns = non-significant. NP = non-pulsed.

Collectively, the experiments in this manuscript have demonstrated the engagement of the canonical pyroptotic pathway upon CTL attack in multiple target cell types, downstream of target cell perforation and characterized by K^+^ efflux, NLRP3 inflammasome activation, caspase-1 activity, and GSDMD activation. The GSDMD pathway was engaged in parallel to the previously described granzyme B-dependent, GSDME-mediated pyroptotic pathway^54,55^ during the acute killing phase. Finally, sustained interaction with CTLs unmasked a delayed apoptotic phenotype in the remaining target cells after prolonged conjugations. **Figure 7** illustrates the multi-pronged cell death pathways triggered in target cells upon CTL attack.

**Figure 7.**
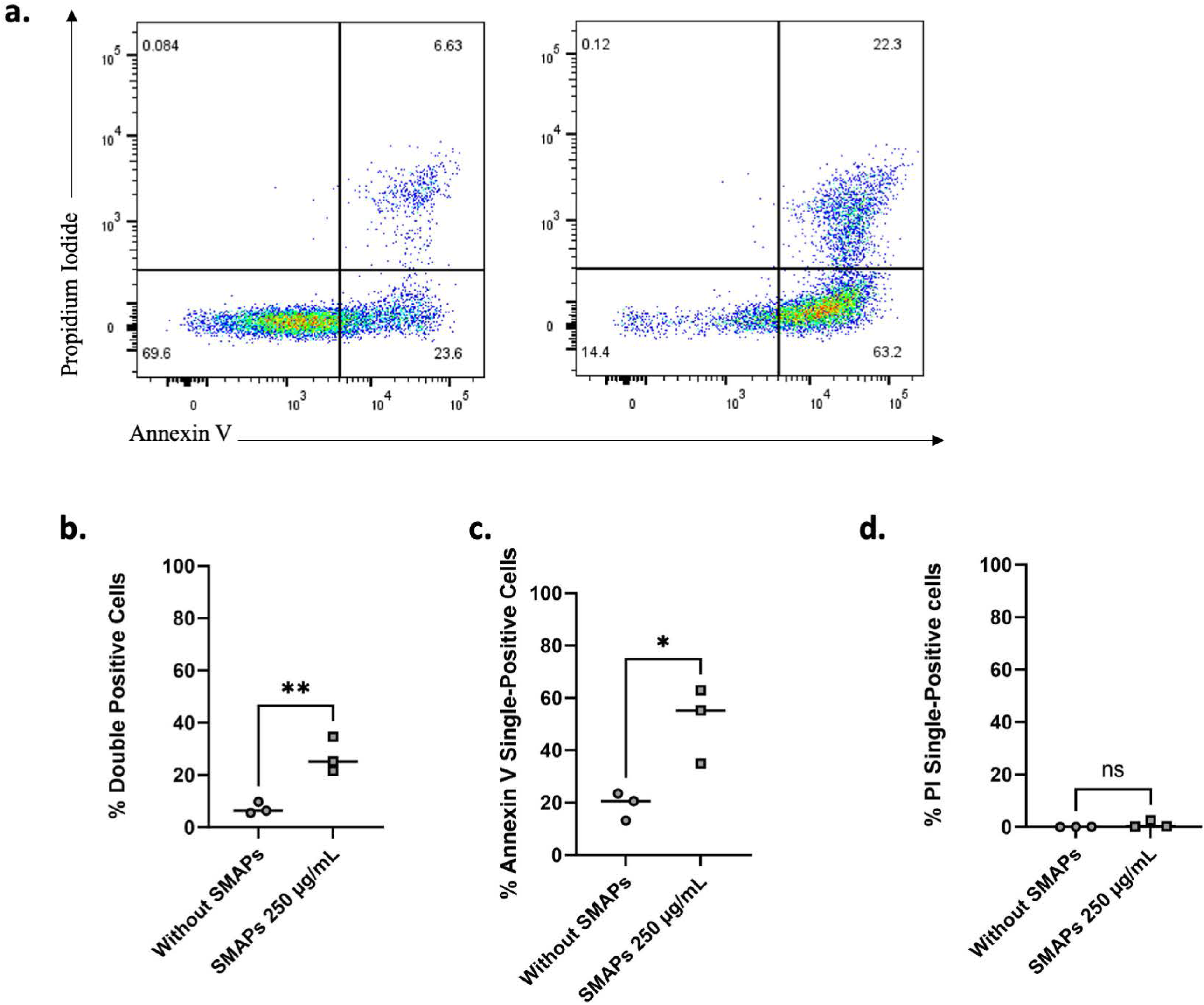
Exogenous SMAPs induce target cell apoptotic cell death a-d. JY cells were incubated with or without SMAPs at a protein concentration of 250 µg/mL for 4 hours. Cells were subjected to Annexin V/PI staining and analyzed by flow cytometry. Data indicate the percentage of cells in each quadrant: Annexin V/PI double positive, Annexin V single-positive and PI single-positive. Each datapoint represents the average of two technical replicates from an individual biological experiment with a total of n=3 independent experiments. Statistical differences between both conditions were assessed by unpaired *t*-test for b. and c. and by Mann-Whitney test for d. ** *p*< 0,01, \**p*<0,05, ns = not significant.

### Exogenous SMAPs trigger apoptosis

SMAPs are newly described components of the CTL and NK cell lytic arsenal that can serve as autonomous killing entities to annihilate target cells from a distance^4,94,95^. While SMAPs are the subject of intense investigation for their therapeutic potential in cancer patients, their mechanisms of action in killing target cells are still elusive. In particular, the potential of SMAPs to trigger either pyroptotic or apoptotic pathways has not been investigated.

To investigate SMAP-triggered cell death pathways JY cells were treated with exogenous SMAPs purified from NK-92 cell culture supernatant as described^5^. As shown in **Figure 7a,b**, exposure of JY cells to SMAPs for 4 hours induced only a moderate increase in the proportion of Annexin V^+^PI^+^ double-positive cells. By contrast, a consistent and significant increase in the Annexin V single-positive population was observed (**Figure 7c**), which is a feature unique to early apoptosis. No increase in PI single-positive cells was observed, confirming that necrotic (non-programmed) cell death did not occur. Together these results are compatible with a SMAP-mediated induction of apoptotic cell death in target cells.

Collectively, the experiments in this manuscript have demonstrated the engagement of the canonical pyroptotic pathway upon CTL attack in multiple target cell types, downstream of target cell perforation and characterized by K^+^ efflux, NLRP3 inflammasome activation, caspase-1 activity, and GSDMD activation. The GSDMD pathway was engaged in parallel to the previously described granzyme B-dependent, GSDME-mediated pyroptotic pathway^54,55^ during the acute killing phase. Sustained interaction with CTLs unmasked a delayed apoptotic phenotype in the remaining target cells after prolonged conjugations. Conversely exogenous SMAPs triggered the activation of early apoptotic cell death. **Supplementary Figure 17** illustrates the multi-pronged cell death pathways triggered in target cells by the diverse killer cell arsenal.

## DISCUSSION

In our study we provide two specific mechanistic insights into CTL-mediated target cell death: first, we identified a novel mechanism for the engagement of pyroptotic machinery upon CTL attack, in which perforin itself autonomously engages the caspase-1/GSDMD pyroptotic signaling pathway, and second, we demonstrated that isolated SMAPs fail to activate this pyroptotic pathway in order to induce a distinctly apoptotic cell death phenotype. These observations are consistent with the observed wave of early pyroptotic cell death immediately upon exposure of target cells to CTLs, that eventually gives way to slower apoptotic death in the remaining targets.

Our observation that CTLs trigger an initial wave of pyroptotic cell death supports previous studies that postulated a role for pyroptosis upon CTL attack^53–55^, and offer a novel mechanism for the engagement of pyroptotic machinery downstream of perforin-mediated pore formation. Conceptually, the pathway we identified links two different perforation events which may cooperate to promote target cell death: the initial perforation of target cells by CTL-derived perforin (which is spatially constrained to the LS and temporally constrained by ultra-rapid synaptic membrane repair mechanisms^7–9^) followed by a more global perforation of target cells by their own endogenous pore-forming gasdermin proteins. Although perforin has been shown to induce osmotic lysis (ie. necrotic non-programmed cell death) at lytic concentrations which is potentiated by blockade of membrane repair mechanisms^10^, this is the first report to our knowledge that ascribes a pro-pyroptotic function to perforin, alongside its classical role as a conduit for granzymes. The idea that perforin pore formation can activate the NLRP3/caspase-1/GSDMD pathway is consistent with a well-developed body of literature showing that cytotoxic PFTs, including *Aeromonas hydrophila* -derived aerolysin^22,23^, *Vibrio pyarahaemolyticus*-derived thermostable direct hemolysins (TDH)^24^, *Staphylococcus aureus-*derived Paton-Valentine leukocidin (PVL)^26^, *Trueperella pyogenes*-derived pyolysin (PLO)^27^ and *S. aureus-*derived α-hemolysin^22,25^, trigger caspase-1 activation in perforated cells via the NLRP3 inflammasome. The pores formed from these bacterial toxins are almost universally permeable to K^+^ (reviewed in^18^), and not surprisingly, NLRP3/caspase-1 activation in these contexts is often K^+^ efflux-dependent^22–27^. Thus, perforin functionally mimics the activity of the ancestral pore-forming bacterial toxins described above, demonstrating that this ancient attack mechanism retains utility in the human CTL arsenal today. This may be advantageous for CTLs attempting to circumvent granzyme-specific resistance mechanisms, such as the serine protease inhibitors (**serpins**), which have been shown to confer resistance to both chimeric antigen receptor (**CAR**) T cells and immune checkpoint blockade (**ICB**) therapy^96,97^. As such, engineering immunotherapies that potentiate perforin-activated pyroptotic signaling pathways may be a useful complementary approach. Of note, loss of intracellular potassium homeostasis downstream of membrane perforation by bacterial PFTs has also been suggested to trigger other adaptations in addition to inflammasome activation (including arrest of protein synthesis, activation of the mitogen-activated protein kinase (MAPK) pathway, and induction of autophagy)^18,98^; the extent to which such downstream responses occur upon perforin pore formation in tumor cells, and if so whether they ultimately enhance or attenuate CTL cytotoxicity, remains to be determined.

Although SMAPs have been shown to contain significant quantities of perforin, it and other cytotoxic molecules are understood to be largely encapsulated within the SMAP’s impermeable glycoprotein shell^4^. This shell may prevent soluble perforin from encountering the target cell membrane, thus limiting the crucial perforation step that leads to the K^+^ efflux and inflammasome activation and encouraging a slower apoptotic form of cell death. Whether the alternative pyroptotic GSDME pathway is not triggered by SMAPs that contain granzyme B is presently elusive. It is tempting to speculate that since the uptake of SMAPs by target cells should not occur via perforin pores, but rather be mediated by the endocytic compartment as we observed in CTL/target cell conjugates^99^, the SMAP-associated granzyme B might trigger a different death cascade than that triggered by the perforin-dependent cytosolic granzyme B.

We and others have shown that ultra-rapid Ca^2+^-dependent membrane repair is a vital resistance mechanism against CTL attack^7–10^. Our current data implies that K^+^ -dependent pro-death signalling may be initiated by the same perforation event as Ca^2+^-dependent pro-survival repair mechanisms. Previous studies by ourselves and others have shown that BAPTA-AM-mediated inhibition of Ca^2+^-dependent synaptic membrane repair results in an overt increase in target cell death in response to either perforin exposure or CTL attack^7,10^, and the current study suggests that promotion of caspase-1-dependent pyroptosis may be one mechanism by which this occurs. Importantly, Ca^2+^-dependent membrane repair mechanisms also facilitate the removal of gasdermin pores from the plasma membrane^39,40^, and thus inhibition of membrane repair might be a doubly effective strategy for promoting both effective target perforation by perforin and ultimately gasdermin-mediated cell death.

Of note, Ca^2+^ flux has also been proposed as a common mediator of NLRP3 inflammasome activation^21,100^, but in this study, chelation of intracellular Ca^2+^ with BAPTA-AM in D10 melanoma cells increased caspase-1 activation, suggesting that inflammasome activation in this system is not dependent on Ca^2+^ flux.

As mentioned above, the effect of siRNA-mediated NLRP3 inhibition on caspase-1 activity was modest in this system, and this observation is consistent with the presence of multiple other inflammasomes which may promote caspase-1 activation upon CTL attack^101^. Concurrent activation of other inflammasome sensors in combination with NLRP3 is well-documented (reviewed in ^21^) and NLRC4 in particular has been shown to be activated alongside NLRP3 upon membrane perforation by bacterial PFTs^24^. Likewise, NLRP1 can be activated upon disruption of intracellular K^+^ homeostasis^68^ and may serve as an additional platform for caspase-1 activation in our system. In addition to detecting loss of intracellular K^+^, inflammasome sensors also collectively detect a diverse array of danger signals (including abnormal protease activity, abnormal localization of nucleic acids, mitochondrial dysfunction, ROS production, disruption of the actin cytoskeleton, ribotoxic stress, and others)^101^. Many of these intracellular danger signals are present upon CTL attack^3^, which may firstly ensure vigorous and continuous caspase-1 activation in the presence of continued danger and secondly make the activation of caspase-1 relatively robust to the depletion of an individual inflammasome sensor (such as NLRP3). Nonetheless, the results of our study align with a previous report that showed that antigen-dependent CTL feedback can induce NLRP3-dependent caspase-1 activation in antigen-presenting cells (**APCs**) through a perforin-dependent and K^+^ -dependent mechanism, although these interactions were non-lethal and instead led to sustained IL-1β secretion by viable APCs^102^.

In this study, we provide evidence that caspase-1 is activated upon human CTL attack, as well as upon exposure of target cells to recombinant human perforin (**Figure 2**). In addition to promoting GSDMD-mediated pyroptosis, caspase-1 activation may have additional downstream consequences; proteomics studies have revealed dozens of substrates for caspase-1, including cytoskeletal proteins, chaperone proteins (including heat shock proteins), and trafficking proteins (such as RAB7), as well as cytokines (pro-IL1β and pro-IL-18) and a notable constellation of proteins involved in glycolysis^103,104^. Although GSDMD is thought to be cleaved the fastest^103^, this does not preclude a role for other caspase-1 substrates in the target cell response to CTL attack, which may become particularly relevant in the context of prolonged or sub-lethal encounters. Further studies will be necessary to elucidate other functions of caspase-1 prior to induction of CTL-driven pyroptosis.

Building upon previous studies of GSDME-mediated pyroptosis upon CTL attack^54,55^, we also provide evidence in this study that concurrent inhibition of both GSDMD and GSDME diminishes specific killing upon CTL attack, despite robust caspase-1 and -3 activation (**Supplementary Figure 16**). This suggests that caspase-1 and -3 activity alone is insufficient to achieve fast, optimal perforin/granzyme-mediated cytotoxicity and supports a role for gasdermins as the final executioners of target cell death. A multi-pronged approach in which both GSDMD and GSDME are activated upon CTL attack would be particularly important in transformed cells wherein individual gasdermins can be functionally inhibited through repression, inactivation, or mutation (reviewed in ^3,47,105^). Thus, activation of multiple pyroptotic pathways upon CTL attack would increase the probability of successfully circumventing such resistance mechanisms. The evolution of multiple gasdermins has been speculated to serve precisely this purpose, offering multiple redundant or partially redundant cell death pathways that are activated during cellular stress^28^.

More globally, the initiation of pyroptotic signaling pathways may be particularly vital in transformed cells wherein apoptotic pathways are inhibited, while induction of apoptosis by SMAPs may be crucial in transformed cells whose pyroptotic pathways are inhibited. Thus, the activation of two pyroptotic pathways (by perforin and granzyme respectively) upon CTL attack offers functional redundancy to circumvent inhibition of individual pyroptotic pathways, whereas the activation of an apoptotic pathway by SMAPs offers more global redundancy in the context of dual inhibition of both pyroptotic pathways.

Typically, plasma membrane permeabilization during pyroptosis is considered the main mechanisms of gasdermin-mediated toxicity; however, gasdermins have been shown to target other organelles enriched in cardiolipins, including lysosomes and mitochondria^80,106^, causing mitochondrial depolarization, disruption of the mitochondrial network, release of mtDNA and mitochondrial proteins, and lysosome deacidification/decay, which ultimately lead to a failure of cell viability prior to PMR^80,107^ (reviewed in ^35^). Although we observed PMR in target cells in our system using live-cell imaging (**Figure 4**), the extent to which intracellular gasdermin-mediated injures contribute to target cell demise prior to lysis remains to be characterized.

Despite the aforementioned publications demonstrating roles for GSDME and GSDMB in CTL-mediated pyroptosis^53–55^, the notion that CTL attack activates pyroptosis remains controversial within the field. Historically, apoptosis was the only well-recognized form of RCD in the literature and thus the notion that CTLs killed by apoptosis became a well-established paradigm. In this study, we provide evidence for activation of caspase-1/GSDMD upon CTL attack and demonstrate that cytotoxicity of antigen-specific human CTLs is functionally diminished through either siRNA-mediated inhibition of GSDMD and GSDME expression or Crispr Cas9-mediated *GSDMD* and *GSDME* knockout in target cells, providing proof-of-principle that target cells are capable of dying by pyroptosis under the conditions of this study. Although these results do not suggest that the majority of CTL-driven cytotoxicity events across all species, diseases, and contexts is pyroptotic in nature, they do provide ample rationale for considering pyroptosis as a viable alternative mechanism for perforin/granzyme-mediated target cell death. Our results are consistent not only with the recent studies directly implicating pyroptosis in perforin/granyzme-mediated killing^53–55^, but also with a number of earlier studies that highlighted non-apoptotic and/or immunogenic cell death modalities upon CTL attack^48–52^. Collectively, these studies involve a number of different contexts, across both mouse and human model systems, *in vitro* and *in vivo*, illustrating that our experimental conditions are not unique in promoting pyroptotic cell death upon CTL attack. At the same time, our observation of a SMAP-mediated apoptotic phenotype highlights another mechanism by which observations of apoptotic cell death and pyroptotic cell death across different study conditions may be reconciled.

Further to this, it is worth emphasizing that cell death modality upon CTL attack is almost certainly context-dependent. On the CTL side, even clonal human CTLs are phenotypically and functionally heterogeneous, with notable differences in the expression of cytotoxic molecules such as perforin and granzymes across clonal populations^108–111^. Given that different classes of cytotoxic molecules have discrete biological functions, the effect of a given CTL on a target cell may depend at least partially upon its individual cytotoxic payload. CTLs subpopulations with high perforin expression but relatively low expression of granzymes have been observed^110,111^ and these cells may indeed be endowed with perforin-dependent, granzyme-independent cytotoxic mechanisms; conversely, CTLs with lower perforin expression may preferentially induce killing through a SMAP-mediated apoptotic pathway. Indeed it has previously been shown that the presence or absence of certain lytic molecules can change not just the extent but also the modality of CTL-mediated killing^50^, with

CTLs deficient in both granzyme A and B inducing a non-apoptotic PCD modality characterized by swelling, membrane rupture, and simultaneous onset of PI uptake with Annexin V staining^50^. As such, it stands to reason that CTL-intrinsic factors play a role in determining target cell death modality. Further live-cell imaging studies could reveal at the single-cell level whether individual CTLs preferentially induce a specific cell death modality, or whether individual CTLs trigger different cell death modalities as a result of different cytotoxic mechanisms engaged sequentially during serial killing (as has been proposed for NK cells^93^, reviewed herein^112^).

On the target cell side, cancer cells are highly heterogeneous, both at the intra-tumoral and inter-tumoral level, in terms of differentiation state, gene expression profile, mutational burden, presence or absence of viral genomic integration, epigenetic modifications, and ultimately functional behaviour^113^. This heterogeneity, particularly as it pertains to the dysregulation of cell death machinery, likely plays a large role in determining cell death modality. Dysregulation of apoptotic circuitry specifically was one of the six foundational hallmarks of cancer identified by Hanahan and Weinberg^114^ and resistance to cell death through mutation, suppression, or dysregulation of cell death machinery (both apoptotic and pyroptotic) continues to be considered a defining characteristic to the present day and major functional obstacle to CTL-mediated therapies^3,115,116^. The integration of viral genomes into cancer cells (particularly EBV, hepatitis B and human papilloma virus)^117^ may introduce an additional level of cell death suppression through specific virally encoded modulators of cell death pathways. Thus, since an attacking CTL may face a transformed target wherein one or more cell death pathways have been rendered functionally impotent, the apoptosis-versus-pyroptosis decision balance is influenced by which cell death pathways still retain functionality following oncogenic transformation. Because multiple cell death pathways are initiated in parallel by an attacking CTL, it would not be unexpected that dying targets likewise display characteristics of multiple cell death modalities; some of these signaling pathways may be prematurely terminated due to the mutation, suppression, or inactivation of key components, while others may be actively involved in executing the cell death decision. The presence or absence of membrane repair mechanisms might also affect cell death modality, and indeed there is precedent in the literature to suggest that reparative membrane turnover promotes a slower, apoptotic cell death modality whereas inhibition of these Ca^2+^-dependent repair mechanisms unleashes a fast, lytic, and non-apoptotic cell death modality following target perforation^10^. For all of these reasons, cell death modality upon CTL attack is likely to be context-dependent and may not fit neatly into a single well-described PCD modality due to the multi-pronged nature of CTL attack.

In the context of oncology, multiple chemotherapeutics^29,118,119^, small molecule inhibitors^120^, radiation^121–123^, and tumor immunotherapy^53,55^ as well as nanoparticle-based approaches for therapeutic delivery of gasdermins^39,124^ have been shown to drive tumor cell pyroptosis, which strongly favours anti-tumor immunity and tumor clearance, particularly in the context of turning “cold” tumors “hot” (reviewed in ^105,125^). Mass spectrometry-based human proteomics studies have revealed that GSDMD-mediated pyroptosis releases several hundred different proteins, through a combination of conventional secretion, unconventional secretion, vesicle/lysosome release, and passive cell lysis (e.g. LDH)^126^. The pyroptotic secretome is particularly enriched in DAMPs (e.g, HMGB1, galectins, S100 family proteins, and lysosomal cathepsins) which collectively serve as potent proinflammatory danger signals in the extracellular environment, through various mechanisms including direct TLR ligation^126^. Activation of local inflammatory signaling cascades by the pyroptotic secretome potentiates anti-tumor immunity by acting on multiple steps of the cancer-immunity cycle^127^, including enhanced antigen presentation, priming of APCs, and immune infiltration, as reviewed elsewhere^105,125^. In addition, recent studies have suggested that the pyroptotic corpse itself may be directly immunogenic due to “crowning” of the corpse by F-actin-rich filopodia that engage the F-actin receptor CLEC9A on dendritic cells and promote engagement of the adaptive immune system^128^. As such, our results fit within a mature field of literature that has repeatedly highlighted how pyroptotic cell death can serve as a bridge between inflammation and adaptive immunity.

Nonetheless, the inflammation induced by pyroptosis is a double-edged sword, which has been associated with tumorigenesis^105^ as well as adverse inflammatory side effects including cytokine release syndrome^55,129^ and certain side-effects of chemotherapeutics^29^. Since cell death modality is known to have such a profound impact on treatment efficacy and side effect profile, one can envision there would be clinical utility in delineating how given CTL-mediated immunotherapies induce cell death. Given that expression of gasdermin family proteins is heterogeneous both within and between tumor types (reviewed in ^47,105^), it is conceivable that individual tumors have a pre-existing “fingerprint” of gene expression which sensitizes them to different cell death modalities and may impact efficacy and side effect profile of CTL-mediated therapies. Depending upon whether it is clinically advisable to promote or inhibit inflammatory cell death, different combination therapies could be envisioned that are designed to enhance or prevent pyroptosis. Proof of concept for this has already been demonstrated with chemotherapeutics, wherein the methyltransferase inhibitor decitabine enhanced GSDME expression and promoted pyroptosis following nanoparticle-mediated delivery of cisplatin^118^.

In the context of CTL-mediated immune surveillance against cancer, the observation that perforin and SMAPs trigger pyroptosis and apoptosis respectively is intriguing. While perforin-mediated cell death might serve as immunogenic cell death (**ICD**), important to trigger and amplify immune response, SMAP-induced apoptosis might serve as a source of tumor antigens for cross-priming. Accordingly, SMAPs contain CCL5 and XCL2 that are chemoattractants of cDC1, the key DC subtype responsible for cross-priming anti-tumor CD8^+^ T cells^4,130^. It is tempting to speculate that the parallel activation of perforin-mediated pyroptosis in some tumor cells and of SMAP-mediated apoptosis in other cells might alert the immune system while ensuring antigen presentation, resulting in sustained and efficient immune responses. Harnessing the apoptotic/pyroptotic cell death balance in cancer patients might become a novel and interesting path to follow in the development of immunotherapeutic strategies against cancer.

Outside the context of tumor immunology, understanding whether CTLs and/or CTL-derived perforin can trigger target cell or bystander cell pyroptosis may be crucially important for neuroinflammatory conditions (such as multiple sclerosis^85,131^), autoimmune disease, and chronic infection, wherein canonical GSDMD-dependent pyroptosis has been implicated as a driver of chronic inflammation^47,132,133^. In such conditions, the inflammatory pyroptotic secretome perpetuates bystander tissue damage, leading to a feed-forward cycle of tissue damage and inflammation^47,132^; as such, understanding and preventing CTL-driven pyroptosis may be a widely applicable therapeutic approach. Perforin itself has been identified as a driver of tissue damage in chronic viral infections, and its inhibition has quantifiable protective effects in various models^134^.

In conclusion, our study provides evidence that perforin-mediated pore formation at the LS leads to potassium efflux, triggering a proinflammatory NLRP3/caspase-1/GSDMD pyroptotic signaling cascade. We propose a novel pro-pyroptotic function for perforin, which is consistent with the role of prokaryotic pore-forming toxins and expands its cytotoxic role beyond that of a simple conduit for granzymes. Conversely, we show that SMAPs preferentially trigger apoptosis, thus providing a complementary source of antigens for anti-tumor immune responses. Our results highlight a novel mechanism for the engagement of pyroptotic machinery upon CTL attack and highlight the complexity and diversity of the human CTL lytic arsenal.

## METHODS

### Cell Culture

Cell culture was performed as previously described^7,8^. Human CD8^+^ T cells were purified from healthy donor blood samples using the RosetteSep Human CD8^+^ T Cell Enrichment Cocktail (StemCell Technologies). For cloning, HLA-A2-restricted CD8^+^ T cells specific for the NLVPMVATV peptide of the cytomegalovirus protein pp65 were single-cell sorted into 96-U-bottom plates using a BD FACSAria II cell sorter using tetramer staining. Cells were cultured in RPMI 1640 medium supplemented with 5% human AB serum (Inst. Biotechnologies J.BOY), 50µmol/L 2-mercaptoethanol, 10mM HEPES, 1X MEM NEAA (Gibco), 1X Sodium pyruvate (Sigma), 10µg/ml ciprofloxacine (AppliChem), 100 IU/ml human rIL-2 and 50 ng/ml human rIL-15 (Miltenyi). CD8^+^ T-cell clones were stimulated in complete RPMI/HS medium containing 1 μg/ml PHA with 1 x 10^6^ per ml 35 Gy irradiated allogeneic peripheral blood mononuclear cells (isolated on FicollPaque Gradient from blood of healthy donors). Clones were restimulated every 2 weeks. Blood samples were collected and processed following standard ethical procedures after obtaining written informed consent from each donor and approval by the French Ministry of Research (transfer agreement AC-2020-3971). Approbation by the ethical department of the French Ministry of the Research for the preparation and conservation of cell lines and clones starting from healthy donor human blood samples has been obtained (authorization no. DC-2021-4673). Approbation by the ethical department of the French Ministry of the Research for the preparation and conservation of cell lines issued from cancer patients’ tissue samples has been obtained (authorization N° DC-2021-4673) and written informed consent was obtained from each donor.

EBV-transformed B cells (JY) HLA-A2^+^ were used as sensitive target cells. D10 HLA-A2^+^ human melanoma cells were kindly provided by Dr G. Spagnoli, Basel, Switzerland. LB2259-MEL.A HLA-A2^+^ human melanoma cells were kindly provided by Dr P. Coulie and N.V. Baren, Brussels, Belgium. TouM3 cells were isolated in-house from primary melanomas, which were first dissociated using the Human Tumor Dissociation Kit (MiltenyiBiotec) on the GentleMACS Dissociator. Total cells were then filtered on a 70µm filter (BD Falcon) and red blood cells eliminated by ACK treatment. Tumor-infiltrating lymphocytes (**TILs**) were removed by positive selection using a TIL Isolation Kit (MiltenyiBiotec) by magnetic separation (LS Column) and the negative fraction of this separation (corresponding to retained cells) was incubated with magnetic beads from the Tumor Cells Isolation Kit (MiltenyiBiotec) and separated by magnetic sorting using LS column. After isolation, melanoma cells were cultured in T25cm^2^ flasks for several days and amplified twice before freezing (FCS/90%DMSO).

All target cells were cultured in RPMI 1640 GlutaMAX supplemented with 10% FCS and 50µmol/L 2-mercaptoethanol, 10mM HEPES, 1X MEM NEAA (Gibco), 1X Sodium pyruvate (Sigma), 10µg/ml ciprofloxacine (AppliChem). All cell lines were screened biweekly for mycoplasma contamination using MycoAlert mycoplasma detection kit (Lonza). Melanoma and JY cells were subjected to cell line typing (Microsynth Ecogenics, Switzerland) to authenticate their identity and ensure that no contamination of human origin was detectable.

### Drug treatments

To induce pyroptosis, cells were plated at a density of 2.5 x 10^4^ cells per well in a 96-well plate, treated with 100μM nigericin (CliniSciences #ICT-6698 or Invivogen #tlrl-nig) and incubated at 37°C and 5% CO2 for the indicated time (2hr unless otherwise indicated). This high concentration of nigericin was utilized to ensure that the majority of cells underwent swift and definitive pyroptosis and to circumvent the need for an LPS-priming step. To induce apoptosis, cells were plated using the same conditions and treated with 6.6μM staurosporine (*Streptomyces*) (InSolution, Merck #569396) and incubated at 37°C and 5% CO2 for the indicated time.

To assess the effect of Ca^2+^ chelation in target cells on pyroptotic signaling pathways, target cells were treated with 50µM BAPTA-AM for 2hr at 37°C (ThermoFisher Scientific, #B6769) and thoroughly washed before conjugation with CTLs.

To test the effect of CTL lytic granule depletion, CTLs were pre-treated with monensin (FisherScientific GolgiStop^TM^ containing 0.26% monensin, diluted 1:1000, #BDB554724) for 2hr at 37°C and thoroughly washed before conjugation with targets.

To assess the effects of caspase-1 inhibition, targets were pre-treated for 2hr with 100μM VX-765 (InvivoGen, #inh-vx765) during the 2hr antigen pulse step and washed prior to conjugation with CTLs.

### Annexin V/PI Apoptosis Detection

Target cells (JY and D10) were either non-pulsed or pulsed with 10µM CMV p65 antigenic peptide for 2hr at 37°C and 5% CO2, washed and transferred to a 96-well U-bottom plate at 1 x 10^4^ cells per 100µl RPMI 5% FCS/HEPES. CTL were previously stained with CellTrace^TM^ Violet (ThermoFisher Scientific, #C34557) for 30min at 37°C and 5% CO2, washed and added to the target cells at an effector:target ratio of 4:1 in 100µl RPMI 5% FCS/HEPES. Plates containing samples were spun for 1min and incubated at 37°C and 5% CO2 for either 1 or 4hr. At the endpoint, cells were spun, washed in ice cold PBS, and assayed for Annexin V and PI staining according to manufacturer’s instructions (BD Pharmingen FITC Annexin V Apoptosis Detection Kit I, #AB_2869082). In a subset of experiments, CTLs were treated with monensin (BD GolgiStop^TM^; #51-2092KZ) at a dilution of 1:1000 for 2hr and washed thoroughly prior to conjugation with targets. Samples were acquired using the MACSQuant Analyzer 10 flow cytometer (Miltenyi Biotec) and analyzed using FlowJo 10.

For drug controls, target cells were plated at a density of 2.5 x 10^4^ cells per well in a 96-well U-bottom plate, incubated with either nigericin (pyroptosis control), staurosporine (apoptosis control), or solvent controls for the indicated time, and subjected to Annexin V and PI staining as described above.

### Lactate Dehydrogenase Supernatant Activity Assay

Target cells (JY and D10) were either non-pulsed or pulsed with 10µM CMV p65 antigenic peptide for 2hr at 37°C and 5% CO2, washed and transferred to a 96-well U-bottom plate at 1 x 10^4^ cells per 100µl RPMI 5% FCS/HEPES. CTL were added to the target cells at an effector:target ratio of 4:1 in 100µl RPMI 5% FCS/HEPES. Plates containing samples were spun for 1min and incubated at 37°C and 5% CO2 for 4hr. At the endpoint, cells were spun and supernatant removed for storage at -20°C and then analyzed using the CytoSelect^TM^ LDH Cytotoxicity Assay Kit (Cell Biolabs Inc, #CBA-241) according to manufacturer’s instructions. Absorbance (optical density at 450nm) was assessed using a CLARIOstar plate reader (BMG Labtech), with background subtracted (optimal density at 650nm).

As a positive control, JY and D10 target cells were plated at a density of 2.5 x 10^4^ cells per well in a 96-well U-bottom plate and incubated with 100μM nigericin or solvent controls for 2hr. Supernatants were harvested and subjected to LDH detection as described above.

### HMGB1 Immunoassay

Target cells (JY and D10) were either non-pulsed or pulsed with 10µM CMV p65 antigenic peptide for 2hr at 37°C and 5% CO2, washed and transferred to a 96-well U-bottom plate at 2.5 x 10^4^ cells per 100µl RPMI 5% FCS/HEPES. CTL were added to the target cells at an effector:target ratio of 4:1 in 100µl RPMI 5% FCS/HEPES. Plates containing samples were spun for 1min and incubated at 37°C and 5% CO2 for 1hr. At the endpoint, supernatant was removed for immediate analysis using the LumitTM HMGB1 (Human/Mouse) Immunoassay (Promega, #W6110). Luminescent samples were read in opaque white 96-well plates using a CLARIOstar plate reader (BMG Labtech).

As a positive control, JY and D10 target cells were plated at a density of 2.5 x 10^4^ cells per well in a 96-well U-bottom plate and incubated with 100μM nigericin or solvent controls for 2hr. Supernatants were harvested and subjected to HMGB1 assay as described above.

### Caspase-1 and Caspase-3 Activity-Dependent Fluorescent Assays

Target cells were either non-pulsed or pulsed with 10µM CMV p65 antigenic peptide unless otherwise indicated for 2hr at 37°C and 5% CO2, washed and transferred to a 96-well U-bottom plate at 1 x 10^4^ cells per 100µl RPMI 5% FCS/HEPES. Plated target cells were single-or double-stained with either far-red fluorescent FLICA 660 Caspase-1 (YVAD) reagent (ImmunoChemsitry, #9122) or green fluorescent FAM-FLICA Caspase-3/7 (DEVD; detects active form of both caspase-3 and -7) reagent (ImmunoChemsitry, #93), according to manufacturer’s instructions for 20min at 37°C and 5% CO2. CTL were stained with CellTrace^TM^ Violet (ThermoFisher Scientific, #C34557) for 30min at 37°C and 5% CO2, washed and added to the target cells at an effector:target ratio of 4:1 unless otherwise indicated in 100µl RPMI 5% FCS/HEPES. Plates containing samples were spun for 1min and incubated at 37°C and 5% CO2 for either 5 or 15min. At the endpoint, cells were washed, fixed with the provided fixative (ImmunoChemistry #636), and stored at 4°C before being acquired using the MACSQuant Analyzer 10 flow cytometer (Miltenyi Biotec) and analyzed using FlowJo 10. In a subset of experiments, CTLs were treated with monensin (BD GolgiStop^TM^; #51-2092KZ) at a dilution of 1:1000 for 2hr and washed thoroughly prior to conjugation with targets.

As a positive control, JY and D10 target cells were plated at a density of 2.5 x 10^4^ cells per well in a 96-well U-bottom plate and incubated with 100μM nigericin or solvent controls for 2hr, before being stained with FLICA 660 Caspase-1 (YVAD) reagent for 20 mins and analyzed as described above.

### Transfection of Target Cells

From 8 x 10^5^ to 1x 10^6^ JY cells were washed and resuspended in 100µl OPTI-MEM and transferred in electroporation cuvettes (0.2cm). 200 pmoles of each different siRNA targeting mRNA coding for *GSDMD*, *GSDME*, *CASP1*, *NLRP3*, or non-targeting siRNA were added in the cell suspension. After electroporation at 160V during 10 ms (500µF) with a exponential waveform using the Gene PulserXcell Electroporation systems (Bio-rad), transfected JY cells were seeded in pre-warmed culture medium and used for experiments 24-48h after electroporation. Transfection efficiency was evaluated by qPCR or immunoblot.

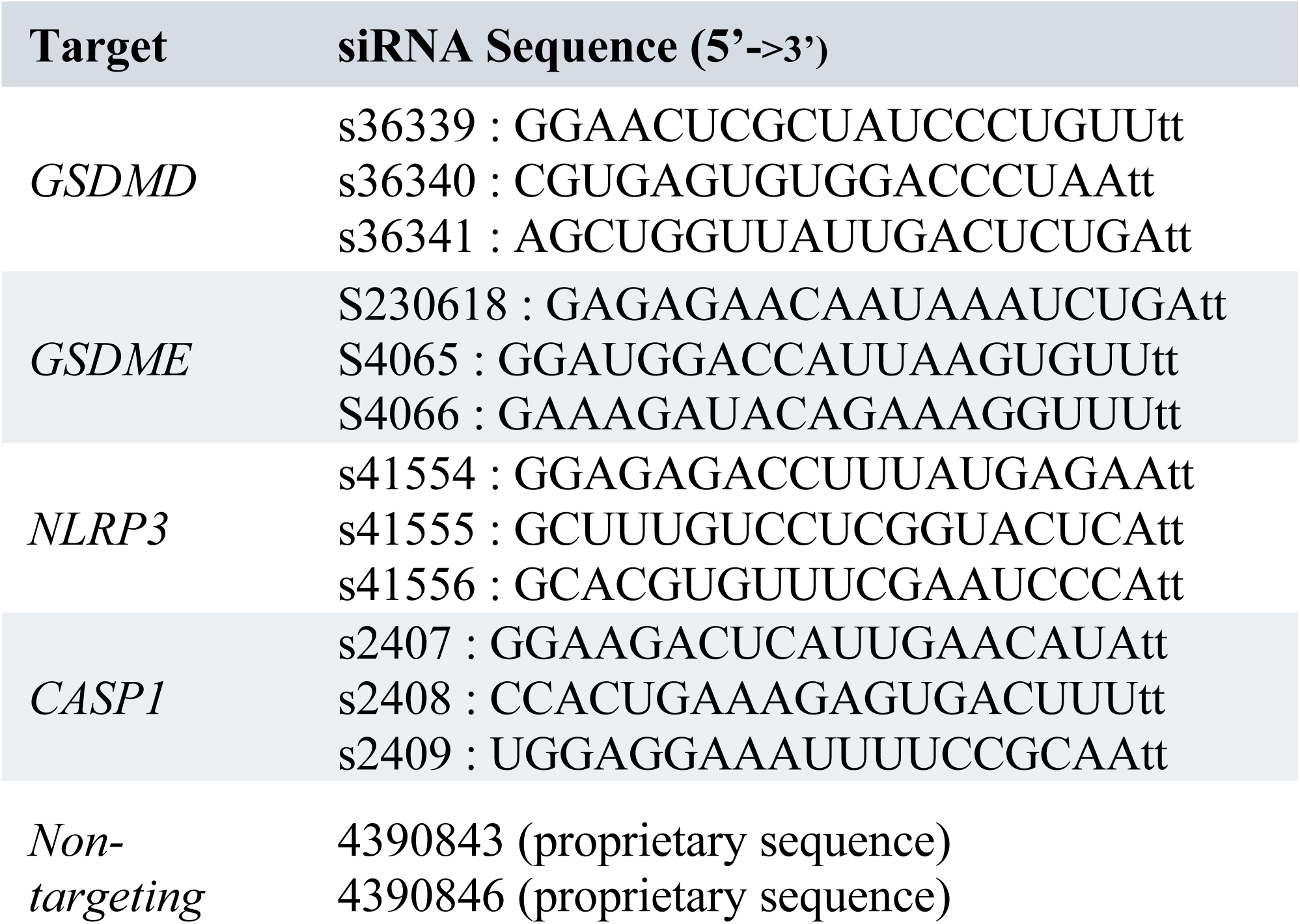

### Generation of GSDMD and GSDME Knock-out JY cells

GSDMD/E double knock-out JY cells were generated by electroporation of ribonucleoprotein complex sgRNA/Cas9 in JY cells using the Neon transfection system (Thermo Fisher). Briefly, 1× 10^6^ JY cells were resuspended in 10 μL of R buffer, mixed with 240 ng of sgRNA for GSDMD, 240 ng of sgRNA for GSDME and 2 µg of HiFi Cas9 endonuclease creating a functional editing RNP complex (ordered from IDT) and electroporated using a 10 μL electroporation pipette tip with two 30 ms pulses at a voltage of 1100 V. Cells were then dispensed directly into a 6-well plate containing 3 mL of R10 cell medium without antibiotic and cultured for 3–5 days before assessing bulk KO efficiency by qPCR. To generate negative control cell lines, JY cells were transfected using non-targeting sgRNA and HiFi Cas9 endonuclease under identical conditions.

### Potassium Efflux Assays

To assess the relative intracellular concentration of K^+^ upon target cell perforation, JY cells were non-pulsed or pulsed with 10μM antigenic peptide (CMV p65) for 2hr at 37° and 5% CO2, washed, and loaded with 2.5 μM Ion Potassium Green-2 AM (Abcam, ab142806) for 30min. Stained JY cells were then conjugated with antigen-specific CTLs at the indicated effector:target ratios for 5 and 15min, stained with viability dye (eFluor 450) and subjected to analysis by flow cytometry. Samples were acquired using the MACSQuant Analyzer 10 or MACS Quant Analyzer VYB flow cytometers (Miltenyi Biotec) and analyzed using FlowJo 10.

As a positive control, JY cells were plated in 96-well U-bottom plates at a density of 2.5 x 10^4^ cells/well and treated with the K^+^ ionphore nigericin (100μM) at time points ranging from 1min to 2hr, before being stained with IPG-2 AM and viability dye and analyzed as above.

### Exogenous Perforin Experiments

JY target cells were plated in 96-well U-bottom plates at a density of 2.5 x 10^4^ cells/well in perforin binding buffer to facilitate perforin phospholipid binding (20mM HEPES, 150nM NaCl, 2.5mM CaCl2). Recombinant human perforin-1 (AA 22-555, Enzo Life Science ENZ-PRT313-0200) was added at the indicated concentrations for a duration of 15min, after which cells were spun, washed, and re-suspended for analysis by flow cytometry. In experiments with excess extracellular K^+^, the perforin binding buffer above was replaced with buffer containing KCl (20mM HEPES, 150nM NaCl, 2.5mM CaCl2).

To assess intracellular K^+^ concentration, JY cells were plated in 96-well U-bottom plates at a density of 2.5 x 10^4^ cells/well in perforin binding buffer and stained with the fluorescent K^+^ probe, Ion Potassium Green-2 AM for 30min as described above. Recombinant perforin was added at a concentration of 4 or 8μg/mL for 5 and 15min, after which cells were spun, washed and analyzed by flow cytometry.

To assess caspase-1 activation, JY cells were plated in 96-well U-bottom plates at a density of 2.5 x 10^4^ cells/well in perforin binding buffer and pre-loaded with far-red caspase-1 activity-dependent probe (ImmunoChemsitry, #9122). Recombinant perforin was added at a concentration of 8μg/mL for 30min, after which cells were spun, washed, and fixed with the provided fixative (ImmunoChemistry #636), prior to analysis by flow cytometry.

For Annexin V/PI assays, JY cells were plated in 96-well U-bottom plates at a density of 2.5 x 10^4^ cells/well in perforin binding buffer and recombinant perforin added at a concentration of 8μg/mL for 60min, after which cells were washed and subjected to Annexin V/PI staining as described above (BD Pharmingen FITC Annexin V Apoptosis Detection Kit I, #AB_2869082) before analysis by flow cytometry.

### Real-time quantitative PCR

Isolation of RNA was performed using the Quick-RNA mini Prep kit (ZymoResearch) and RNA concentration was assessed with the NanoDrop 1000 system (Thermo Scientific). cDNA was synthesized using the SuperScript™ IV VILO™ (SSIV VILO) Master Mix (Invitrogen).

The gene expression of *GSDMA-E*, *CASP1* and *NLRP3* was evaluated by real-time quantitative (RTqPCR) using TaqMan gene expression assays (Applied Biosystems) according to manufacturer’s recommendations, using a StepOne Real-Time PCR System (Applied Biosysrems). All reactions were performed in triplicates and relative gene expression levels were evaluated using the comparative CT (threshold cycle) method (2-deltaCT). GAPDH was used as endogenous controls for normalization.

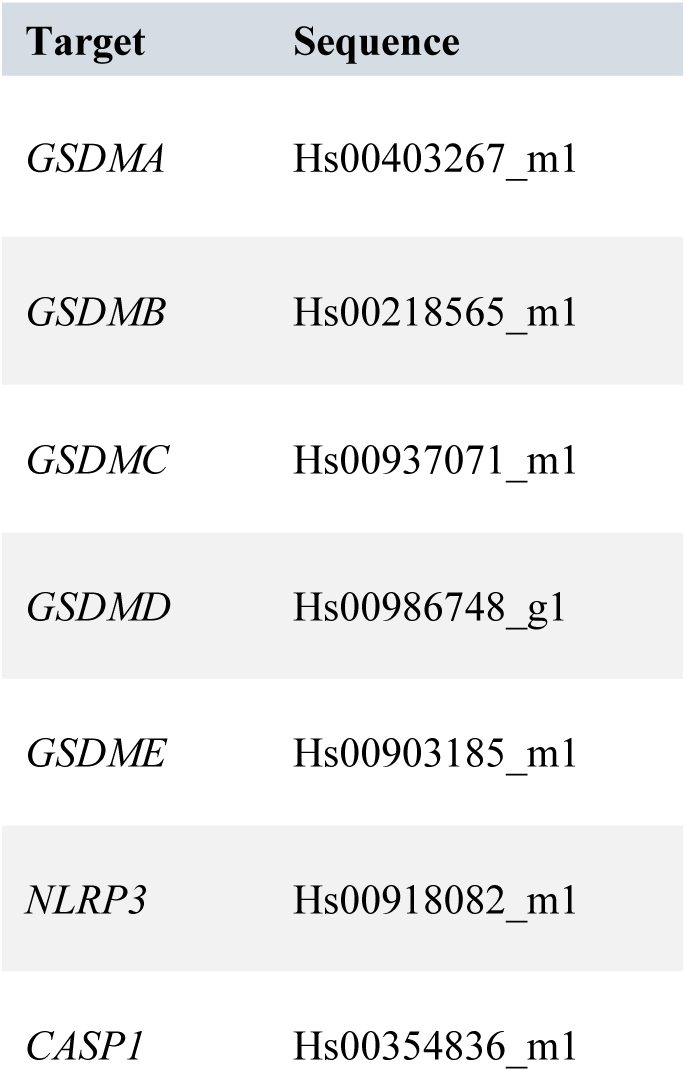
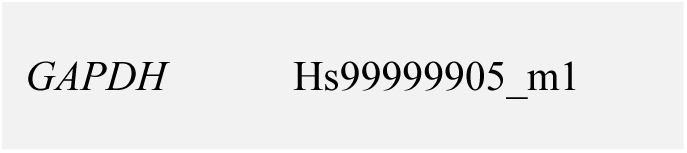

### Immunoblot

*Sample collection:* To assess baseline expression of pyroptosis proteins (GSDMD, GSDME, NLRP3 and caspase-1), 2.5 x 10^6^ JY, D10, or MDA-MB231 cells were washed in cold PBS and resuspended in RIPA buffer with cOmplete^TM^ protease inhibitor cocktail (Sigma-Aldrich).

*Protein extraction and immunoblot:* After 15min on ice with agitation every 5min, extracts were centrifuged to pellet DNA and the supernatant was collected. Proteins were quantified by Pierce BCA protein assay kit (Thermo Fisher Scientific) according to manufacturer’s instructions. Proteins were then denatured in Laemli buffer (5min at 95° with agitation). 15μg of protein was loaded for each sample in precast 4-15% Mini-Protean TGX Stain-Free 30μl 10-well polyacrylamide gels (Bio-Rad #456803) along with Kaleidoscop protein ladder (Bio-Rad). Gels were run and blotted onto a nitrocellulose membrane and blocked with 3% non-fat dry milk in Tris-buffered saline with Tween (TBST) for 1hr. Membrane was incubated with the indicated primary antibodies overnight at 4°C with agitation, followed by 1hr incubation with the corresponding HRP-tagged secondary antibodies. Blots were developed using ECL Clarity (Max) detection reagent (Bio-Rad) and imaged using ChemiDoc MP System (Bio-Rad).

### Primary Antibodies

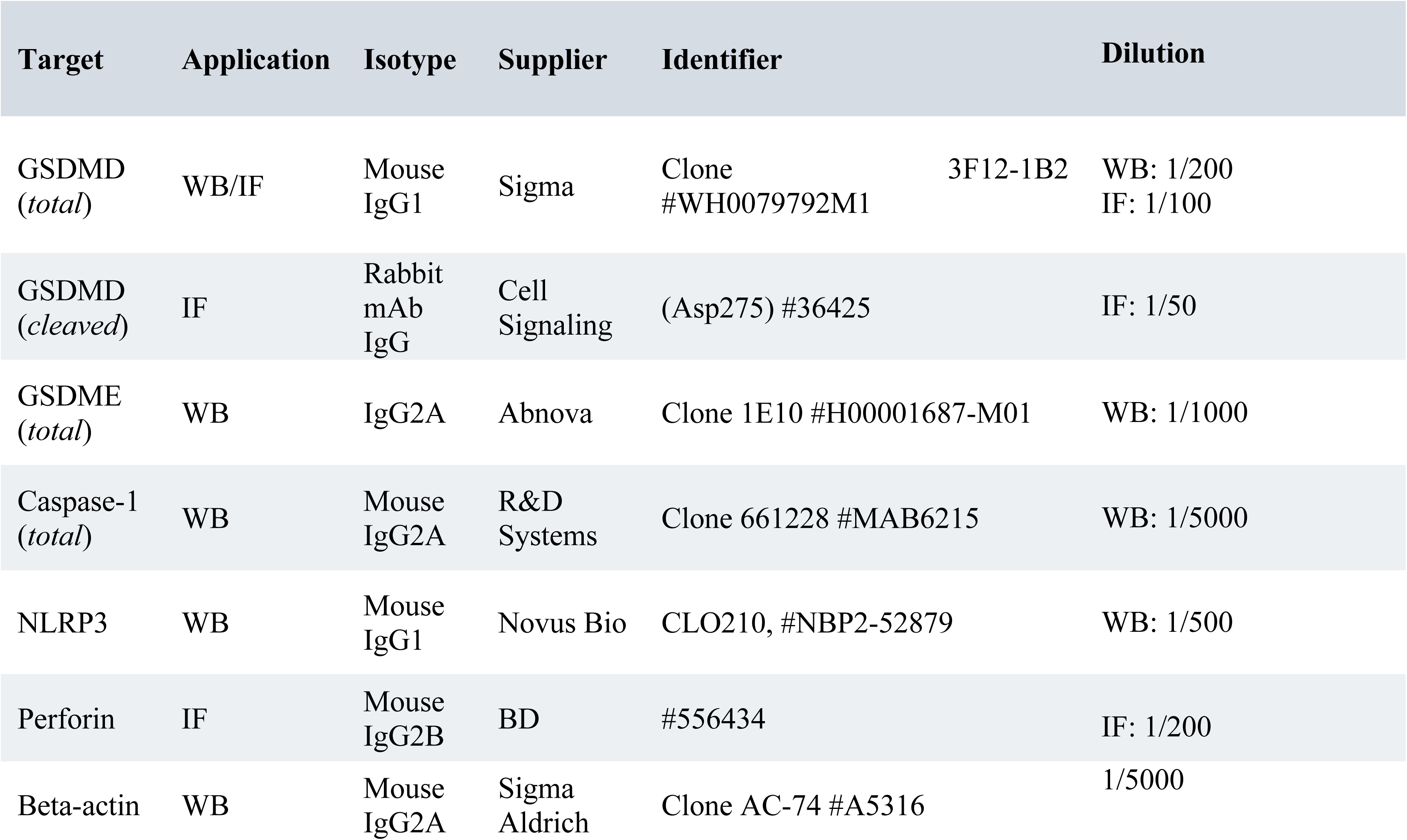

Secondary Antibodies

| Name | Application | Supplier | Identifier | Concentration |
| --- | --- | --- | --- | --- |
| Goat anti-Rabbit HRP | WB | Cell<br>Signaling` | 7074S | 1/2500 |
| Goat anti-Mouse IgG HRP | WB | Cell Signaling | 7076S | 1/2500 |
| Goat anti-rabbit Alexa Fluor™<br>Plus 488 | IF | Invitrogen | A32731 | 1/100 |
| Goat anti-Mouse IgG2b Alexa<br>Fluor™ 555 | IF | Invitrogen | A21147 | 1/100 |
| Goat anti-Mouse IgG1 Alexa<br>Fluor™ 647 | IF | Invitrogen | A21240 | 1/100 |

### Immunofluorescence

JY target cells were either non-pulsed or pulsed with 10μM antigenic peptide (CMV p65) for 2hr at 37 °C and 5% CO2 in RPMI 5% FCS/HEPES, washed three times, and conjugated at an effector:target ratio of 1:1 with antigen-specific human CTLs using 1 min centrifugation at 455g. Conjugates were incubated for 5 or 15 min, gently disrupted, and subsequently seeded on Poly-L-Lysine-coated slides. Cells were fixed with 3% paraformaldehyde at 37 °C for 10min. Cells were then permeabilized with 0.1% saponin (in PBS/3%BSA/HEPES), and stained with the indicated primary antibodies overnight at RT followed by isotype-matched secondary antibodies for 1hr at RT. Samples were washed, mounted in 90% glycerol-PBS containing 2.5% DABCO (Sigma) and examined using a LSM 880 (Zeiss) confocal microscope using a 63x objective. 3D images (using the z-stack function) were acquired at 1μm intervals.

To assess GSDMD expression in target cells, cells were manually outlined and the MFI of total and cleaved GSDMD within the target cell measured (using the Measure function of ImageJ) on 5 different planes of the *z*-stack (separated by 1μm) at the height of the IS and subsequently averaged. To ensure consistency in the quantification, only target cells in the pre-bubbling stage were included in the analysis, and cells with highly irregular bubbling protrusions or ruptured membranes were excluded.

To assess whether total and cleaved GSDMD were expressed evenly throughout the target cell or enriched proximal to the synapse, each target cell was divided into two halves in the x-y plane (synaptic and distal) and MFI quantified separately for each half of the cell. This analysis was repeated in both the cleaved and total GSDMD channels across 5 sequential planes of the z-stack at the height of the IS and the average ratio of synaptic:distal signal was calculated for each cell. According to this analysis, a synaptic:distal ratio greater than 1 indicates synaptic enrichment; equal to 1 indicates even distribution of signal throughout the cell, and less than 1 indicates distal enrichment. For this analysis, the mean synaptic:distal ratio for each channel was compared to a hypothetical value of 1 (indicating even intracellular distribution) using a one-sample *t*-test.

### Time Lapse Microscopy

For drug treatment experiments, JY cells were preloaded with CellMask^TM^ Deep Red plasma membrane stain (1µg/mL) (Invitrogen, #C10046) for 30 min at 37°C with 5% CO2, washed, and then seeded in RPMI 1640 medium with 10% FCS at 2 x 10^5^ - 3 x 10^5^ cells per well on poly-D-lysine coated 8-well chambered slides (Ibidi) for 5 min before imaging. Propidium iodide was added at a concentration of 6.6µg/mL. Drugs were added at the indicated concentrations: nigericin (100µM), staurosporine (6.7 µM), or solvent control. Timelaspe micrsocopy was performed within a temperature-controlled chamber using multi position acquisition (3 positions) overnight on a spinning-disk microscope. All time-lapse microscope experiments were acquired on a spinning disk microscope equipped with APO lbda 60X NA 1.4 objective. The temperature controlled chamber was maintained at 37°C and a constant 5% CO2 concentration. A Prime 95B Scientific sCMOS Camera (Photometrics) was used for acquisition. Image acquisition was controlled by MetaMorph Software (Molecular Devices 7.10.5.476) and by Modular V2.0 GATACA software.

To analyze the morphological features (cross-sectional area and aspect ratio) of dying target cells over the duration of the imaging period, target cell masks were generated using ImageJ software (version 1.53o), using the following semi-automated image processing pipeline: first, images were split into channels. Then, the variance filter was applied to the transmission channel (radius 5.0) followed by “Threshold” and “Transform to Binary”. Next, the “Erode” binary function option was applied and manually cross-checked against the transmission channel. Finally, the “Fill Holes” function was applied and where possible, the “Add Mask” function was utilized to generate the outline. The “Watershed” function was applied as the final step and manual correction was performed as needed on a slice-by-slice basis to remove debris and artifacts. “Analyze Particle” was utilized to draw outlines after manual correction, after which a macro was used to generate the black background and white foreground, and to label and save the mask area. Thus, a cell mask was generated for each cell at every time-point throughout the imaging period, from which the cross-sectional area and aspect ratio could be calculated and plotted over time.

To visualize the morphological features of JY cells dying upon CTL attack, JY cells were pulsed with 10µM peptide, preloaded with Cell Mask DeepRed plasma membrane stain (1µg/mL) (Invitrogen), washed and seeded at 2.5 x 10^4^ cells per wells on poly-D-lysine coated 16 wells-chambered slides (Ibidi). CTL clones were preloaded with Fluo-8-AM (0.25ng/µL) (AAT Bioquest) and SPY650 Tubulin (2X) (Spirochrome) or Tubulin Tracker Deep Red (2.56µg/µL) (Invitrogen) for 30min, washed and added at 4 x 10^4^ to 5 x 10^5^ cells per wells. Propidium iodide (6.6µg/mL) was added immediately before the acquisition. In these experiments, cells were kept in RPMI 1640 with 5% FCS medium for the imaging period.

For the analysis, the three channels were split (transmission, CellMask, Fluo-8-AM). Then, the variance filter was applied to the transmission channel (radius 5.0) and a binary option applied (manually cross-checked against the transmission image). In order to remove the CTL from analysis, the “Create Mask” function was applied to the Fluo-8-AM channel and subtracted from the selection (either the Brightfield/variance or CellMask Channel) using Image Calculator. Finally, the “Fill Holes” and “Watershed” functions were applied and adjusted manually as required. “Analyze Particle” was utilized as a final step described above.

### Exposure of Targets to Purified SMAPs

10, 000 JY cells/well were treated for 4 hours with exogenous SMAPs purified from NK-92 cell supernatant as described (https://doi.org/10.1101/2025.11.10.687675). Briefly, particles were concentrated using100 kDa Amicon spin filters, soluble proteins were removed using Captocore columns, SMAPs were purified by S1000 size exclusion chromatography (https://doi.org/10.1101/2025.11.10.687675). After 4hr, killing modality was assessed using an Annexin V/PI assay (BD Pharmingen FITC Annexin V Apoptosis Detection Kit I, #AB_2869082) as described above. Samples were acquired using the MACSQuant Analyzer 10 flow cytometer (Miltenyi Biotec) and analyzed using FlowJo 10.

### Material Availability

Materials are available from the authors upon request, subject to the appropriate Material Transfer Agreements and constrained by the relevant ethical guidelines. All commercial reagents and materials are available from the indicated sources.

### Statistical Analysis

Statistical analyses were performed using GraphPad Prism 10.1.2. All tests were 2-sided unless otherwise indicated. Data distributions were assumed to follow a normal distribution unless otherwise indicated. Multiple comparisons were performed when more than two groups were being compared, and the specific tests utilized for multiple comparisons have been indicated throughout the manuscript.

## Acknowledgements

We thank the members of the European Research Council ATTACK consortium for their discussion and Dr. Eric Espinosa and Dr. Cosima Baldari for their helpful critiques of this manuscript. We thank the flow cytometry and microscopy core facilities of the INSERM UMR 1037 (CRCT). Parts of the figures (e.g. organelles, membranes) were drawn by using pictures from Servier Medical Art, which is licensed under a Creative Commons Attribution 3.0 Unported License. This research in this manuscript has received funding from the European Research Council (ERC) under the European Union’s Horizon 2020 Research and Innovation Programme (Grant agreement No. Syn- 951329). The research was also supported by INSERM institutional funding, from the Fondation Toulouse Cancer Santé and from the Oncopole Claudius-Regaud. Funders had no role in the preparation of this manuscript.

## Author contributions

BM designed the research, performed experiments, analyzed results, and wrote the paper. MPP, FR, LD, EC, AB, and SM performed experiments, analyzed data, and edited the paper. AJ, MW, CS, LC provided reagents, MLD discussed results and provided research insights, SV supervised the research, designed the project, and wrote the paper.

## Data Availability

All original data generated for this study will be deposited in figshare repository server (https://figshare.com/) with a unique identifier listed here upon acceptance of this article.

## List of Supplementary Movies

|  |  |
| --- | --- |
| **Supplementary Movie 1** | ***Target cell death upon CTL attack resembles pyroptosis and is characterized by rapid erratic membrane bubbling, formation of pyroptotic bodies, loss of membrane integrity***<br>JY target cells were pulsed with 10μM antigenic peptide, and conjugated at an E:T ratio of 1:1 with antigen-specific human CTLs pre-loaded with Fluo-8-AM (*green*) and SPY650 Tubulin (*magenta*) in the presence of 6.6μg/mL propidium iodide (*red*). Conjugate formation and cell death morphology was assessed using spinning disk time lapse microscopy. |
| **Supplementary Movie 2** | ***Target cell death upon CTL attack resembles pyroptosis and is characterized by rapid erratic membrane bubbling, formation of pyroptotic bodies, loss of membrane integrity***<br>JY target cells were pulsed with 10μM antigenic peptide, and conjugated at an E:T ratio of 1:1 with antigen-specific human CTLs pre-loaded with Fluo-8-AM (*green*) and SPY650 Tubulin (*magenta*) in the presence of 6.6μg/mL propidium iodide (*red*). Conjugate formation and cell death morphology was assessed using spinning disk time lapse microscopy. |
| **Supplementary Movie 3** | ***Example of pyroptotic cell death in nigericin-treated cell, characterized by erratic plasma membrane bubbling, cell swelling, and membrane rupture***<br>JY cells were treated with nigericin (100μM), in the presence of 6.6μg/mL propidium iodide (*red*) and imaged using spinning disk confocal microscopy. |
| **Supplementary Movie 4** | ***Example of pyroptotic cell death in nigericin-treated cell, characterized by erratic plasma membrane bubbling, cell swelling, and membrane rupture***<br>JY cells were treated with nigericin (100μM), in the presence of 6.6μg/mL propidium iodide (*red*) and imaged using spinning disk confocal microscopy. |
| **Supplementary Movie 5** | ***Example of pyroptotic cell death in nigericin-treated cell, characterized by erratic plasma membrane bubbling, cell swelling, and membrane rupture***<br>JY cells were treated with nigericin (100μM), in the presence of 6.6μg/mL propidium iodide (*red*) and imaged using spinning disk confocal microscopy. |
| **Supplementary Movie 6** | ***Example of apoptotic cell death in staurosporine-treated cell***<br>JY cells were treated with staurosporine (6.7μM) in the presence of 6.6μg/mL propidium iodide (*red*) and imaged using spinning disk confocal microscopy. |
| **Supplementary Movie 7** | ***Example of apoptotic cell death in staurosporine-treated cell followed by secondary necrosis***<br>JY cells were treated with staurosporine (6.7μM) in the presence of 6.6μg/mL propidium iodide (*red*) and imaged using spinning disk confocal microscopy. |
| **Supplementary Movie 8** | ***Example of apoptotic cell death in staurosporine-treated cell***<br>JY cells were treated with staurosporine (6.7μM) in the presence of 6.6μg/mL propidium iodide (*red*) and imaged using spinning disk confocal microscopy. |
| **Supplementary Movie 9** | ***Example of apoptotic cell death in staurosporine-treated cell***<br>JY cells were treated with staurosporine (6.7μM) in the presence of 6.6μg/mL propidium iodide (*red*) and imaged using spinning disk confocal microscopy. |
| **Supplementary Movie 10** | ***Non-lethal interactions between non-pulsed JY cells and CTLs***<br>Non-pulsed JY target cells were loaded with CellMask DeepRed (*magenta*) and conjugated at an E:T ratio of 1:1 with antigen-specific human CTLs pre-loaded with Fluo-8-AM (*green*) and Tubulin Tracker Deep Red (*magenta*) for 60min and monitored by spinning disk confocal microscopy. |
| **Supplementary Movie 11** | ***Target cell death upon CTL attack resembles pyroptosis and is characterized by rapid erratic membrane bubbling, formation of pyroptotic bodies, loss of membrane integrity***<br>JY target cells were loaded with CellMask DeepRed (*magenta*), pulsed with 10μM antigenic peptide, and conjugated at an E:T ratio of 1:1 with antigen-specific human CTLs pre-loaded with Fluo-8-AM (*green*) and Tubulin Tracker Deep Red (*magenta*) for 60min and monitored by spinning disk confocal microscopy. |
| **Supplementary Movie 12** | ***Target cell death upon CTL attack resembles pyroptosis and is characterized by rapid erratic membrane bubbling, formation of pyroptotic bodies, loss of membrane integrity (additional example)***<br>JY target cells were loaded with CellMask DeepRed (*magenta*), pulsed with 10μM antigenic peptide, and conjugated at an E:T ratio of 1:1 with antigen-specific human CTLs pre-loaded with Fluo-8-AM (*green*) and Tubulin Tracker Deep Red (*magenta*) for 60min and monitored by spinning disk confocal microscopy. |

## REFERENCES

1 Cassioli, C. & Baldari, C. T. The Expanding Arsenal of Cytotoxic T Cells. Front Immunol 13, 883010, doi:10.3389/fimmu.2022.883010 (2022).

2 McKenzie, B. & Valitutti, S. Resisting T cell attack: tumor-cell-intrinsic defense and reparation mechanisms. Trends Cancer 9, 198–211, doi:10.1016/j.trecan.2022.12.003 (2023).

3 McKenzie, B., Khazen, R. & Valitutti, S. Greek Fire, Poison Arrows, and Scorpion Bombs: How Tumor Cells Defend Against the Siege Weapons of Cytotoxic T Lymphocytes. Front Immunol 13, 894306, doi:10.3389/fimmu.2022.894306 (2022).

4 Balint, S. et al. Supramolecular attack particles are autonomous killing entities released from cytotoxic T cells. Science 368, 897–901, doi:10.1126/science.aay9207 (2020).

5. Jainarayanan, A. et al. Isolation of functional supramolecular attack particles (SMAPs). *bioRxiv*, 2025.2011.2010.687675, doi:10.1101/2025.11.10.687675 (2025).

6 Lopez, J. A. et al. Perforin forms transient pores on the target cell plasma membrane to facilitate rapid access of granzymes during killer cell attack. Blood 121, 2659–2668, doi:10.1182/blood-2012-07-446146 (2013).

7 Filali, L. et al. Ultrarapid lytic granule release from CTLs activates Ca(2+)-dependent synaptic resistance pathways in melanoma cells. Sci Adv 8, eabk3234, doi:10.1126/sciadv.abk3234 (2022).

8 Khazen, R. et al. Melanoma cell lysosome secretory burst neutralizes the CTL-mediated cytotoxicity at the lytic synapse. Nat Commun 7, 10823, doi:10.1038/ncomms10823 (2016).

9 Ritter, A. T. et al. ESCRT-mediated membrane repair protects tumor-derived cells against T cell attack. Science 376, 377–382, doi:10.1126/science.abl3855 (2022).

10 Keefe, D. et al. Perforin triggers a plasma membrane-repair response that facilitates CTL induction of apoptosis. Immunity 23, 249–262, doi:10.1016/j.immuni.2005.08.001 (2005).

11 Weigelin, B. & Friedl, P. T cell-mediated additive cytotoxicity - death by multiple bullets. Trends Cancer 8, 980–987, doi:10.1016/j.trecan.2022.07.007 (2022).

12 Weigelin, B. et al. Cytotoxic T cells are able to efficiently eliminate cancer cells by additive cytotoxicity. Nat Commun 12, 5217, doi:10.1038/s41467-021-25282-3 (2021).

13 Voskoboinik, I., Whisstock, J. C. & Trapani, J. A. Perforin and granzymes: function, dysfunction and human pathology. Nat Rev Immunol 15, 388–400, doi:10.1038/nri3839 (2015).

14 Ivanova, M. E. et al. The pore conformation of lymphocyte perforin. Sci Adv 8, eabk3147, doi:10.1126/sciadv.abk3147 (2022).

15 Stewart, S. E. et al. The perforin pore facilitates the delivery of cationic cargos. J Biol Chem 289, 9172–9181, doi:10.1074/jbc.M113.544890 (2014).

16 Shewell, L. K. et al. All major cholesterol-dependent cytolysins use glycans as cellular receptors. Sci Adv 6, eaaz4926, doi:10.1126/sciadv.aaz4926 (2020).

17 Shinkai, Y., Takio, K. & Okumura, K. Homology of perforin to the ninth component of complement (C9). Nature 334, 525–527, doi:10.1038/334525a0 (1988).

18 Dal Peraro, M. & van der Goot, F. G. Pore-forming toxins: ancient, but never really out of fashion. Nat Rev Microbiol 14, 77–92, doi:10.1038/nrmicro.2015.3 (2016).

19 Margheritis, E., Kappelhoff, S. & Cosentino, K. Pore-Forming Proteins: From Pore Assembly to Structure by Quantitative Single-Molecule Imaging. Int J Mol Sci 24, doi:10.3390/ijms24054528 (2023).

20 Tapia-Abellan, A. et al. Sensing low intracellular potassium by NLRP3 results in a stable open structure that promotes inflammasome activation. Sci Adv 7, eabf4468, doi:10.1126/sciadv.abf4468 (2021).

21 Swanson, K. V., Deng, M. & Ting, J. P. The NLRP3 inflammasome: molecular activation and regulation to therapeutics. Nat Rev Immunol 19, 477–489, doi:10.1038/s41577-019-0165-0 (2019).

22 Munoz-Planillo, R. et al. K(+) efflux is the common trigger of NLRP3 inflammasome activation by bacterial toxins and particulate matter. Immunity 38, 1142–1153, doi:10.1016/j.immuni.2013.05.016 (2013).

23 Gurcel, L., Abrami, L., Girardin, S., Tschopp, J. & van der Goot, F. G. Caspase-1 activation of lipid metabolic pathways in response to bacterial pore-forming toxins promotes cell survival. Cell 126, 1135–1145, doi:10.1016/j.cell.2006.07.033 (2006).

24 Higa, N. et al. Vibrio parahaemolyticus effector proteins suppress inflammasome activation by interfering with host autophagy signaling. PLoS Pathog 9, e1003142, doi:10.1371/journal.ppat.1003142 (2013).

25 Craven, R. R. et al. Staphylococcus aureus alpha-hemolysin activates the NLRP3-inflammasome in human and mouse monocytic cells. PLoS One 4, e7446, doi:10.1371/journal.pone.0007446 (2009).

26 Holzinger, D. et al. Staphylococcus aureus Panton-Valentine leukocidin induces an inflammatory response in human phagocytes via the NLRP3 inflammasome. J Leukoc Biol 92, 1069–1081, doi:10.1189/jlb.0112014 (2012).

27 Liang, H., Wang, B., Wang, J., Ma, B. & Zhang, W. Pyolysin of Trueperella pyogenes Induces Pyroptosis and IL-1beta Release in Murine Macrophages Through Potassium/NLRP3/Caspase-1/Gasdermin D Pathway. Front Immunol 13, 832458, doi:10.3389/fimmu.2022.832458 (2022).

28 De Schutter, E. et al. Punching Holes in Cellular Membranes: Biology and Evolution of Gasdermins. Trends Cell Biol 31, 500–513, doi:10.1016/j.tcb.2021.03.004 (2021).

29 Wang, Y. et al. Chemotherapy drugs induce pyroptosis through caspase-3 cleavage of a gasdermin. Nature 547, 99–103, doi:10.1038/nature22393 (2017).

30 Kayagaki, N. et al. Caspase-11 cleaves gasdermin D for non-canonical inflammasome signalling. Nature 526, 666–671, doi:10.1038/nature15541 (2015).

31 Liu, X. et al. Inflammasome-activated gasdermin D causes pyroptosis by forming membrane pores. Nature 535, 153–158, doi:10.1038/nature18629 (2016).

32 Mulvihill, E. et al. Mechanism of membrane pore formation by human gasdermin-D. EMBO J 37, doi:10.15252/embj.201798321 (2018).

33 Ding, J. et al. Pore-forming activity and structural autoinhibition of the gasdermin family. Nature 535, 111–116, doi:10.1038/nature18590 (2016).

34 DiPeso, L., Ji, D. X., Vance, R. E. & Price, J. V. Cell death and cell lysis are separable events during pyroptosis. Cell Death Discov 3, 17070, doi:10.1038/cddiscovery.2017.70 (2017).

35 Devant, P. & Kagan, J. C. Molecular mechanisms of gasdermin D pore-forming activity. Nat Immunol 24, 1064–1075, doi:10.1038/s41590-023-01526-w (2023).

36 Xia, S. et al. Gasdermin D pore structure reveals preferential release of mature interleukin-1. Nature 593, 607–611, doi:10.1038/s41586-021-03478-3 (2021).

37 Yu, P. et al. Pyroptosis: mechanisms and diseases. Signal Transduct Target Ther 6, 128, doi:10.1038/s41392-021-00507-5 (2021).

38 Ruhl, S. & Broz, P. Regulation of Lytic and Non-Lytic Functions of Gasdermin Pores. J Mol Biol 434, 167246, doi:10.1016/j.jmb.2021.167246 (2022).

39 Li, Z. et al. Enhancing Gasdermin-induced tumor pyroptosis through preventing ESCRT-dependent cell membrane repair augments antitumor immune response. Nat Commun 13, 6321, doi:10.1038/s41467-022-34036-8 (2022).

40 Ruhl, S. et al. ESCRT-dependent membrane repair negatively regulates pyroptosis downstream of GSDMD activation. Science 362, 956–960, doi:10.1126/science.aar7607 (2018).

41 Nozaki, K. et al. Caspase-7 activates ASM to repair gasdermin and perforin pores. Nature 606, 960–967, doi:10.1038/s41586-022-04825-8 (2022).

42 Shkarina, K. et al. Optogenetic activators of apoptosis, necroptosis, and pyroptosis. J Cell Biol 221, doi:10.1083/jcb.202109038 (2022).

43 Kovacs, S. B. & Miao, E. A. Gasdermins: Effectors of Pyroptosis. Trends Cell Biol 27, 673–684, doi:10.1016/j.tcb.2017.05.005 (2017).

44 Costigan, A., Hollville, E. & Martin, S. J. Discriminating Between Apoptosis, Necrosis, Necroptosis, and Ferroptosis by Microscopy and Flow Cytometry. Curr Protoc 3, e951, doi:10.1002/cpz1.951 (2023).

45 Zhang, Y., Chen, X., Gueydan, C. & Han, J. Plasma membrane changes during programmed cell deaths. Cell Res 28, 9–21, doi:10.1038/cr.2017.133 (2018).

46 Chen, X. et al. Pyroptosis is driven by non-selective gasdermin-D pore and its morphology is different from MLKL channel-mediated necroptosis. Cell Res 26, 1007–1020, doi:10.1038/cr.2016.100 (2016).

47 Liu, X., Xia, S., Zhang, Z., Wu, H. & Lieberman, J. Channelling inflammation: gasdermins in physiology and disease. Nat Rev Drug Discov 20, 384–405, doi:10.1038/s41573-021-00154-z (2021).

48 Zychlinsky, A., Zheng, L. M., Liu, C. C. & Young, J. D. Cytolytic lymphocytes induce both apoptosis and necrosis in target cells. J Immunol 146, 393–400 (1991).

49 Blanchard, D. K. et al. Role of extracellular adenosine triphosphate in the cytotoxic T-lymphocyte-mediated lysis of antigen presenting cells. Blood 85, 3173–3182 (1995).

50 Waterhouse, N. J. et al. Cytotoxic T lymphocyte-induced killing in the absence of granzymes A and B is unique and distinct from both apoptosis and perforin-dependent lysis. J Cell Biol 173, 133–144, doi:10.1083/jcb.200510072 (2006).

51 Jaime-Sanchez, P. et al. Cell death induced by cytotoxic CD8(+) T cells is immunogenic and primes caspase-3-dependent spread immunity against endogenous tumor antigens. J Immunother Cancer 8, doi:10.1136/jitc-2020-000528 (2020).

52 Martinez-Lostao, L., Anel, A. & Pardo, J. How Do Cytotoxic Lymphocytes Kill Cancer Cells? Clin Cancer Res 21, 5047–5056, doi:10.1158/1078-0432.CCR-15-0685 (2015).

53 Zhou, Z. et al. Granzyme A from cytotoxic lymphocytes cleaves GSDMB to trigger pyroptosis in target cells. Science 368, doi:10.1126/science.aaz7548 (2020).

54 Zhang, Z. et al. Gasdermin E suppresses tumour growth by activating anti-tumour immunity. Nature 579, 415–420, doi:10.1038/s41586-020-2071-9 (2020).

55 Liu, Y. et al. Gasdermin E-mediated target cell pyroptosis by CAR T cells triggers cytokine release syndrome. Sci Immunol 5, doi:10.1126/sciimmunol.aax7969 (2020).

56 Galluzzi, L. et al. Molecular mechanisms of cell death: recommendations of the Nomenclature Committee on Cell Death 2018. Cell Death Differ 25, 486–541, doi:10.1038/s41418-017-0012-4 (2018).

57 Reiss, C. S. et al. Development and characterization of allospecific long-term human cytolytic T-cell lines. Proc Natl Acad Sci U S A 77, 5432–5436, doi:10.1073/pnas.77.9.5432 (1980).

58 Caramalho, I., Faroudi, M., Padovan, E., Muller, S. & Valitutti, S. Visualizing CTL/melanoma cell interactions: multiple hits must be delivered for tumour cell annihilation. J Cell Mol Med 13, 3834–3846, doi:10.1111/j.1582-4934.2008.00586.x (2009).

59 Faroudi, M. et al. Lytic versus stimulatory synapse in cytotoxic T lymphocyte/target cell interaction: manifestation of a dual activation threshold. Proc Natl Acad Sci U S A 100, 14145–14150, doi:10.1073/pnas.2334336100 (2003).

60 Nagahama, M. et al. The p38 MAPK and JNK pathways protect host cells against Clostridium perfringens beta-toxin. Infect Immun 81, 3703–3708, doi:10.1128/IAI.00579-13 (2013).

61 Rana, P. S. et al. Calibration and characterization of intracellular Asante Potassium Green probes, APG-2 and APG-4. Anal Biochem **567**, 8-13, doi:10.1016/j.ab.2018.11.024 (2019).

62 Wong, F. et al. Cytoplasmic condensation induced by membrane damage is associated with antibiotic lethality. Nat Commun 12, 2321, doi:10.1038/s41467-021-22485-6 (2021).

63 Camilli, G. et al. beta-Glucan-induced reprogramming of human macrophages inhibits NLRP3 inflammasome activation in cryopyrinopathies. J Clin Invest 130, 4561–4573, doi:10.1172/JCI134778 (2020).

64 Ren, G. et al. ABRO1 promotes NLRP3 inflammasome activation through regulation of NLRP3 deubiquitination. EMBO J 38, doi:10.15252/embj.2018100376 (2019).

65 Perregaux, D. & Gabel, C. A. Interleukin-1 beta maturation and release in response to ATP and nigericin. Evidence that potassium depletion mediated by these agents is a necessary and common feature of their activity. J Biol Chem 269, 15195–15203 (1994).

66 Walev, I., Reske, K., Palmer, M., Valeva, A. & Bhakdi, S. Potassium-inhibited processing of IL-1 beta in human monocytes. EMBO J 14, 1607–1614, doi:10.1002/j.1460-2075.1995.tb07149.x (1995).

67 Russo, H. M. et al. Active Caspase-1 Induces Plasma Membrane Pores That Precede Pyroptotic Lysis and Are Blocked by Lanthanides. J Immunol 197, 1353–1367, doi:10.4049/jimmunol.1600699 (2016).

68 Rozario, P. et al. Mechanistic basis for potassium efflux-driven activation of the human NLRP1 inflammasome. Proc Natl Acad Sci U S A 121, e2309579121, doi:10.1073/pnas.2309579121 (2024).

69 Sun, Y. & Guo, Y. Expression of Caspase-1 in breast cancer tissues and its effects on cell proliferation, apoptosis and invasion. Oncol Lett 15, 6431–6435, doi:10.3892/ol.2018.8176 (2018).

70 Zheng, W. et al. Caspase-1-dependent spatiality in triple-negative breast cancer and response to immunotherapy. Nat Commun 15, 8514, doi:10.1038/s41467-024-52553-6 (2024).

71 Duhalde Vega, M., et al. PD-1/PD-L1 blockade abrogates a dysfunctional innate-adaptive immune axis in critical beta-coronavirus disease. Sci Adv 8, eabn6545, doi:10.1126/sciadv.abn6545 (2022).

72 Clark, K. M. et al. Chemical inhibition of DPP9 sensitizes the CARD8 inflammasome in HIV-1-infected cells. Nat Chem Biol 19, 431–439, doi:10.1038/s41589-022-01182-5 (2023).

73 Yu, J. et al. Inflammasome activation leads to Caspase-1-dependent mitochondrial damage and block of mitophagy. Proc Natl Acad Sci U S A 111, 15514–15519, doi:10.1073/pnas.1414859111 (2014).

74 Boucher, D. et al. Caspase-1 self-cleavage is an intrinsic mechanism to terminate inflammasome activity. J Exp Med 215, 827–840, doi:10.1084/jem.20172222 (2018).

75 Mitchell, P. S., Sandstrom, A. & Vance, R. E. The NLRP1 inflammasome: new mechanistic insights and unresolved mysteries. Curr Opin Immunol 60, 37–45, doi:10.1016/j.coi.2019.04.015 (2019).

76 Yang, X. et al. Granzyme B mimics apical caspases. Description of a unified pathway for trans-activation of executioner caspase-3 and -7. J Biol Chem 273, 34278-34283, doi:10.1074/jbc.273.51.34278 (1998).

77 Adrain, C., Murphy, B. M. & Martin, S. J. Molecular ordering of the caspase activation cascade initiated by the cytotoxic T lymphocyte/natural killer (CTL/NK) protease granzyme B. J Biol Chem 280, 4663–4673, doi:10.1074/jbc.M410915200 (2005).

78 Nakazato, K. & Hatano, Y. Monensin-mediated antiport of Na+ and H+ across liposome membrane. Biochim Biophys Acta 1064, 103–110, doi:10.1016/0005-2736(91)90416-6 (1991).

79 Kataoka, T. et al. Concanamycin A, a powerful tool for characterization and estimation of contribution of perforin- and Fas-based lytic pathways in cell-mediated cytotoxicity. J Immunol 156, 3678–3686 (1996).

80 Miao, R. et al. Gasdermin D permeabilization of mitochondrial inner and outer membranes accelerates and enhances pyroptosis. Immunity 56, 2523–2541 e2528, doi:10.1016/j.immuni.2023.10.004 (2023).

81 Khazen, R., Muller, S., Lafouresse, F., Valitutti, S. & Cussat-Blanc, S. Sequential adjustment of cytotoxic T lymphocyte densities improves efficacy in controlling tumor growth. Sci Rep 9, 12308, doi:10.1038/s41598-019-48711-2 (2019).

82 Yan, H. et al. Cisplatin Induces Pyroptosis via Activation of MEG3/NLRP3/caspase-1/GSDMD Pathway in Triple-Negative Breast Cancer. Int J Biol Sci 17, 2606–2621, doi:10.7150/ijbs.60292 (2021).

83 Zhang, Z. et al. Caspase-3-mediated GSDME induced Pyroptosis in breast cancer cells through the ROS/JNK signalling pathway. J Cell Mol Med 25, 8159–8168, doi:10.1111/jcmm.16574 (2021).

84 McKenzie, B. A. et al. Activation of the executioner caspases-3 and -7 promotes microglial pyroptosis in models of multiple sclerosis. J Neuroinflammation 17, 253, doi:10.1186/s12974-020-01902-5 (2020).

85 McKenzie, B. A. et al. Caspase-1 inhibition prevents glial inflammasome activation and pyroptosis in models of multiple sclerosis. Proc Natl Acad Sci U S A 115, E6065–E6074, doi:10.1073/pnas.1722041115 (2018).

86 Megli, C., Morosky, S., Rajasundaram, D. & Coyne, C. B. Inflammasome signaling in human placental trophoblasts regulates immune defense against Listeria monocytogenes infection. J Exp Med 218, doi:10.1084/jem.20200649 (2021).

87 Ma, H., Jeppesen, J. F. & Jaenisch, R. Human T Cells Expressing a CD19 CAR-T Receptor Provide Insights into Mechanisms of Human CD19-Positive beta Cell Destruction. Cell Rep Med 1, 100097, doi:10.1016/j.xcrm.2020.100097 (2020).

88 Mariathasan, S. et al. Cryopyrin activates the inflammasome in response to toxins and ATP. Nature 440, 228–232, doi:10.1038/nature04515 (2006).

89 Flores, J. et al. Caspase-1 inhibition alleviates cognitive impairment and neuropathology in an Alzheimer’s disease mouse model. Nat Commun 9, 3916, doi:10.1038/s41467-018-06449-x (2018).

90 Dhani, S., Zhao, Y. & Zhivotovsky, B. A long way to go: caspase inhibitors in clinical use. Cell Death Dis 12, 949, doi:10.1038/s41419-021-04240-3 (2021).

91 Hassin, D., Garber, O. G., Meiraz, A., Schiffenbauer, Y. S. & Berke, G. Cytotoxic T lymphocyte perforin and Fas ligand working in concert even when Fas ligand lytic action is still not detectable. Immunology 133, 190–196, doi:10.1111/j.1365-2567.2011.03426.x (2011).

92 Meiraz, A., Garber, O. G., Harari, S., Hassin, D. & Berke, G. Switch from perforin-expressing to perforin-deficient CD8(+) T cells accounts for two distinct types of effector cytotoxic T lymphocytes in vivo. Immunology 128, 69–82, doi:10.1111/j.1365-2567.2009.03072.x (2009).

93 Prager, I. et al. NK cells switch from granzyme B to death receptor-mediated cytotoxicity during serial killing. J Exp Med 216, 2113–2127, doi:10.1084/jem.20181454 (2019).

94 Chang, H. F. et al. Identification of distinct cytotoxic granules as the origin of supramolecular attack particles in T lymphocytes. Nat Commun 13, 1029, doi:10.1038/s41467-022-28596-y (2022).

95 Ambrose, A. R., Hazime, K. S., Worboys, J. D., Niembro-Vivanco, O. & Davis, D. M. Synaptic secretion from human natural killer cells is diverse and includes supramolecular attack particles. Proc Natl Acad Sci U S A 117, 23717–23720, doi:10.1073/pnas.2010274117 (2020).

96 Kimman, T. et al. Serpin B9 controls tumor cell killing by CAR T cells. J Immunother Cancer 11, doi:10.1136/jitc-2022-006364 (2023).

97 Jiang, P. et al. Signatures of T cell dysfunction and exclusion predict cancer immunotherapy response. Nat Med 24, 1550–1558, doi:10.1038/s41591-018-0136-1 (2018).

98 Gonzalez, M. R. et al. Pore-forming toxins induce multiple cellular responses promoting survival. Cell Microbiol 13, 1026–1043, doi:10.1111/j.1462-5822.2011.01600.x (2011).

99 Demeersseman, L., et al. Stenocytosis: a mechanism for supramolecular attack particle transfer at the CTL lytic synapse. *bioRxiv*, 2025.2011.2005.686769, doi:10.1101/2025.11.05.686769 (2025).

100 Gong, T., Yang, Y., Jin, T., Jiang, W. & Zhou, R. Orchestration of NLRP3 Inflammasome Activation by Ion Fluxes. Trends Immunol 39, 393–406, doi:10.1016/j.it.2018.01.009 (2018).

101 Barnett, K. C., Li, S., Liang, K. & Ting, J. P. A 360 degrees view of the inflammasome: Mechanisms of activation, cell death, and diseases. Cell 186, 2288–2312, doi:10.1016/j.cell.2023.04.025 (2023).

102 Yao, Y. et al. Antigen-specific CD8(+) T cell feedback activates NLRP3 inflammasome in antigen-presenting cells through perforin. Nat Commun 8, 15402, doi:10.1038/ncomms15402 (2017).

103 Agard, N. J., Maltby, D. & Wells, J. A. Inflammatory stimuli regulate caspase substrate profiles. Mol Cell Proteomics 9, 880–893, doi:10.1074/mcp.M900528-MCP200 (2010).

104 Shao, W., Yeretssian, G., Doiron, K., Hussain, S. N. & Saleh, M. The caspase-1 digestome identifies the glycolysis pathway as a target during infection and septic shock. J Biol Chem 282, 36321–36329, doi:10.1074/jbc.M708182200 (2007).

105 Loveless, R., Bloomquist, R. & Teng, Y. Pyroptosis at the forefront of anticancer immunity. J Exp Clin Cancer Res 40, 264, doi:10.1186/s13046-021-02065-8 (2021).

106 Rogers, C. et al. Gasdermin pores permeabilize mitochondria to augment caspase-3 activation during apoptosis and inflammasome activation. Nat Commun 10, 1689, doi:10.1038/s41467-019-09397-2 (2019).

107 de Vasconcelos, N. M., Van Opdenbosch, N., Van Gorp, H., Parthoens, E. & Lamkanfi, M. Single-cell analysis of pyroptosis dynamics reveals conserved GSDMD-mediated subcellular events that precede plasma membrane rupture. Cell Death Differ 26, 146–161, doi:10.1038/s41418-018-0106-7 (2019).

108 Vasconcelos, Z. et al. Individual Human Cytotoxic T Lymphocytes Exhibit Intraclonal Heterogeneity during Sustained Killing. Cell Rep 11, 1474–1485, doi:10.1016/j.celrep.2015.05.002 (2015).

109 Lemaitre, F., Moreau, H. D., Vedele, L. & Bousso, P. Phenotypic CD8+ T cell diversification occurs before, during, and after the first T cell division. J Immunol 191, 1578–1585, doi:10.4049/jimmunol.1300424 (2013).

110 Kelso, A. et al. The genes for perforin, granzymes A-C and IFN-gamma are differentially expressed in single CD8(+) T cells during primary activation. Int Immunol 14, 605–613, doi:10.1093/intimm/dxf028 (2002).

111 Cortacero, K. et al. Evolutionary design of explainable algorithms for biomedical image segmentation. Nat Commun 14, 7112, doi:10.1038/s41467-023-42664-x (2023).

112 Ramirez-Labrada, A. et al. All About (NK Cell-Mediated) Death in Two Acts and an Unexpected Encore: Initiation, Execution and Activation of Adaptive Immunity. Front Immunol 13, 896228, doi:10.3389/fimmu.2022.896228 (2022).

113 Fisher, R., Pusztai, L. & Swanton, C. Cancer heterogeneity: implications for targeted therapeutics. Br J Cancer 108, 479–485, doi:10.1038/bjc.2012.581 (2013).

114 Hanahan, D. & Weinberg, R. A. The hallmarks of cancer. Cell 100, 57–70, doi:10.1016/s0092-8674(00)81683-9 (2000).

115 Hanahan, D. & Weinberg, R. A. Hallmarks of cancer: the next generation. Cell 144, 646–674, doi:10.1016/j.cell.2011.02.013 (2011).

116 Hanahan, D. Hallmarks of Cancer: New Dimensions. Cancer Discov 12, 31–46, doi:10.1158/2159-8290.CD-21-1059 (2022).

117 Zapatka, M. et al. The landscape of viral associations in human cancers. Nat Genet 52, 320–330, doi:10.1038/s41588-019-0558-9 (2020).

118 Fan, J. X. et al. Epigenetics-Based Tumor Cells Pyroptosis for Enhancing the Immunological Effect of Chemotherapeutic Nanocarriers. Nano Lett 19, 8049–8058, doi:10.1021/acs.nanolett.9b03245 (2019).

119 Erkes, D. A. et al. Mutant BRAF and MEK Inhibitors Regulate the Tumor Immune Microenvironment via Pyroptosis. Cancer Discov 10, 254–269, doi:10.1158/2159-8290.CD-19-0672 (2020).

120 Lu, H. et al. Molecular Targeted Therapies Elicit Concurrent Apoptotic and GSDME-Dependent Pyroptotic Tumor Cell Death. Clin Cancer Res 24, 6066–6077, doi:10.1158/1078-0432.CCR-18-1478 (2018).

121 Liu, Y. G. et al. NLRP3 inflammasome activation mediates radiation-induced pyroptosis in bone marrow-derived macrophages. Cell Death Dis 8, e2579, doi:10.1038/cddis.2016.460 (2017).

122 Tan, G. et al. Radiosensitivity of colorectal cancer and radiation-induced gut damages are regulated by gasdermin E. Cancer Lett 529, 1–10, doi:10.1016/j.canlet.2021.12.034 (2022).

123 Li, H. et al. Pyroptotic cell death: an emerging therapeutic opportunity for radiotherapy. Cell Death Discov 10, 32, doi:10.1038/s41420-024-01802-0 (2024).

124 Wang, Q. et al. A bioorthogonal system reveals antitumour immune function of pyroptosis. Nature 579, 421–426, doi:10.1038/s41586-020-2079-1 (2020).

125 Zhang, Z., Zhang, Y. & Lieberman, J. Lighting a Fire: Can We Harness Pyroptosis to Ignite Antitumor Immunity? Cancer Immunol Res 9, 2–7, doi:10.1158/2326-6066.CIR-20-0525 (2021).

126 Phulphagar, K. et al. Proteomics reveals distinct mechanisms regulating the release of cytokines and alarmins during pyroptosis. Cell Rep 34, 108826, doi:10.1016/j.celrep.2021.108826 (2021).

127 Mellman, I., Chen, D. S., Powles, T. & Turley, S. J. The cancer-immunity cycle: Indication, genotype, and immunotype. Immunity 56, 2188–2205, doi:10.1016/j.immuni.2023.09.011 (2023).

128 Holley, C. L. et al. Pyroptotic cell corpses are crowned with F-actin-rich filopodia that engage CLEC9A signaling in incoming dendritic cells. Nat Immunol, doi:10.1038/s41590-024-02024-3 (2024).

129 Deng, T., Tang, C., Zhang, G. & Wan, X. DAMPs released by pyroptotic cells as major contributors and therapeutic targets for CAR-T-related toxicities. Cell Death Dis 12, 129, doi:10.1038/s41419-021-03428-x (2021).

130 Luri-Rey, C. et al. Cytotoxicity as a form of immunogenic cell death leading to efficient tumor antigen cross-priming. Immunol Rev 321, 143–151, doi:10.1111/imr.13281 (2024).

131 Li, S. et al. Gasdermin D in peripheral myeloid cells drives neuroinflammation in experimental autoimmune encephalomyelitis. J Exp Med 216, 2562–2581, doi:10.1084/jem.20190377 (2019).

132 McKenzie, B. A., Dixit, V. M. & Power, C. Fiery Cell Death: Pyroptosis in the Central Nervous System. Trends Neurosci 43, 55–73, doi:10.1016/j.tins.2019.11.005 (2020).

133 Wu, Y. et al. Cell pyroptosis in health and inflammatory diseases. Cell Death Discov 8, 191, doi:10.1038/s41420-022-00998-3 (2022).

134 Welz, M. et al. Perforin inhibition protects from lethal endothelial damage during fulminant viral hepatitis. Nat Commun 9, 4805, doi:10.1038/s41467-018-07213-x (2018).

