## Supplemental figures for "Perforin-mediated pore formation at the lytic synapse triggers the canonical pyroptotic cell death pathway"

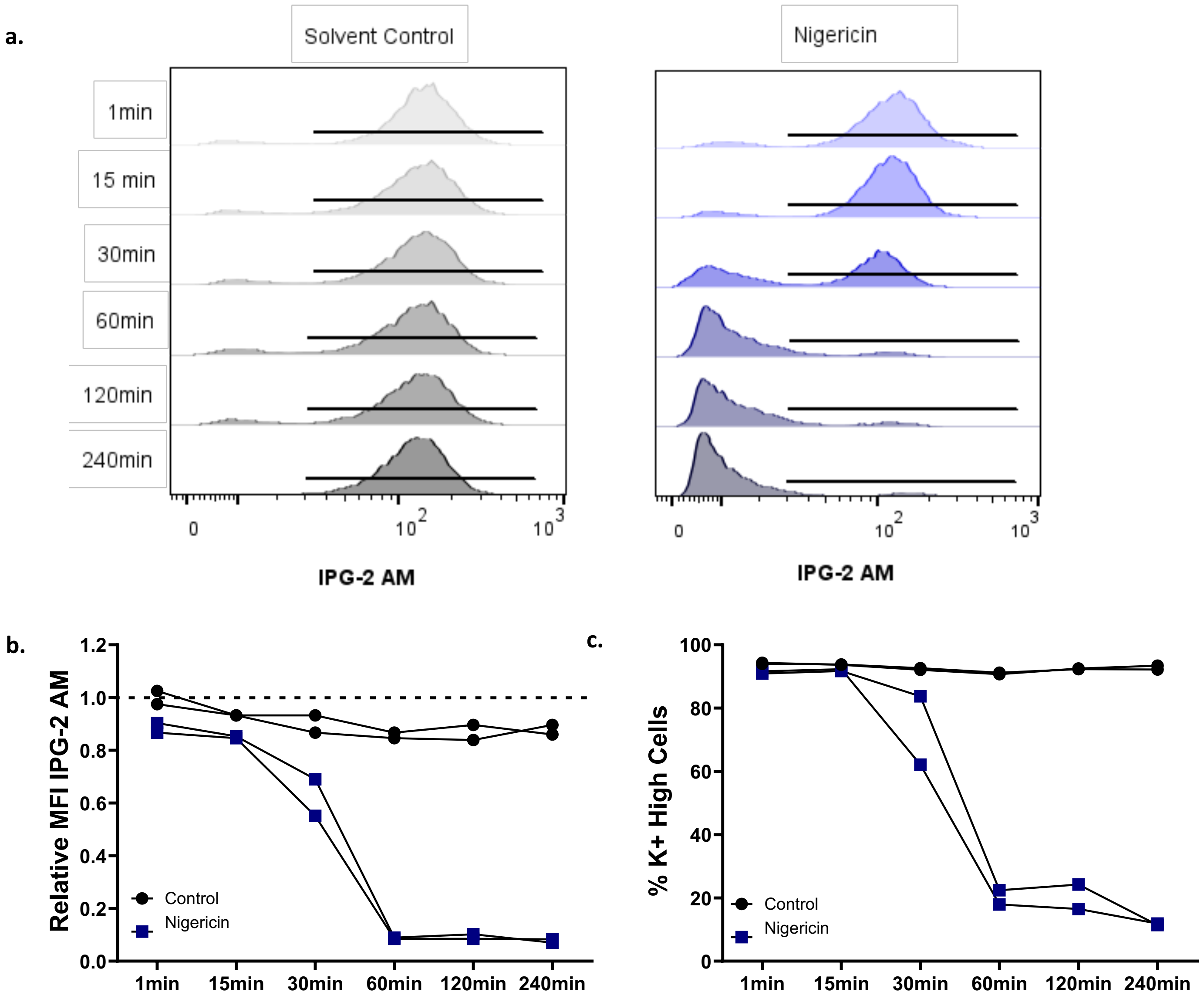


**Supplementary Figure 1| Validation of Ion Potassium Green (IPG)-2 AM probe**

**a-c.** JY cells were treated with the potassium ionophore nigericin (100µM) or solvent control (EtOH) for the indicated times before being stained with IPG-2 AM and analyzed by flow cytometry. **a.** Representative histograms shown for each timepoint (n=2 technical replicates of a single proof-of-concept experiment). Black bar indicates IPG-2 $AM^{bright}$ cells. **b.** Median fluorescence intensity of IPG 2-AM normalized to the 1min solvent control condition (indicated by dotted line). Datapoints represent technical replicates. **c.** Percentage of cells within the IPG-2 $AM^{Bright}$ gate at each time point following exposure to nigericin or solvent control. Datapoints represent technical replicates.

**a.**

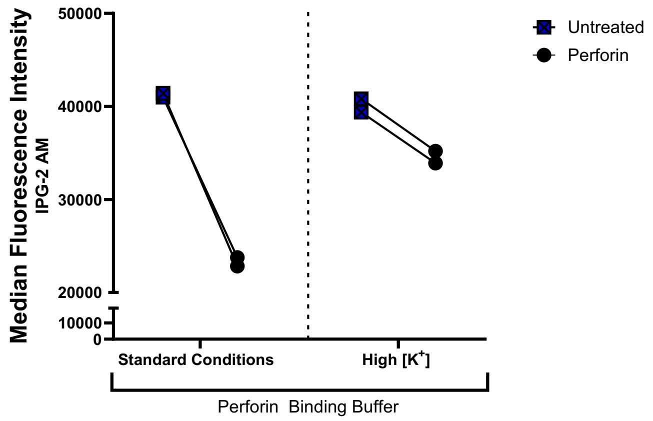


**Supplementary Figure 2| Decrease in IPG-2 AM intensity upon perforin exposure is attributable to $K^+$ efflux**

**a.** JY cells were loaded with the potassium sensor, Ion Potassium Green-2 AM (**IPG-2 AM**) and exposed to 4µg/mL recombinant human perforin for 15min in either standard perforin-binding buffer or high [$K^+$] perforin binding buffer. At the experimental endpoint, cells were collected and IPG-2 AM intensity assessed by flow cytometry. Data shown represent the raw median fluorescence intensity of IPG-2 AM in perforin-treated or untreated cells under each buffer condition. Each point shown on the graph represents a technical replicate from a single experiment, which is representative of n=5 independent experiments.

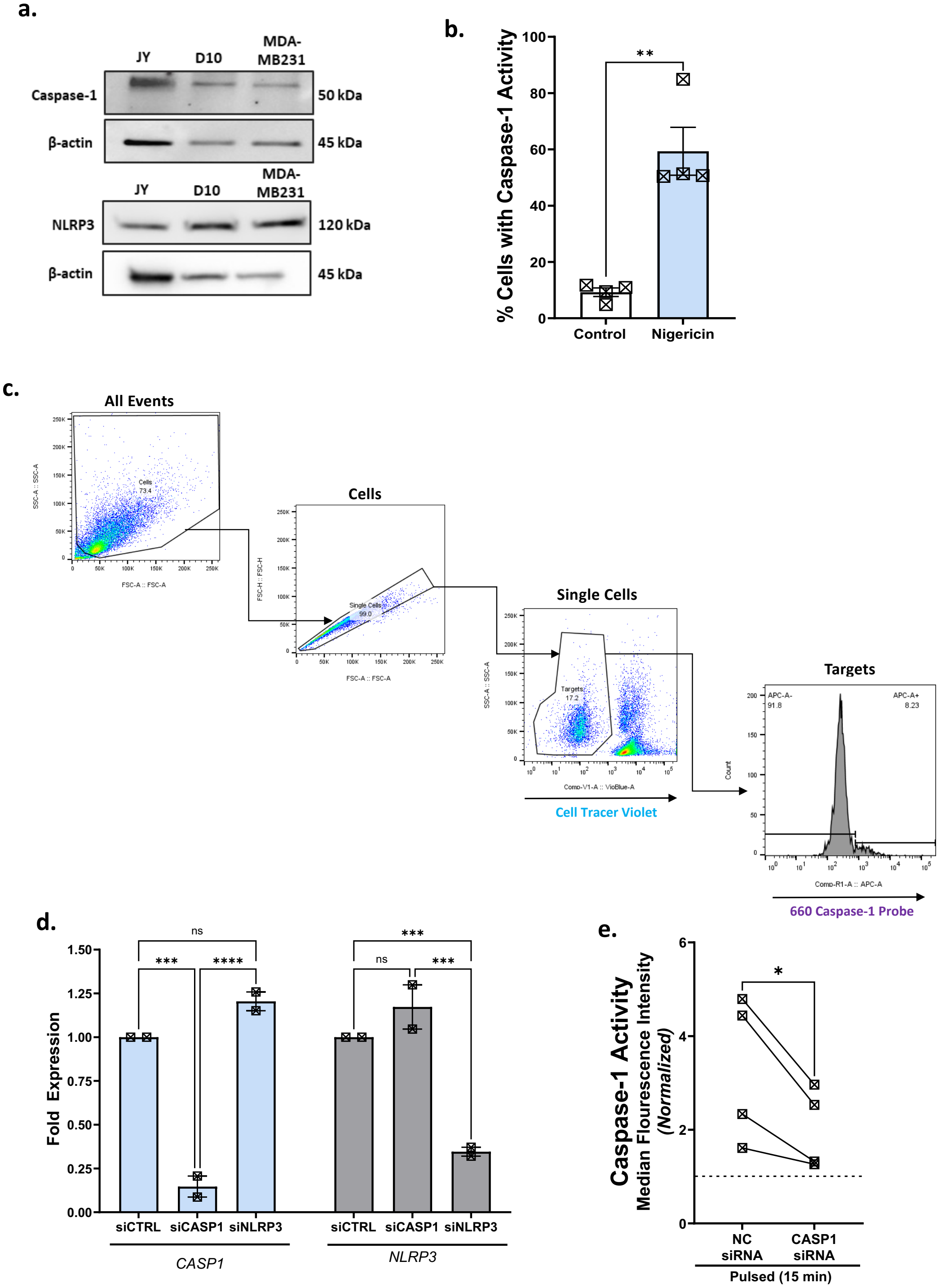


**Supplementary Figure 3| Validation of molecular tools for assessing caspase-1 pathway**

**a.** Immunoblot of JY, D10, and MDA-MB231 target cells at baseline probed for total caspase-1 and NLRP3 protein expression, with β-actin as a loading control. Representative of n=2 immunoblots. **b.** JY cells were treated with 100µM nigericin for 2hr, assayed for caspase-1 activity at the experimental endpoint using the caspase-1 activity-dependent fluorescent probe, and assessed by flow cytometry. Each datapoint represents the average of two technical replicates from an individual experiment, with a total of n=4 independent experiments represented on the graph. Bars represent mean ± SEM. An unpaired *t*-test was utilized to assess statistical significance. **c.** Gating strategy for detecting caspase-1 activity by flow cytometry. Total cells were differentiated from debris on the basis of forward (**FSC-A**) and side-scatter (**SSC-A**). Single cells were selected by comparing FSC-A and FSC-H. Target cells were differentiated from Cell Tracer Violet-stained or Hoescht-stained CTLs based on SSC-A and exclusion of violet dye. Finally, the selected target cells were represented as a histogram with caspase-1 probe intensity on the x-axis. **d.** JY cells were transfected with siRNA targeting either *CASP1* (**siCASP1**) or *NLRP3* (**siNLRP3**) or non-targeting control (**siCTRL**) and harvested for RT-PCR to assess *CASP1* and *NLRP3* expression (data shown is normalized to the non-targeting control; each datapoint represents an independent transfection experiment) **e.** To test the specificity of the activity-dependent probe for caspase-1, JY cells were transfected with non-coding siRNA (**NC siRNA**) or siRNA targeting *CASP1* (**CASP1 siRNA**) before being pulsed with antigenic peptide, conjugated with CTLs for 15min, stained with the caspase-1 activity-dependent probe, and assessed by flow cytometry. Each datapoint represents the average of two technical replicates from an individual experiment, with a total of n=4 independent experiments represented on the graph. A paired *t*-test was utilized to assess statistical significance. * $p<0.05$. *** $p<0.001$, **** $p<0.0001$, ns = not significant.

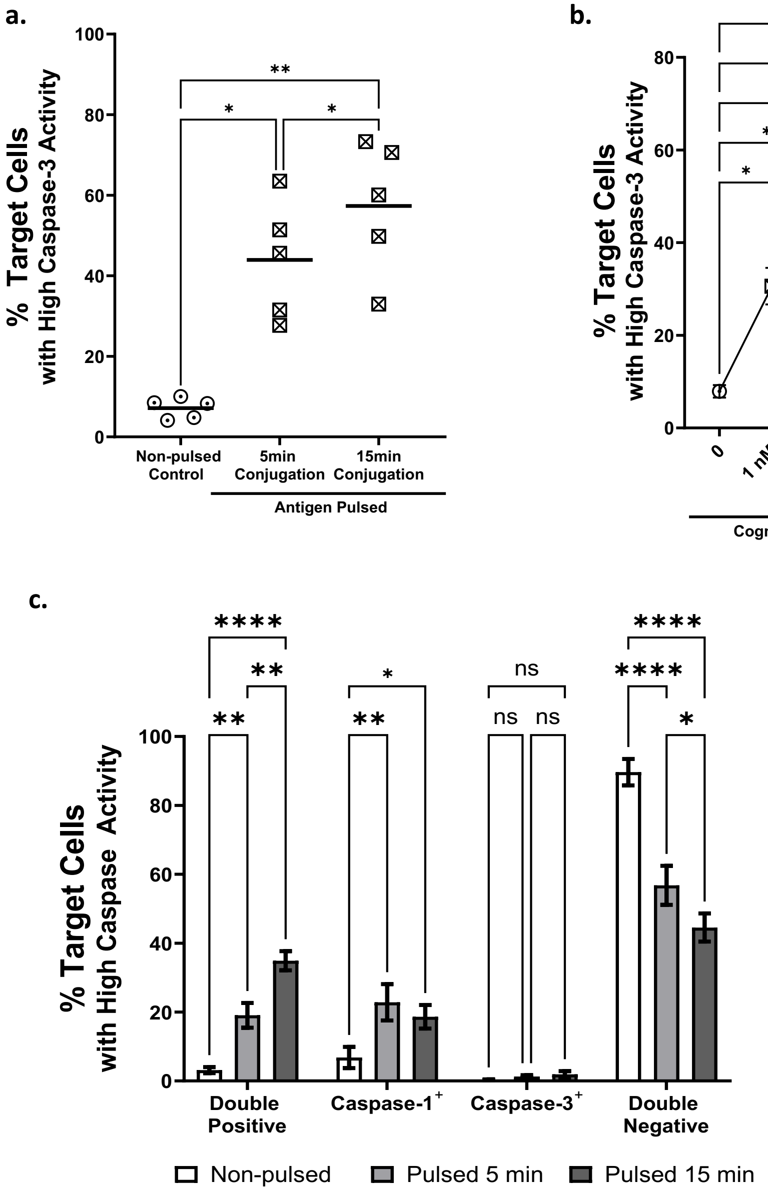


**Supplementary Figure 4| Caspase-3 is activated alongside caspase-1 during CTL attack**

**a.** JY target cells were pulsed with 10µM antigenic peptide and conjugated with CTLs at an E:T ratio of 4:1 for 5 or 15min, before being stained with an activity-dependent fluorescent caspase-3 probe and assessed using flow cytometry. Non-pulsed target cells were conjugated with CTLs under identical conditions for 15min as a control. Each datapoint represents the average of two technical replicates from an individual experiment, with n=5 independent experiments represented on the graph. RM one-way ANOVA with Tukey's test for multiple comparisons was utilized to assess statistical significance. **b.** JY cells were pulsed with the indicated concentration of antigenic peptide at an E:T ratio of 4:1 for 15min and assessed as in **(a).** Non-pulsed target cells were conjugated with CTLs under identical conditions for 15min as a control. Data shown represents mean +/- SEM, n=5 independent experiments. **c.** JY target cells were conjugated with CTLs as in **(a)** and double-stained with activity-dependent caspase-1 and caspase-3 probes before being assessed by flow cytometry. Cells were classified as single-positive, double-positive, or double-negative for each caspase at each timepoint shown. Bars represent mean +/- SEM, with n=7 independent experiments. RM two-way ANOVA with Tukey's test for multiple comparisons was utilized to assess statistical significance. * $p$<0.05. ** $p$<0.01. *** $p$<0.001, **** $p$<0.0001, ns = not significant.

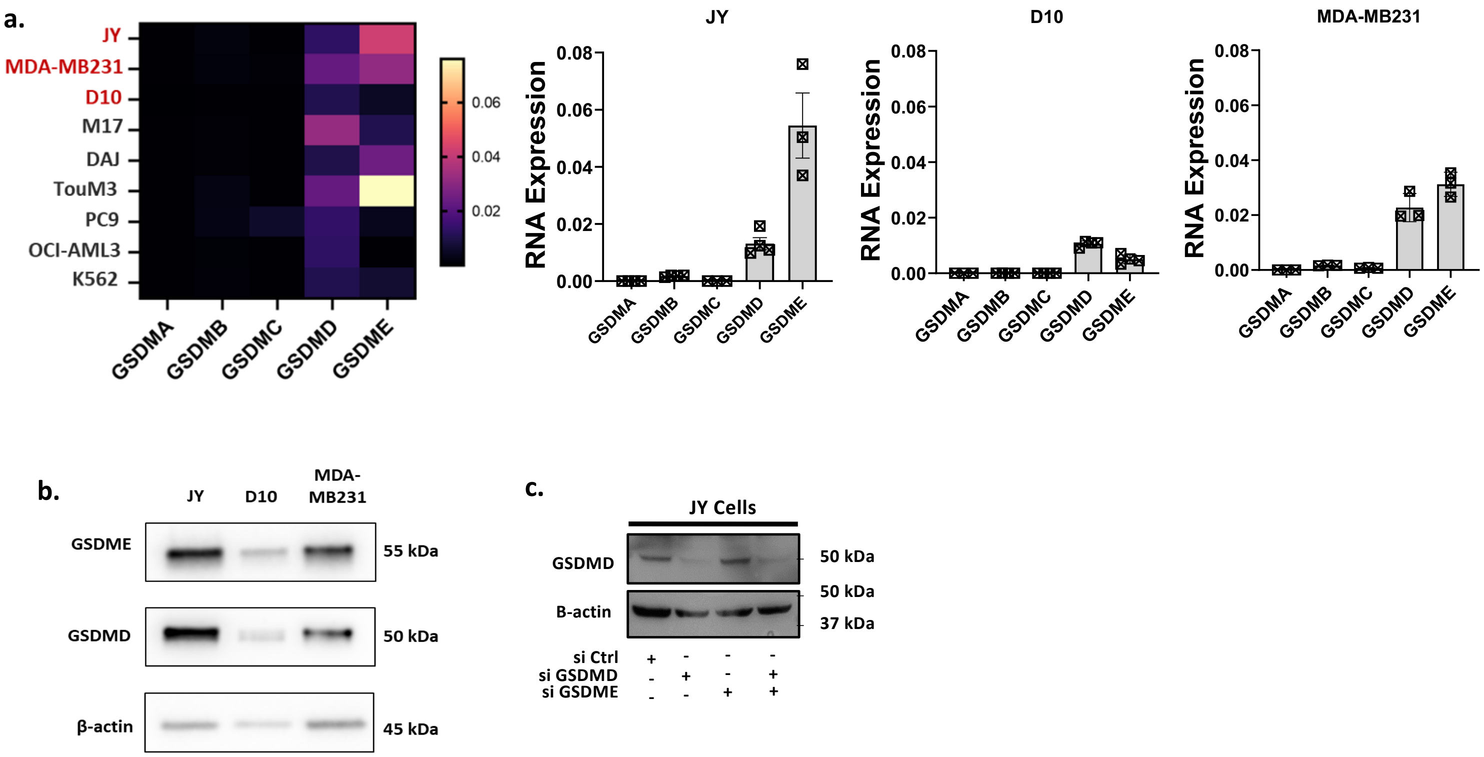


**Supplementary Figure 5| Expression of gasdermins at transcript and protein level in target cells**

**a.** A panel of nine different target cell lines was subjected to RT-PCR to determine baseline transcript expression level of *GSDMA-E*. Heatmap represents average RNA expression level normalized to GAPDH over n=3-4 biological experiments. Bar graphs represent mean ± SEM, with each datapoint representing the results from an individual experiment. **b.** Immunoblots of JY, D10, and MDA-MB231 target cells at baseline probed for full-length GSDMD and GSDME protein expression with β-actin as a loading control. Representative of n=2 immunoblots. **c.** To assess specificity of the GSDMD antibody, JY cells were transfected with non-targeting siRNA (**siCtrl**), or siRNA targeting *GSDMD* (**siGSDMD**), *GSDME* (**siGSDME**), or both for 48hr, harvested, and subject to immunoblot for full-length GSDMD with β-actin as a loading control.

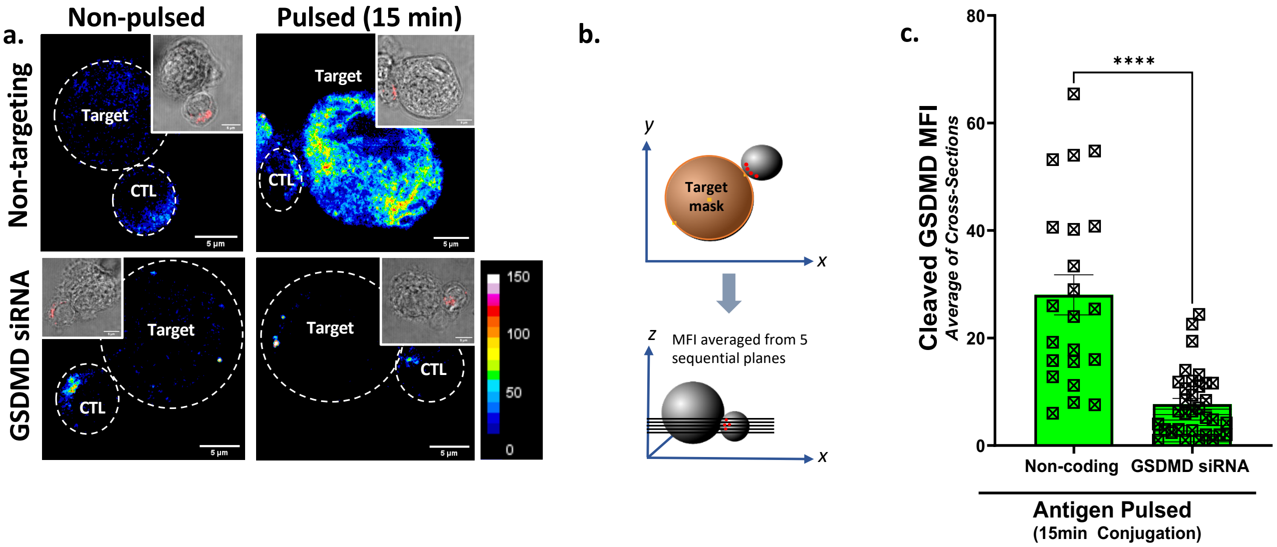


**Supplementary Figure 6| Cleaved GSDMD signal is significantly diminished in GSDMD siRNA-transfected target cells upon CTL attack**

**a.** JY target cells were transfected with non-targeting or *GSDMD*-targeting siRNA before being pulsed (or not) with antigenic peptide and conjugated with antigen-specific CTLs for 15min. Conjugates were fixed and subject to immunostaining for cleaved GSDMD (*pseudocolor*) and perforin (*red*, shown on insets). Images shown are from a single plane of the *z*-stack and are illustrative examples from n=3 independent experiments. **b.** Schematic for the quantification of cleaved GSDMD MFI across five sequential planes of the *z*-stack. The five planes were chosen to correspond to the vertical location of the lytic synapse as indicated by polarized lytic granules. **c.** Mean fluorescence intensity (**MFI**) of cleaved GSDMD within the target cell was measured on 5 different planes of the *z*-stack (separated by 1µm) at the height of the synapse and subsequently averaged. Each datapoint corresponds to one target cell, drawn from n=2 independent experiments. Bars show mean ± SEM. Statistical differences between groups were assessed by unpaired *t*-test. **** $p<0.0001$

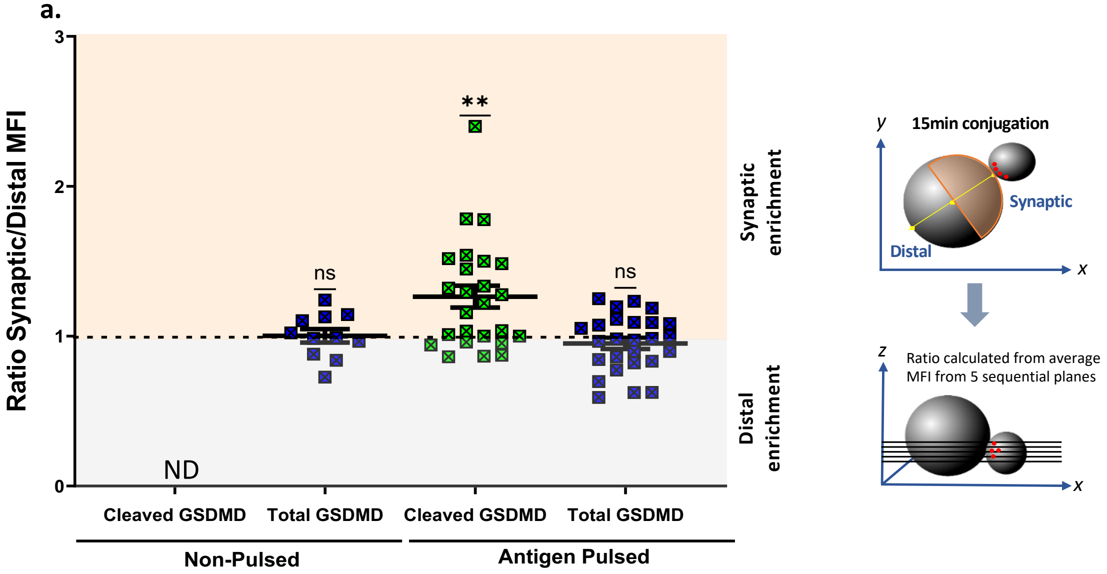


**Supplementary Figure 7│ Cleaved GSDMD is enriched near the synapse after 15min conjugation**

**a.** JY target cells were pulsed with 10µM antigenic peptide and conjugated at an E:T ratio of 1:1 with antigen-specific human CTLs for 15min. Conjugates were fixed and subject to immunostaining for total GSDMD, cleaved GSDMD and perforin (**Figure 3**). **a.** To assess whether cleaved or total GSDMD was enriched proximal to the IS, each target cell was divided into two halves in the x-y plane (synaptic and distal) and MFI quantified separately for each half of the cell. This analysis was repeated across 5 planes of the z-stack and the average ratio of synaptic:distal signal for cleaved and total GSDMD was calculated**.** Ratios higher than 1 indicate synaptic enrichment, lower than 1 indicates distal enrichment, and equal to 1 indicates even distribution throughout the cell. Each datapoint shown represents a single cell. Mean ratio of synaptic:distal MFI for both cleaved and total GSDMD was compared to a hypothetical value of 1 (indicating even distribution of signal throughout the cell) using one-sample *t*-tests. Each datapoint corresponds to one target cell, drawn from n=3 independent experiments. Error bars represent mean +/- SEM. ** $p<0.01$, ns = non-significant, ND = not detectable.

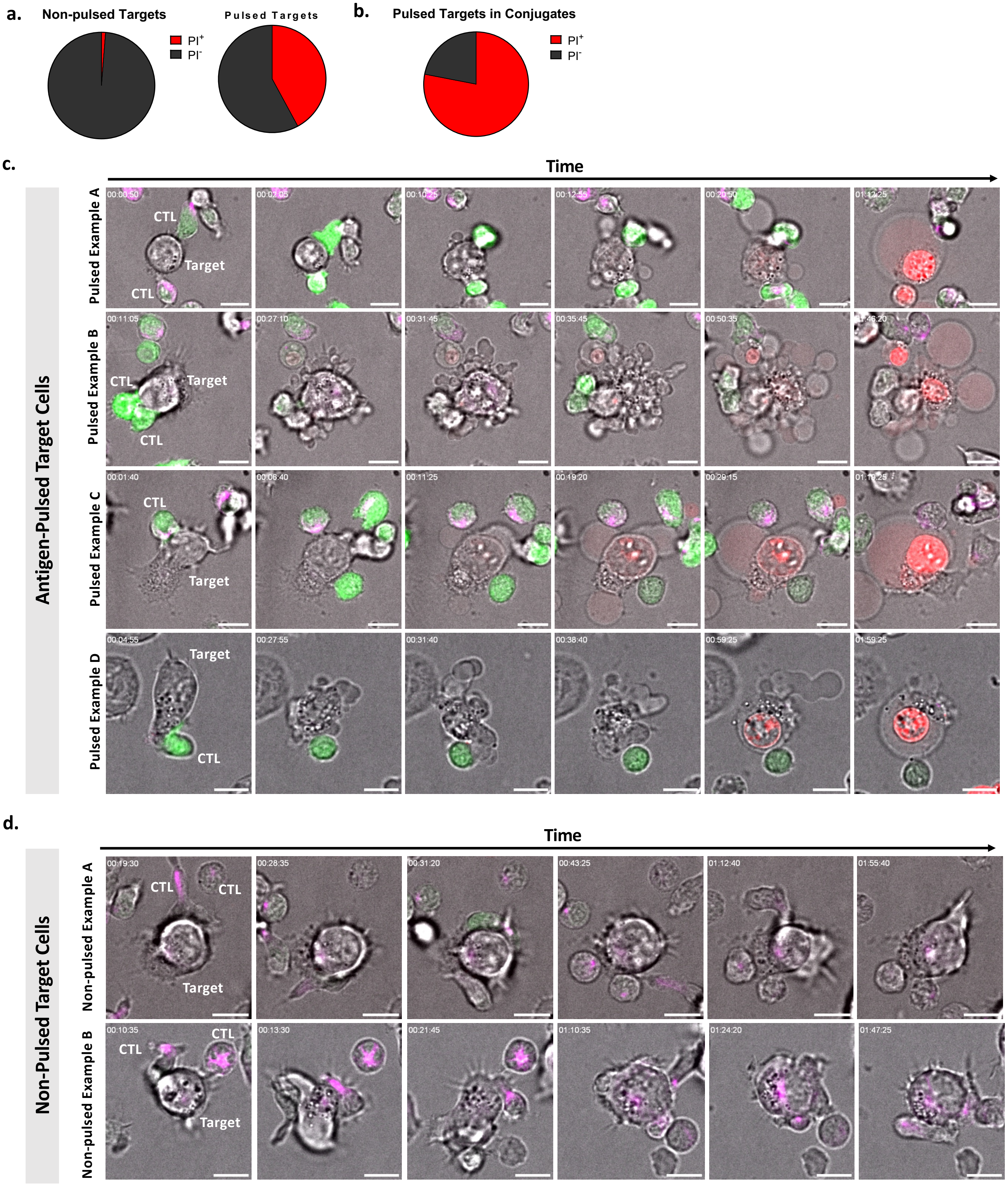


**Supplementary Figure 8| Target cell death upon CTL attack is characterized by violent bubbling and loss of membrane integrity**

**a-d.** JY target cells were pulsed or not with 10µM antigenic peptide, and conjugated at an E:T ratio of 1:1 with antigen-specific human CTLs pre-loaded with Fluo-8-AM (*green*) and SPY650 Tubulin (*magenta*) in the presence of 6.6µg/mL propidium iodide (*red*). Conjugate formation and cell death morphology was assessed using spinning disk time lapse microscopy for 2hr. **a.** Data shown represent the proportion of total JY target cells that were $PI^+$ in non-pulsed (*left*) and pulsed (*right*) conditions after 2hr conjugation with antigen-specific CTLs (n=82 non-pulsed target cells and n=76 pulsed target cells quantified from n=3 independent experiments). **b.** Data shown represent the proportion of pulsed JY target cells in conjugate with CTLs that are $PI^+$ after 2hr conjugation (n=41 target cells in conjugates quantified from n=3 independent experiments). **c.** Snapshots illustrate various stages of cell death including initial CTL contact, synapse formation, pyroptotic body formation, and loss of membrane integrity, indicated by PI entry. Four representative examples from n=3 independent experiments are shown. **d.** Representative images from the non-pulsed condition of the same experiments show transient, non-specific and non-lethal interactions between CTLs and targets, associated with a lack of calcium flux (*green*) PI entry (*red*). Scale bar is 10µm.

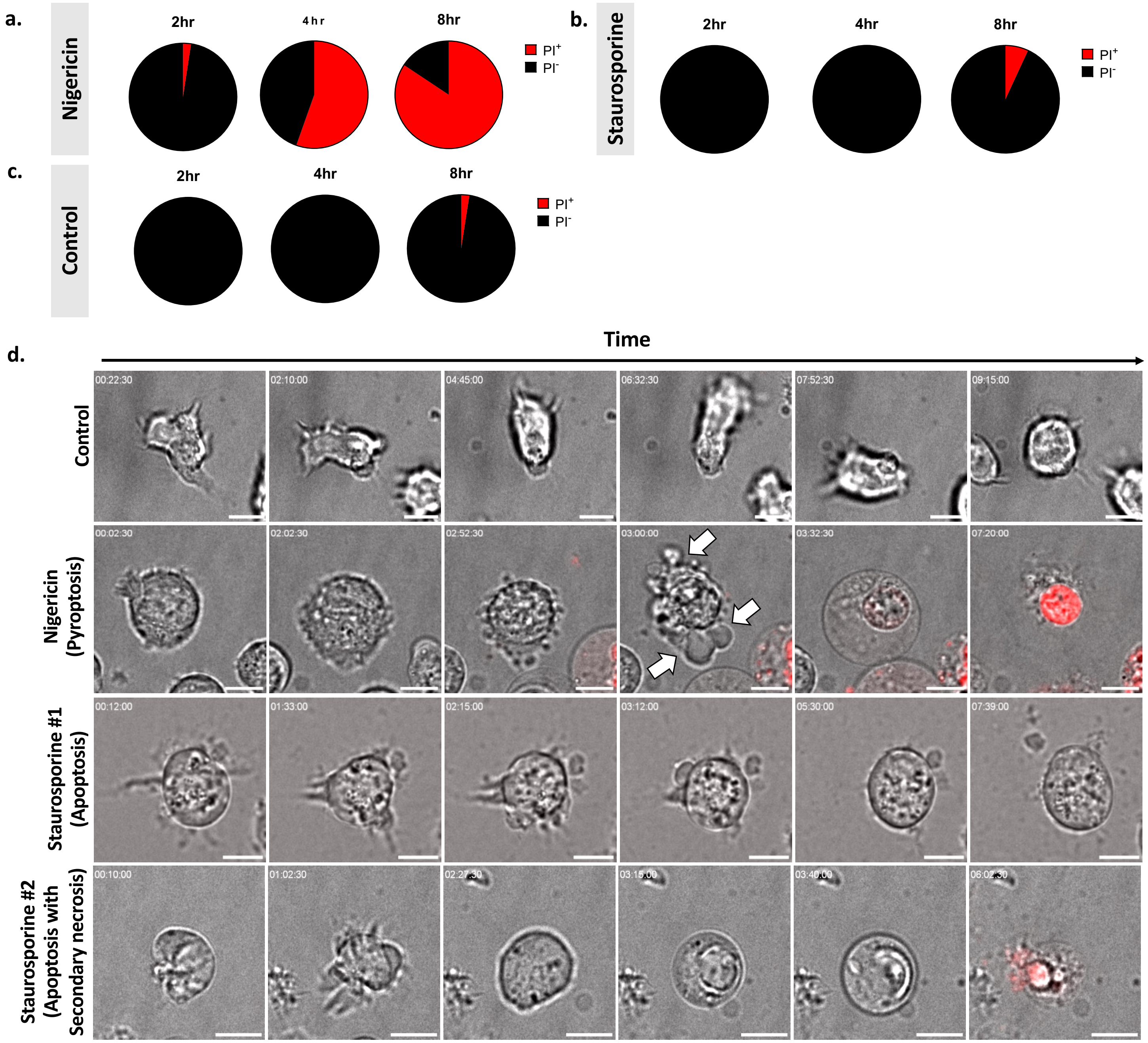


**Supplementary Figure 9| Time-lapse imaging of chemically-induced pyroptosis (nigericin) and apoptosis (staurosporine)**

**a-d.** JY cells were treated with nigericin (100µM), staurosporine (6.7µM) or solvent control in the presence of 6.6µg/mL propidium iodide (**PI**) and imaged using spinning disk confocal microscopy for 10hr. The proportion of nigericin-treated (**a.**), staurosporine-treated (**b.**) or solvent-treated (**c.**) cells that were PI-positive (*red*) and PI-negative (*black*) at each time point is represented. **d.** Representative examples drawn from n=3 biological experiments illustrating the morphology of nigericin-induced pyroptosis and staurosporine-induced apoptosis (with and without secondary necrosis) compared to controls. Snapshots are from **Supplementary Movie 3** (nigericin), **Supplementary Movie 6** (staurosporine), and **Supplementary Movie 7** (staurosporine with secondary necrosis). *White arrows* indicate pyroptotic bodies. Scale bars are 10µm.

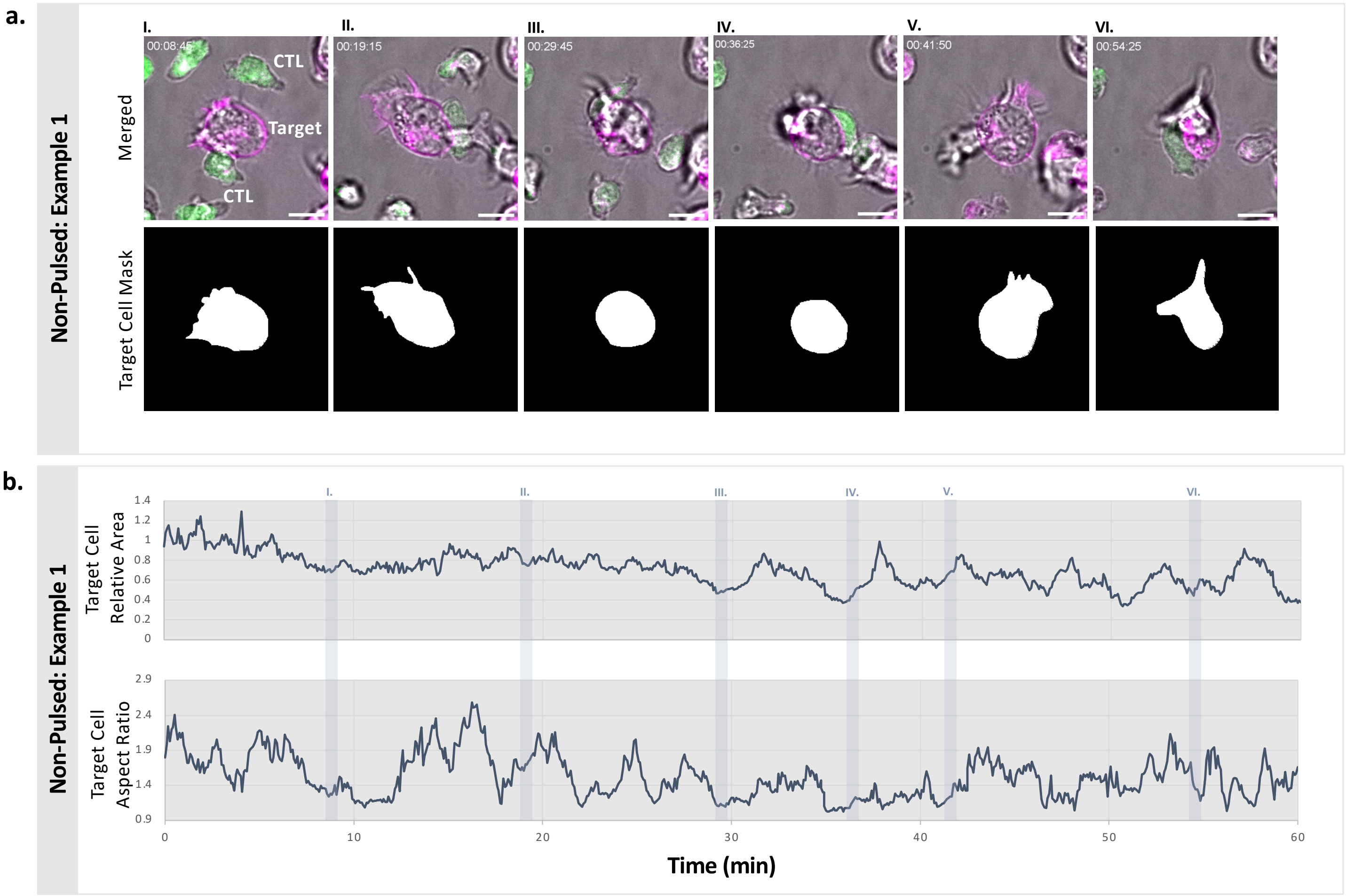


**Supplementary Figure 10| Morphological feature quantification of non-pulsed target cells during interactions with CTLs**

**a-b.** Non-pulsed JY target cells were loaded with CellMask DeepRed (*magenta*) and conjugated at an E:T ratio of 1:1 with antigen-specific human CTLs pre-loaded with Fluo-8-AM (*green*) and Tubulin Tracker Deep Red (*magenta*) for 60min and monitored by spinning disk confocal microscopy. **a.** One example representative of n=3 experiments (snapshots are from **Supplementary Movie 10**). To quantify the morphological features of cells, masks of the target cell were created for each frame using a semi-automated image analysis pipeline to permit the approximation of target cell cross-sectional area and aspect ratio over time. **b.** Data shown represent the non-pulsed target cell mask cross-sectional area (normalized to its size at the start of imaging) and aspect ratio over 60min. Scale bars are 10µm.

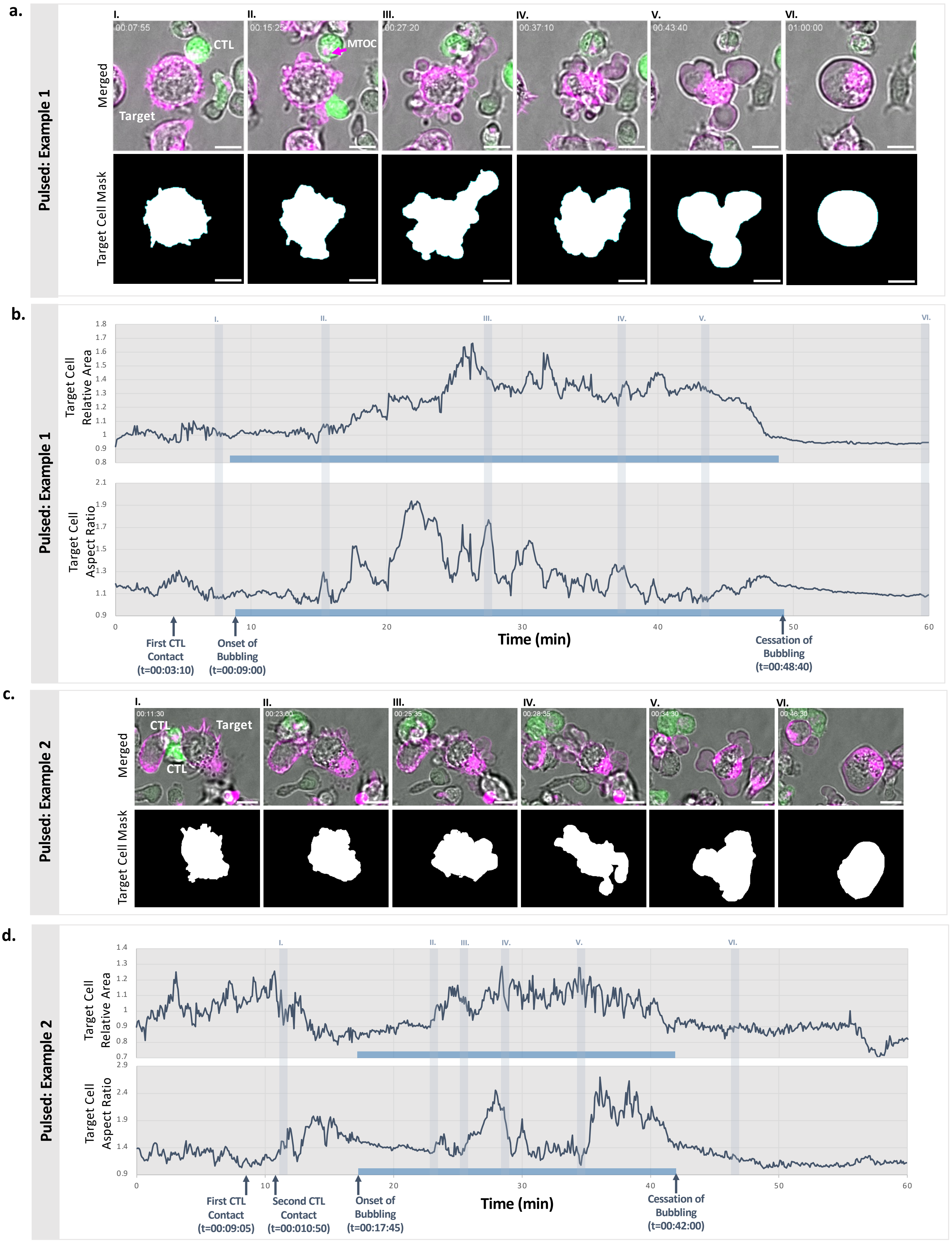
a.
Pulsed: Example 1
Merged
Target Cell Mask
I.
00:07:55
CTL
Target
II.
00:15:25
MTOC
III.
00:27:20
IV.
00:37:10
V.
00:43:40
VI.
01:00:00
b.
Pulsed: Example 1
Target Cell Relative Area
Target Cell Aspect Ratio
Time (min)
First CTL Contact (t=00:03:10)
Onset of Bubbling (t=00:09:00)
Cessation of Bubbling (t=00:48:40)
c.
Pulsed: Example 2
Merged
Target Cell Mask
I.
00:11:30
CTL
Target
CTL
II.
00:23:00
III.
00:25:35
IV.
00:28:35
V.
00:34:30
VI.
00:46:30
d.
Pulsed: Example 2
Target Cell Relative Area
Target Cell Aspect Ratio
Time (min)
First CTL Contact (t=00:09:05)
Second CTL Contact (t=00:010:50)
Onset of Bubbling (t=00:17:45)
Cessation of Bubbling (t=00:42:00)

**Supplementary Figure 11| Morphological feature quantification of antigen-pulsed target cells upon CTL attack**

**a-d.** JY target cells were loaded with CellMask DeepRed (*magenta*), pulsed with 10µM antigenic peptide, and conjugated at an E:T ratio of 1:1 with antigen-specific human CTLs pre-loaded with Fluo-8-AM (*green*) and Tubulin Tracker Deep Red (*magenta*) for 60min and monitored by spinning disk confocal microscopy (snapshots are from **Supplementary Movie 11** and **12**). **a,c.** Two examples representative of n=3 biological experiments. To quantify the rapidly changing morphology of the dying cell, masks of the target cell were created for each frame using a semi-automated image analysis pipeline to permit the approximation of target cell cross-sectional area and aspect ratio over time. **b,d.** Data shown represent the target cell mask cross-sectional area (normalized to its size at the start of imaging) and aspect ratio over 60min. The duration of the bubbling phase of cell death is indicated by the blue line. Time of first CTL contact is indicated with a blue arrow. Scale bars are 10µm.

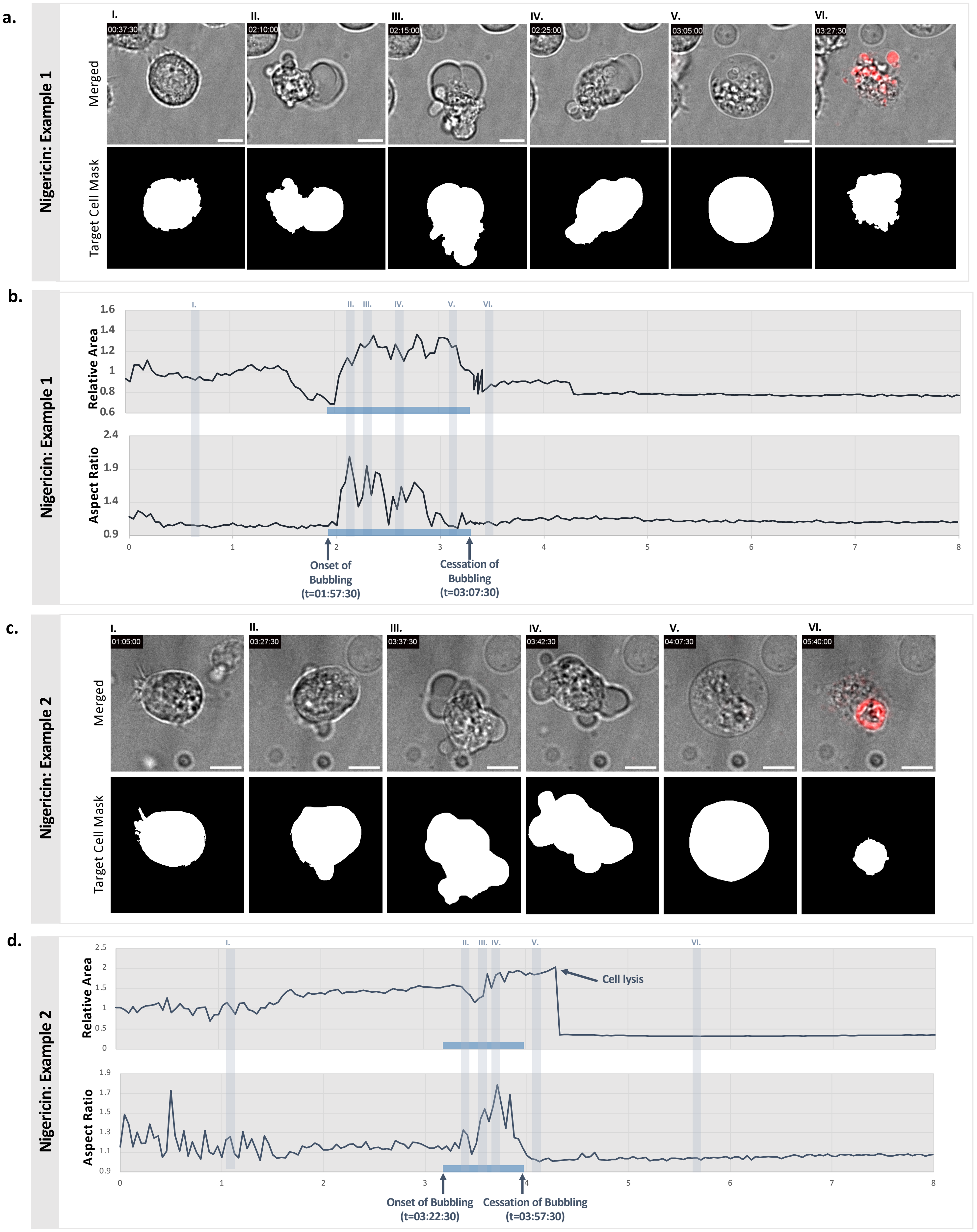

a.
Nigericin: Example 1
Merged
Target Cell Mask
I.
00:37:30
II.
02:10:00
III.
02:15:00
IV.
02:25:00
V.
03:05:00
VI.
03:27:30
b.
Nigericin: Example 1
Relative Area
Aspect Ratio
1.6
1.4
1.2
1
0.8
0.6
2.4
1.9
1.4
0.9
0
1
2
3
4
5
6
7
8
Onset of Bubbling (t=01:57:30)
Cessation of Bubbling (t=03:07:30)
c.
Nigericin: Example 2
Merged
Target Cell Mask
I.
01:05:00
II.
03:27:30
III.
03:37:30
IV.
03:42:30
V.
04:07:30
VI.
05:40:00
d.
Nigericin: Example 2
Relative Area
Aspect Ratio
2.5
2
1.5
1
0.5
0
1.9
1.7
1.5
1.3
1.1
0.9
Cell lysis
Onset of Bubbling (t=03:22:30)
Cessation of Bubbling (t=03:57:30)

**Supplementary Figure 12| Morphological feature quantification of nigericin-treated JY cells during pyroptosis**

**a-d.** JY cells were treated with nigericin (100μM) to induce pyroptosis and monitored by spinning disk confocal microscopy for 8hr in the presence of 6.6μg/mL propidium iodide (*red*). Snapshots are from **Supplementary Movie 4** and **5. a,c.** Examples representative of n=3 biological experiments. To quantify morphology of the pyroptotic cell, masks of the cell were created for each frame using a semi-automated image analysis pipeline to permit approximation of cell cross-sectional area and aspect ratio over time. **b,d.** Data shown represent cell mask cross-sectional area (normalized to the start of imaging) and aspect ratio over 8hr. Onset and cessation of bubbling phase indicated by blue bar. Scale bars are 10μm.

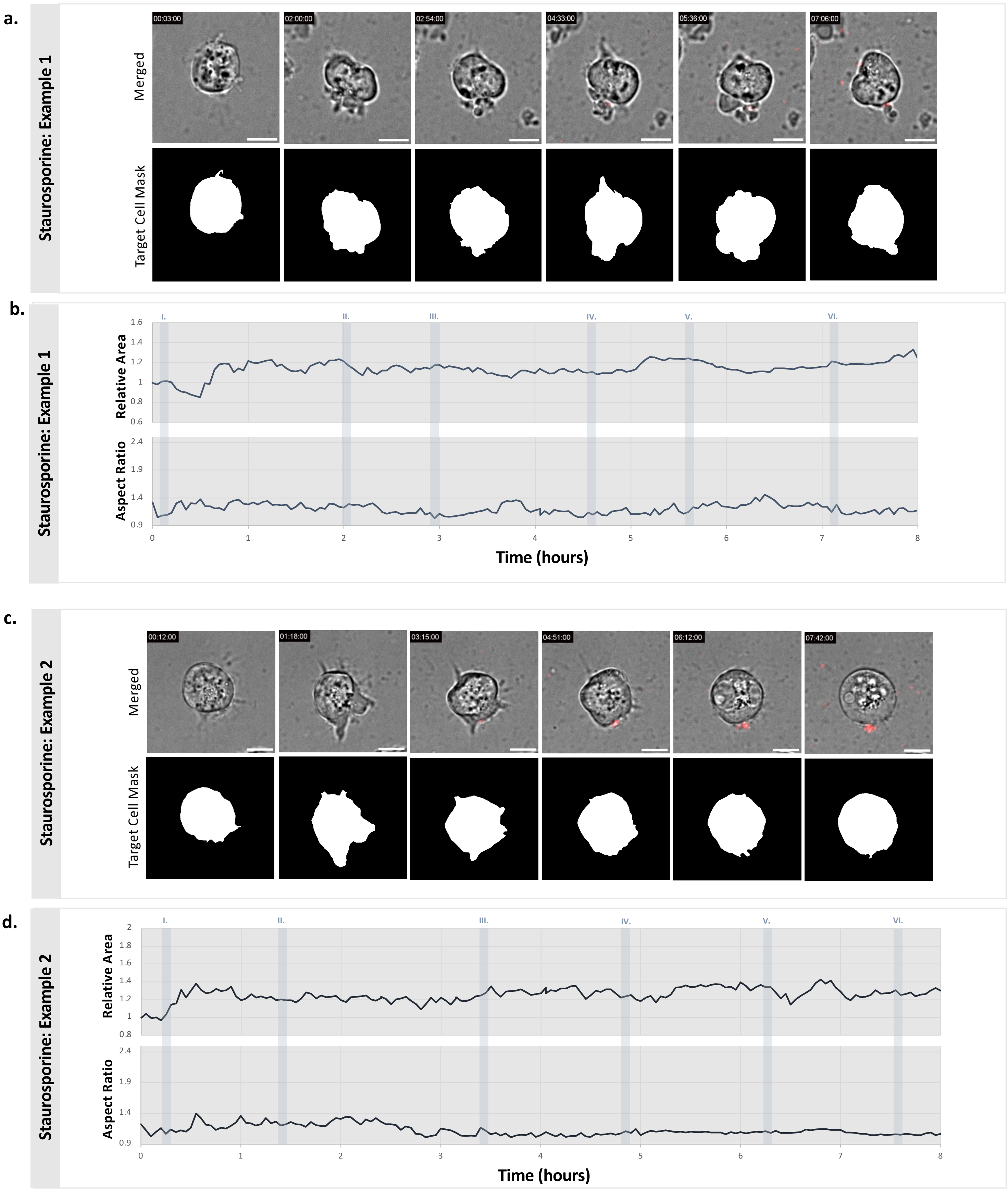


**Supplementary Figure 13| Morphological feature quantification of staurosporine-treated JY cells during apoptosis**

**a-d.** JY cells were treated with staurosporine (6.7µM) to induce apoptosis in the presence of 6.6µg/mL propdium iodide (*red*) and monitored by spinning disk confocal microscopy. Snapshots are from **Supplementary Movie 8** and **9. a,c.** Two examples of staurosporine-treated JY cells, representative of n=3 biological experiments. To quantify the morphology of the apoptotic cell, masks of the cell were created for each frame using a semi-automated image analysis pipeline to permit the approximation of cross-sectional area and aspect ratio over time. **b,d.** Data shown represent target cell mask cross-sectional area (normalized to the start of imaging) and aspect ratio over 8hr. Scale bars are 10µm.

a.

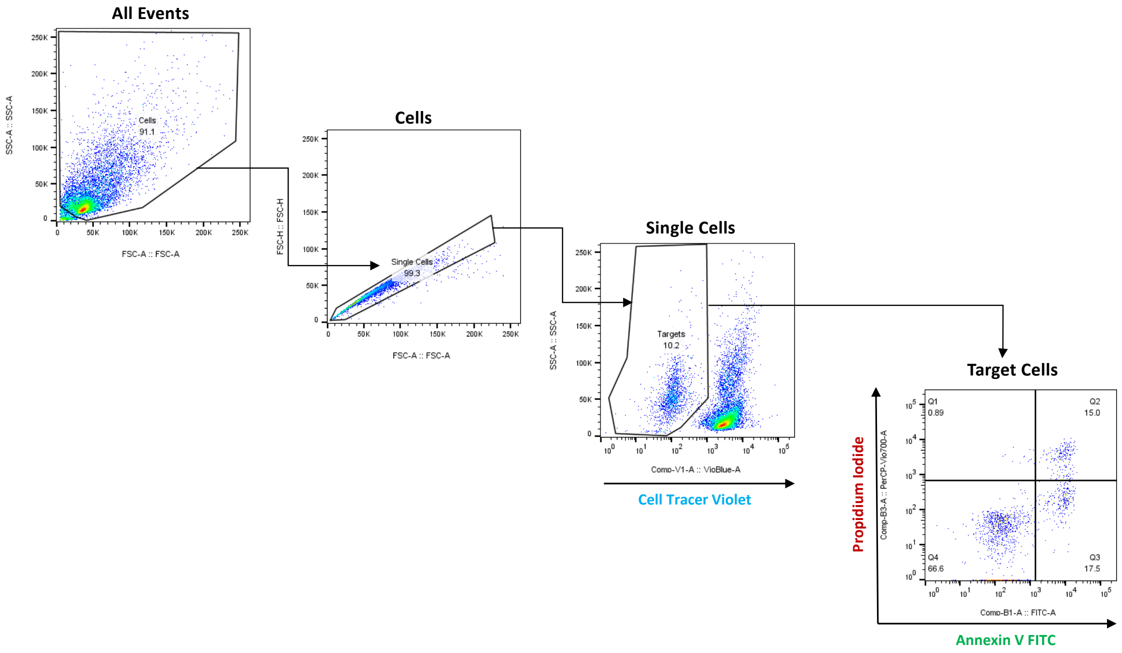


**Supplementary Figure 14| Gating strategy for Annexin V/PI assays**

**a.** Total cells were differentiated from debris on the basis of forward (**FSC-A**) and side-scatter (**SSC-A**). Single cells were selected by comparing FSC-A and FSC-H. Target cells were differentiated from Cell Tracer Violet-stained or Hoescht-stained CTLs based on SSC-A and exclusion of violet dye. Finally, the selected target cells were plotted on a scatterplot with Annexin V FITC on the x-axis and propidium iodide staining on the *y*-axis. The percentage of cells falling in each quadrant was reported.

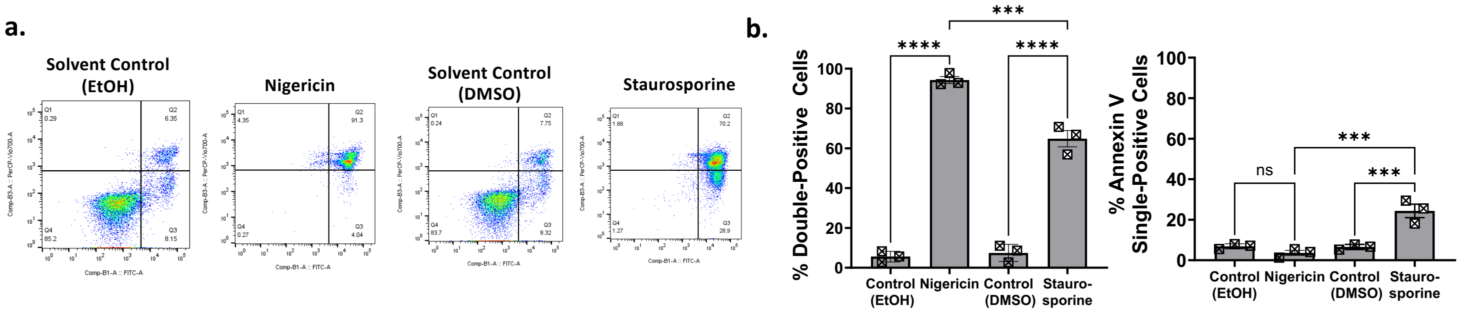


**Supplementary Figure 15| Molecular features of chemically-induced pyroptosis and apoptosis**

**a,b.** JY cells were treated with nigericin (100µM), staurosporine (6.7 µM), or respective solvent controls overnight (18h), subjected to Annexin V/PI staining and analyzed by flow cytometry. **a.** Representative dotplots from n=3 biological experiments. **b.** Data shown represent the percentage of cells in the quadrants of interest (Annexin V/PI double positive, Annexin V single-positive) at the experimental endpoint. Individual datapoints represent the results of a single experiment (average of two technical replicates), with a total of n=3 independent experiments. Ordinary one-way ANOVA with Tukey's test for multiple comparisons was utilized to assess statistical significance. Bars represent mean +/- SEM. *** $p$<0.001, **** $p$<0.0001, ns = non-significant.

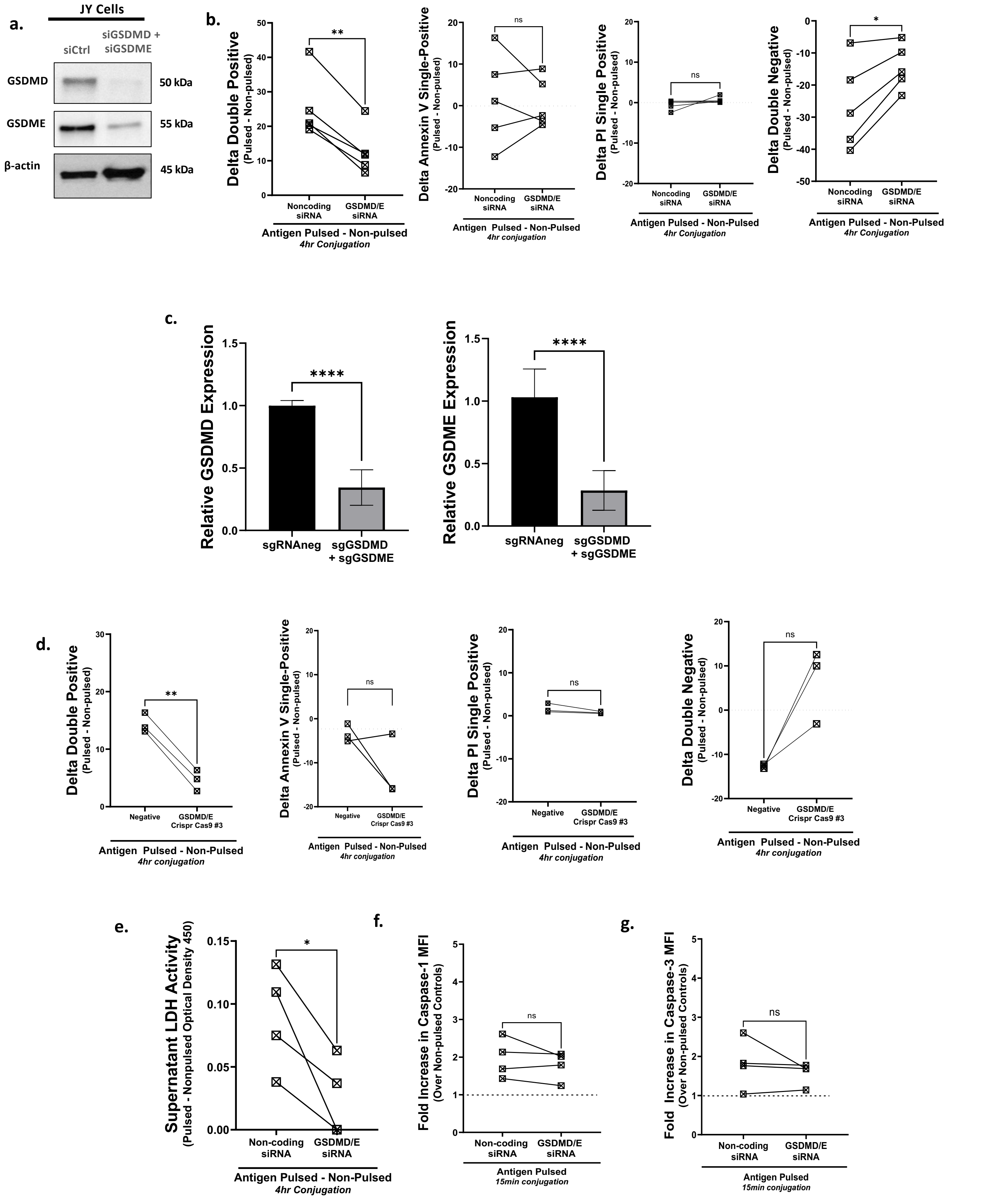
a.
JY Cells
siCtrl
siGSDMD + siGSDME
GSDMD
50 kDa
GSDME
55 kDa
β-actin
45 kDa
b.
Delta Double Positive (Pulsed - Non-pulsed)
**
Noncoding siRNA
GSDMD/E siRNA
Antigen Pulsed - Non-pulsed
4hr Conjugation
Delta Annexin V Single-Positive (Pulsed - Non-pulsed)
ns
Antigen Pulsed - Non-Pulsed
4hr conjugation
Delta PI Single Positive (Pulsed - Non-pulsed)
Delta Double Negative (Pulsed - Non-pulsed)
*
c.
Relative GSDMD Expression
Relative GSDME Expression
****
sgRNAneg
sgGSDMD + sgGSDME
d.
Negative
GSDMD/E Crispr Cas9 #3
e.
Supernatant LDH Activity (Pulsed - Nonpulsed Optical Density 450)
Non-coding siRNA
f.
Fold Increase in Caspase-1 MFI (Over Non-pulsed Controls)
Antigen Pulsed
15min conjugation
g.
Fold Increase in Caspase-3 MFI (Over Non-pulsed Controls)

**Supplementary Figure 16| Gasdermin knockdown and knock-out reduces CTL-mediated target cell death**

a. JY cells were transfected with non-targeting siRNA (**siCtrl**) or siRNAs targeting *GSDMD* and *GSDME* (**siGSDMD+siGSDME**), harvested, and subject to immunoblot for GSDMD and GSDME with β-actin as a loading control. **B.** JY cells were transfected with non-targeting siRNA or siRNAs targeting GSDMD and GSDME, before being pulsed with antigen, conjugated with antigen-specific CTLs for 4hrs, stained with Annexin V/PI and assessed by flow cytometry. Data shown represents the difference in the percentage of Annexin V/PI double-positive cells, Annexin V single-positive cells, PI single-positive cells and double negative cells in the pulsed versus non-pulsed conditions after conjugation with CTLs. Paired *t*-tests were utilized to assess statistical significance. Each datapoint represents the average of two technical replicates from an individual experiment, with n=5 independent experiments represented on the graph. **c.** GSDMD/E double knock-out cells were generated by electroporation of JY cells with sgRNA GSDMD + sgRNA GSDME and Cas9. Control cells were transfected with non-targeting sgRNA and Cas9. Expression of GSDMD and GSDME transcript was assessed in both populations by bulk qPCR. Data shown represent the mean +/- SD for six technical replicates of a representative transfection experiment. Inpaired *t*-tests were utilized to assess statistical significance. d. GSDMD/E double knock-out JY cells were conjugated with CTLs for 4hr and target cell death assessed by Annexin V/PI assay. Data shown represents the difference in the percentage of Annexin V/PI double-positive cells, Annexin V single-positive cells, PI single-positive cells and double negative cells in the pulsed versus non-pulsed conditions after conjugation with CTLs. Paired *t*-tests were utilized to assess statistical significance. Each datapoint represents the average of two technical replicates from an individual experiment, with n=3 independent experiments represented on the graph. e. Supernatants were collected after 4hr conjugation and subjected to LDH assay. Data shown represent the difference in supernatant LDH activity in the pulsed versus non-pulsed conditions after 4hr conjugation with CTLs. Each datapoint represents the average of two technical replicates from an individual experiment, with a total of n=4 independent experiments represented on the graph. Paired *t*-tests were utilized to assess statistical significance. **f,g.** To assess whether GSDMD/E knockdown affected upstream activity of their respective caspases, siRNA-transfected JY cells were conjugated for 15min with CTLs, subjected to activity-dependent fluorescent caspase-1 and -3 assays and assessed by flow cytometry. Data represent fold increase in MFI for each caspase probe, normalized to respective non-pulsed controls (shown by dotted line). Each datapoint represents the average of two technical replicates from an individual experiment, with a total of n=4 independent experiments represented on the graph. Paired *t*-tests were utilized to assess statistical significance. *$p$<0.05, ** $p$<0.01, ns = not significant.

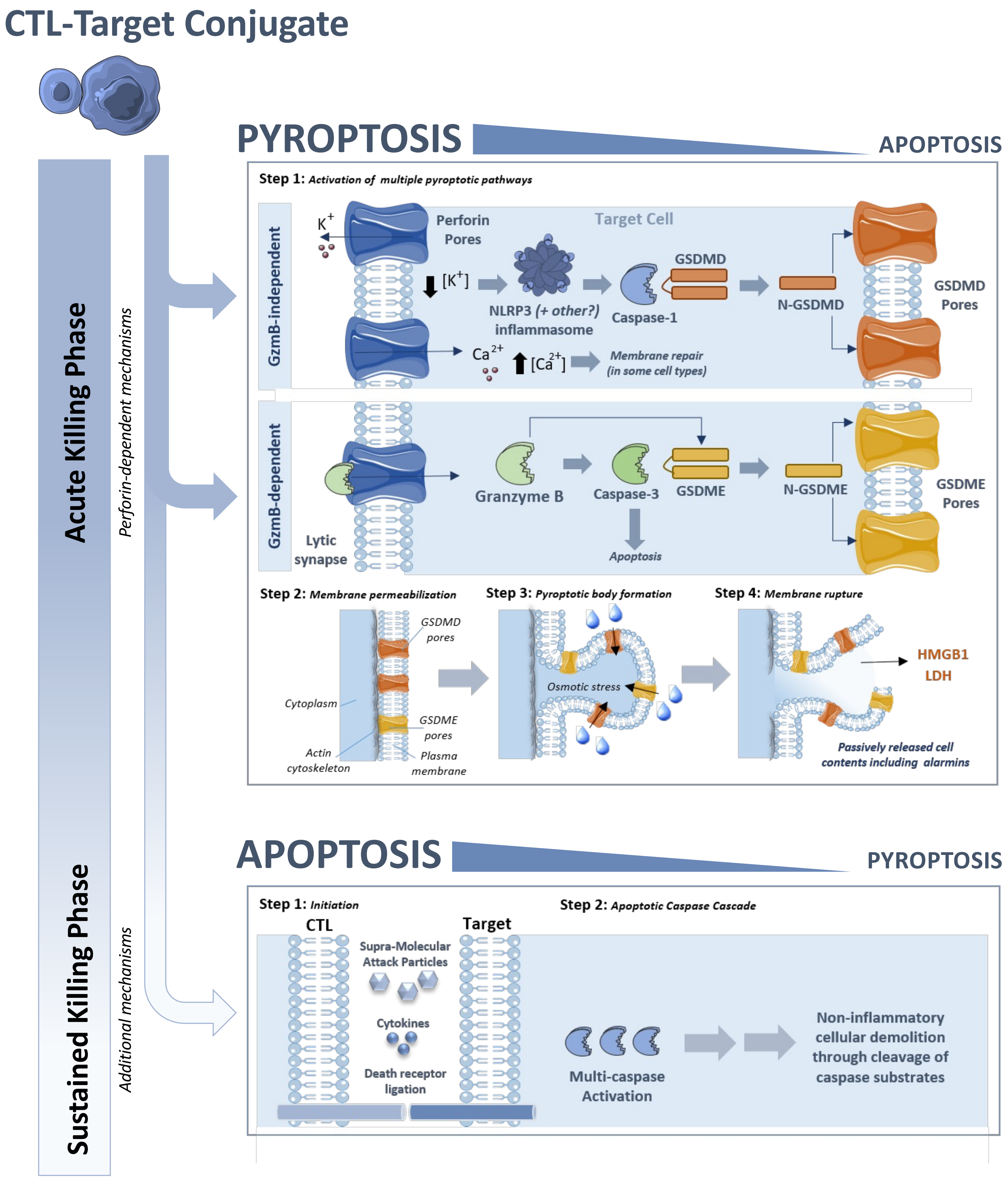

CTL-Target Conjugate
Acute Killing Phase
Perforin-dependent mechanisms
Sustained Killing Phase
Additional mechanisms
PYROPTOSIS
APOPTOSIS
Step 1: Activation of multiple pyroptotic pathways
GzmB-independent
$K^+$
Perforin Pores
Target Cell
$[K^+]$
NLRP3 (+ other?) inflammasome
Caspase-1
GSDMD
N-GSDMD
GSDMD Pores
$Ca^{2+}$
$[Ca^{2+}]$
Membrane repair (in some cell types)
GzmB-dependent
Lytic synapse
Granzyme B
Caspase-3
GSDME
N-GSDME
GSDME Pores
Apoptosis
Step 2: Membrane permeabilization
GSDMD pores
Cytoplasm
GSDME pores
Actin cytoskeleton
Plasma membrane
Step 3: Pyroptotic body formation
Osmotic stress
Step 4: Membrane rupture
HMGB1
LDH
Passively released cell contents including alarmins
APOPTOSIS
PYROPTOSIS
Step 1: Initiation
CTL
Target
Supra-Molecular Attack Particles
Cytokines
Death receptor ligation
Step 2: Apoptotic Caspase Cascade
Multi-caspase Activation
Non-inflammatory cellular demolition through cleavage of caspase substrates

**Supplementary Figure 17| Schematic depicting multi-pronged target cell death pathways activated upon different phases of CTL attack**

*Acute killing phase (top):* upon initial CTL attack, intracellular perforin stores are released into the lytic synapse, whereupon they oligomerize to form pores in the target cell membrane. *GzmB-independent pyroptosis pathway*: perforin pores are permeable to $K^+$ ions, mediating a swift efflux of $K^+$ out of the cell and leading to a drop in intracellular [$K^+$]. Potassium-sensitive inflammasome sensor proteins (such as NLRP3) are activated by the decrease in intracellular [$K^+$], providing a platform for proximity-induced auto-proteolysis of caspase-1. Activated caspase-1 in turn cleaves and activates its major substrate, the pyroptotic executioner GSDMD. *GzmB-dependent pyroptosis pathway*: as previously reported, granzyme B enters target cells through perforin pores, whereupon it may either cleave GSDME directly or cleave and activate caspase-3, which in turn activates GSDME[52-53]. Activated GSDMD and GSDME oligomerize to form their respective transmembrane pores, which permeabilize the target cell plasma membrane, leading to osmotic stress, influx of water, and formation of pyroptotic bodies. Rupture of these pyroptotic bodies leads to release of intracellular contents. *Sustained killing phase (bottom):* under conditions of prolonged interactions with targets, pyroptosis gives way to apoptotic killing mechanisms, which may be initiated by one or more apoptotic signals (e.g. uptake of supra-molecular attack particles, cytokine release and/or death receptor ligation), that ultimately lead to activation of a multi-caspase cascade and systemic intracellular demolition through regulated intracellular proteolysis. While pyroptotic cell death mechanisms dominates at early time points, apoptosis is favoured in the sustained killing phase.
